# Surface-guided computing to quantify dynamic interactions between cell morphology and molecular signals in 3D

**DOI:** 10.1101/2023.04.12.536640

**Authors:** Felix Y. Zhou, Virangika K. Wimalasena, Qiongjing Zou, Andrew Weems, Gabriel M. Gihana, Edward Jenkins, Bingying Chen, Bo-Jui Chang, Meghan Driscoll, Andrew J. Ewald, Gaudenz Danuser

## Abstract

Many signalling circuits are governed by the spatiotemporal organization of membrane-associated molecules. Growing evidence suggests that the mesoscale cell surface geometry is central in modulating this interplay. However, defining the causal hierarchy between geometric and molecular factors that control signals remains challenging. Nonlinearity and redundancy among the components prevent direct experimental perturbation, with shape being the most difficult to independently control. Towards the goal of inferring causality from observational data, we developed u-Unwrap3D as a resource to map arbitrarily complex 3D cell surfaces to diverse representations, each designed to interrogate a different aspect of the dynamic interaction between cell surface geometry and molecular cues. Using u-Unwrap3D, we discover a retrograde protrusion flow on natural killer cells associated with immunological synapse formation with cancer; establish a causal association of K14+ cells with breast tumor organoid invasion; measure the speed of ruffles; and quantify bleb-mediated assembly of septin polymers at the membrane.

## Introduction

Advances in 3D high-resolution live-cell microscopy and biosensor design enable integrative studies of the dynamic interplay between cell morphology and cellular signals *in vitro* and *in vivo*. By reshaping the plasma membrane into diverse morphologies, cells sense, respond to and remodel their local environment^1-6^. Many cell types adopt characteristic shapes tailored to their function^7-11^. Cell morphology has thus long been recognised and used as a proxy of cell state and a marker of differentiation^9,11-13^. Mechanically, the plasma membrane integrates internal and external forces, shaping cell fate through mechanotransductive proteins and inducing changes in the cytoskeleton^14,15^. Structurally, the plasma membrane is a platform for catalysing chemical reactions^16-20^ and a spatiotemporal organiser of signalling activity by creating binding sites, and local confinements to concentrate molecules through scaffolds, diffusion traps, and phase separation^16,21-23^. These reactions occur locally at the nanometer or micrometer length scale or in global bursts that span the entire cell^18,24^. Understanding the salient biophysical processes that govern the formation and persistence of these subcellular signalling domains and establishing that these domains are not merely products of activity but can causally drive biochemical signal transduction, remains enigmatic. One of the first hurdles to enable such studies is the sampling and representation of dynamic and wildly variable 3D cell shape and molecular activities at the appropriate spatial and temporal resolution.

Cell surfaces are usually extracted from binary image segmentations and stored as a mesh, a data structure described by a vertex list of the Cartesian 3D coordinates and a second list specifying how individual vertices are locally connected into triangle faces (Fig. 1A). Meshes parameterize the shape but are not unique representations. The same shape can be remeshed to have a different number of vertices and faces. Consequently, establishing full correspondence between two 3D surface meshes, e.g. over time, is nontrivial and an active area of research in computer graphics^25-28^. Method developments in this field largely focus on well-defined shapes with fixed topology such as human and animal pose^29,30^ or hands^31,32^. Generally, these methods track by pairwise matching of meshes. To match, methods primarily compute a unique signature per vertex to establish vertex correspondences by minimizing a complex loss metric^30,33^. However, this approach is inherently sensitive to mesh quality, the uniqueness of the signature, the disparity in pose, and optimizer convergence. Additionally, matching errors compound, making it difficult to track surfaces over many timepoints, which is a common task in live cell microscopy. A further technical challenge that arises when adapting computer graphics techniques to cell biology is the non-convexity, irregularity and intrinsically high curvature of surface protrusions found with most cell shapes. Critically, protrusions do not have well-defined prototypical shapes. Not only are their shapes transient and remodeling topologically over time, but cells can also dynamically change the spatial distribution of protrusions. This lack of preserved surface features poses significant ambiguity in matching. For similar reasons, registering 3D volumes spatiotemporally before mesh extraction, as done in neuroscience for brain mapping^34-37^, is also suboptimal. Whilst less sensitive to mesh quality and topological errors, these methods can cope only with small deformations. In contrast to these applications of surface matching, cell protrusions decorate a wide spectrum of irregular and branched cortical shapes, and are small relative to the cell volume. Particularly, long or thin cell surface structures, such as lamellipodia and filopodia, suffer voxel undersampling, limiting volumetric-based registration to surfaces with largely globular features such as blebs^38^. Alternatively, individual surface protrusions^39^ can be segmented and reduced to particle representations amenable to 3D object tracking^40,41^. This approach is susceptible to the variable quality of the segmented motifs and non-convexities of the surface but can be effective for quantifying protrusion movement when motion is small and simple between timepoints^39,40^. However, because the entire surface is not tracked, we cannot relate bulk protrusion measurements to local molecular activity, where at each timepoint, the segmented protrusion surface has a different number of vertices and face connectivity. Finally, surface correspondence can be determined by remapping shapes to a standard geometry. In macroscopic imaging applications, mapping 3D surfaces to the unit sphere has enabled registration of complex geometrical features such as brain folds directly on the 3D sphere or in derivative 2D UV unwrapped images^27,42-44^. In developmental biology, parameterization to unwrap the 3D surface into 2D has revolutionized the profiling of tissue flow. Methods developed include Mercator-based projections of the local surface of large organismal volumes^45,46^; spline-fitting to adapt cylindrical coordinates to axisymmetric shapes such as drosophila^47^ and mouse^48^ embryos and tubular intestine^49^ structures; and more advanced conformal^49,50^ and equiareal^51^ maps from differential geometry to map the same surfaces and those of more general, asymmetric shapes like the zebrafish heart^47,51^. Such ideas have also been applied to single cells. Primarily, spherical mapping is used to reparameterize cells into a canonical coordinate system, where spherical harmonics can then be used for generative modelling and classifying shape variation (both within a collection and over time), and describing intracellular organization^13,52-56^. A major limitation of these techniques is that the surfaces must have closed topology, be manifold and have no holes (genus-0 surfaces) - a requirement often violated in live-cell microscopy due to segmentation inaccuracies from resolution limits and signal-to-noise ratio. Cells are also not rigid bodies. Topological changes are integral in biological processes, like migration^57^ and macropinocytosis^58^. Consequently, all biological applications to date have been limited to smooth shapes or a star-convex approximation of the shape. A second limitation is metric distortion. The longer a protrusion, or the more branched a cell, the worse the induced distortion, rendering any distance or area-based analyses inaccurate if performed purely on the 2D unwrapped images. Whilst computer graphics techniques like orbifold maps^59^, surface cutting^60^, or mapping into 2D without fixing the domain boundary^61^ can alleviate this problem, they generate multi-piece maps or irregular 2D domains, both unsuited for temporal analysis.

**Figure 1.**
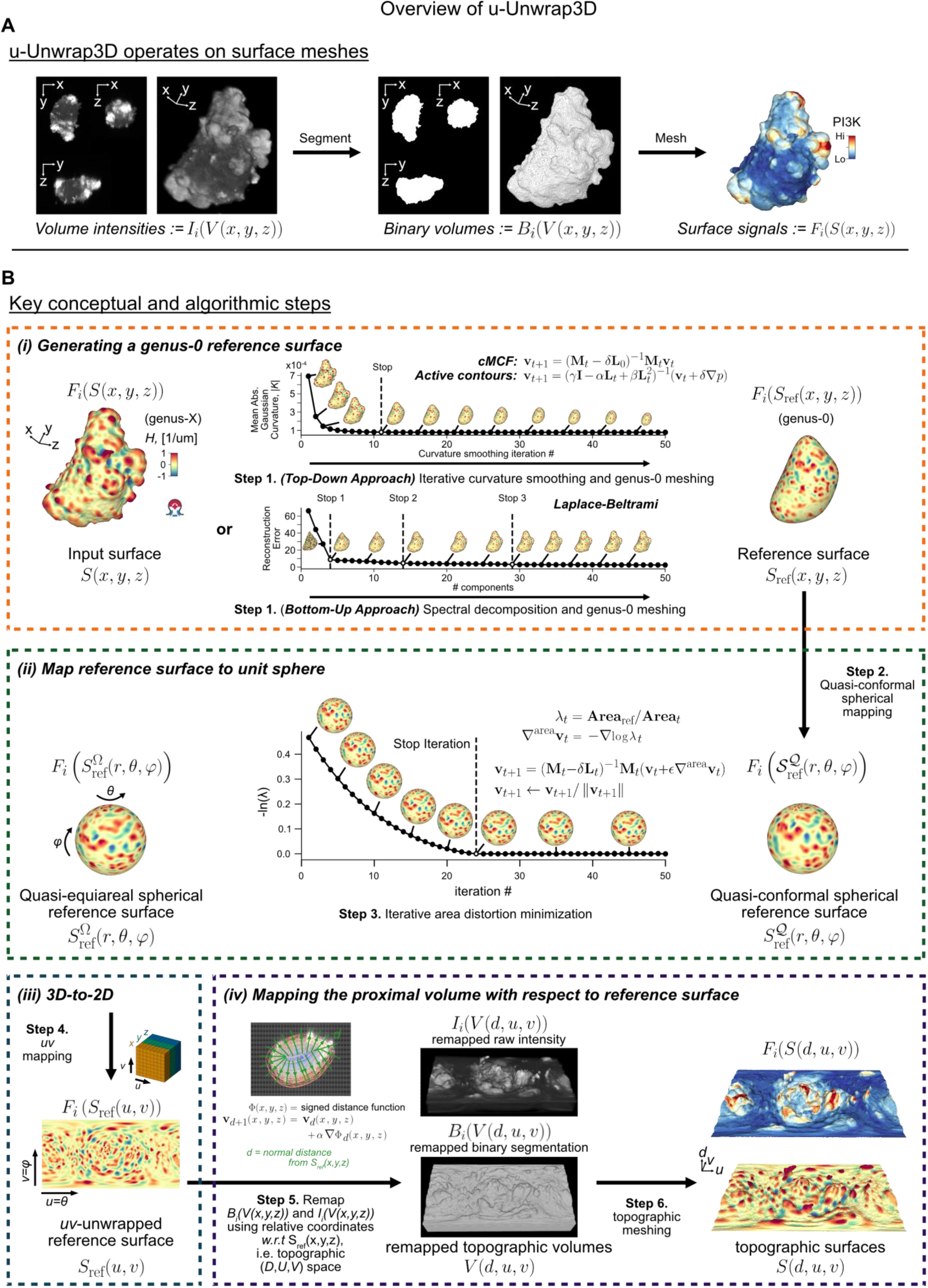
Overview of the surface-guided computing framework u-Unwrap3D. **A)** u-Unwrap3D operates on cell surface meshes such as those extracted from the binary segmentation of an input 3D volume image, *I*_*i*_, and associated scalar measurements, *F*_*i*_(*S*(*x*, y, *z*)) into different surface and volumetric representations for performing computation. **B)** Overview of the primary steps to map an input genus-X Cartesian 3D surface, *S*(*x*, y, *z*), via a genus-0 reference surface, *S*_ref_(*x*, y, *z*), into any of three additional representations; topographic 3D surface, *S*(*d, u, v*), 3D sphere, *S*_ref_(*r, θ, φ*) and 2D plane, *S*_ref_(*u, v*). **v** denotes mesh vertex coordinates, **M** is the mesh mass matrix, **L** is the mesh Laplacian matrix, *ϵ* the step size of area-distortion relaxation, *α* the step size (in pixels) of the propagation distance, *d* the topographic depth (in pixels) and *t* the iteration number. The reference surface can be flexibly defined: user-provided, mean shape of a timeseries, or computed from *S*(*x*, y, *z*) . For the latter, u-Unwrap3D implements curvature-minimizing geometric flows for top-down and spectral decomposition of Laplace-Beltrami operator for bottom-up construction as illustrated with automatic inference of iteration number for reference shape choice (Methods). In all cases, u-Unwrap3D implements shrink-wrapping (Methods) to guarantee the reference mesh is genus-0 and of sufficient mesh quality required for spherical parameterization. u-Unwrap3D can also map directly input genus-0 surfaces or genus-X surfaces after being shrink-wrapped to be genus-0 using u-Unwrap3D, skipping the definition of a reference shape. In the panels, *S*(⋅), *V*(⋅) denote surface and volume geometries, respectively, in either Cartesian 3D, topographic 3D, or 3D spherical coordinates; *F*_*i*_(*S*(⋅)) and *I*_*i*_(*V*(⋅)) denote surface or volumetric signals as a function of a particular surface or volume geometry. *B*_*i*_(*V*(⋅)) is a binary segmentation in the bracket specified volumetric coordinates. *H* and *K* denote mean and Gaussian curvatures respectively.

Based on the review of these existing studies, we recognized that there may not be a single best representation to solve the correspondence problem and to perform diverse quantitative analyses. Different 3D representations each offer unique advantages: Cartesian 3D for geometric measurement, 3D sphere for alignment and registration, and 2D UV planes for tracking. Inspired by this observation, we developed u-Unwrap3D, a software package for surface-guided computing. Developed on differential geometry theory and real-world validated computer graphics algorithms, u-Unwrap3D provides comprehensive geometry processing routines to robustly remap arbitrarily complex 3D cell surface morphology with membrane-associated signals to a diverse selection of equivalent surface and volumetric representations, to facilitate optimized measurement of surface geometry and associated molecular entities, and to spatiotemporally register and track surface features. We demonstrate the power of u-Unwrap3D to flexibly design pipelines for analyzing diverse challenging temporal problems: (i) measuring the information flow of mean curvature, to establish a retrograde protrusion flow on natural killer cells facilitates immunological synapse formation with cancer cells; (ii) measuring the correlation between information flow of tumor organoid strand length and K14 signal, to establish the strength of causal association between K14+ cells and invasion in breast organoids; (iii) measuring the travel speed of actin-enriched surface ruffles; (iv) unsupervised segmentation of surface motifs; and (v) quantifying the extent of septin polymer recruitment to cell blebbing events. u-Unwrap3D is implemented in Python 3, is freely available and installable from https://github.com/DanuserLab/u-unwrap3D or the Python Package Index, PyPI.

## Results

### u-Unwrap3D for surface-guided computing

u-Unwrap3D is a toolkit for mapping arbitrarily complex Cartesian 3D surface *S*(*x, y, z*) to a series of equivalent surface representations including the standard reference geometries of a unit 3D sphere and 2D UV image (Movie S1). Here, ‘complex’ refers to non-convex and branched shapes with narrow necks and a surface decorated by high-curvature protrusions. We exclude neuron-like cells with intricate branching topologies, which share greater similarities with vasculature networks, and more appropriately analyzed by fitting spline-based centerlines. What makes u-Unwrap3D unique are the specific features introduced to facilitate spatiotemporal analysis of protrusion dynamics on complex cell surfaces. These include:

1. Construction of an intermediate protrusion-of-interest free reference surface *S*_ref_(*x, y, z*) from *S*(*x, y, z*) to enable decoupled analysis of protrusion and cortical shape dynamics (Fig. S1).
2. Guaranteed genus-0 shrink-wrapping of any complex genus-X *S*_ref_(*x, y, z*) or *S*(*x, y, z*) to enable spherical parameterization of either (Fig. S2).
3. Robust iterative optimization schemes and loss functions for conformal, isometric and equiareal sphere parameterization of complex genus-0 surfaces (Fig. S3).
4. Optimization of unwrapping axis and (*u, v*) aspect ratio to represent surface features-of-interest in a (*u, v*) parameterization with minimal metric distortion that facilitates quantitative analysis (Fig. S4).
5. Robust schemes to extend the (*u, v*) parameterization to remap a proximal volume around *S*_ref_(*x, y, z*) topographically, and even to sample the entire inner cell volume.
6. A topographic *S*(*d, u, v*) representation that reconstructs the 3D geometry of *S*(*x, y, z*) protrusions relative to *S*_ref_(*x, y, z*). *S*_ref_(*x, y, z*) is integrated out and is a 2D plane in this representation (Fig. S5).

The mappings between representations are ‘bidirectional’. A bijective map is mathematically only possible between two representations with identical topology. Between other representations, u-Unwrap3D constructs an as-bijective-as-possible mapping by surface matching, finding the minimum normal distance between surfaces through axis-aligned bounding boxes and vector cross-products (Methods). Mappings also minimize optimally metric distortion when applicable - conformal (preservation of aspect ratio), equiareal (preservation of surface area fraction) and isometric (balancing conformal and equiareal) errors^62^ (Fig. S3).

Fig. 1B and Movie S1 illustrate the six primary algorithmic steps and mappings to construct u-Unwrap3D representations. Together they implement four conceptual objectives: (i) generation of a protrusion-less genus-0 reference surface, *S*_ref_(*x, y, z*) from *S*(*x, y, z*); (ii) spherical parameterization of *S*_ref_(*x, y, z*); (iii) 2D UV unwrapping of the 3D spherical parameterization and (iv) remapping of the proximal volume surrounding the reference by propagation of the (*u, v*) -parameterized *S*_ref_(*x, y, z*) to reconstruct a 3D topographic representation of *S*(*x, y, z*) highlighting all landmarks and regions protruding normal to the reference shape. Throughout we denote *S*(⋅) and *V*(⋅) for surface and volume geometries, respectively, in the coordinate system specified within brackets. The variables *F*_*i*_(*S*(·)) and *I*_*i*_(*V*(·)) denote surface- and volume-associated signals of interest parameterized by the coordinate system in brackets. They may describe scalar measurements that are geometrical, (like mean curvature, *H*), or molecular (like molecular concentrations or activities), or categorical integer labels (like segmented surface protrusions). We use *B*_*i*_(*V*(⋅)) to denote binary volume segmentations. Notably, these variables can be computed in one representation and, through the geometrical mapping, be transferred to other representations for further analysis. Movie S1 and Fig. 1B illustrates this using the mean curvature of *S*(*x, y, z*) and the voxel intensity at *V*(*x, y, z*) as an exemplar of *F*_*i*_(*S*(·)) and *I*_*i*_(*V*(·)), respectively.

We describe the six algorithmic steps below for a single shape. Crucially, u-Unwrap3D presents multiple implementations and options for each step to offer avenues for tradeoff between speed, accuracy, and general applicability.

**Step 1** determines a genus-0 reference surface *S*_ref_(*x, y, z*) without holes or ‘handles’, representing *S*(*x, y, z*) minus the protrusive features-of-interest. ‘Handles’ are holes made by a loop, such as the handle of a teacup that, unlike ‘holes’, does not involve missing/incomplete surface patches in a mesh. u-Unwrap3D can find a suitable *S*_ref_(*x, y, z*) automatically through either a top-down or bottom-up approach (Fig. 1B, orange box). Going top-down, curvature-minimizing flows are used to iteratively smooth out high-curvature surface features. The flow may be free-form, implemented using conformalized mean curvature flow (cMCF)^63^, with generated shapes unconstrained to be internal to the volume of *S*(*x, y, z*). Alternatively, the flow can be guided, using active contours to propagate *S*(*x, y, z*) along the gradient of a distance transform, with step size weighted by mean curvature (Fig. S1, Methods). Going bottom-up, spectral decomposition of the Laplace-Beltrami operator^64^ is used to iteratively reconstruct *S*(*x, y, z*) using an increasing number of low-frequency modes (Methods). cMCF is fastest, and generates the smoothest shapes, free of neck-pinching singularities that may complicate spherical parameterization later. However, cMCF deformation shifts the centroid causing shape misalignment with the input surface, particularly for branched cell shapes (Fig. S1). For these shapes, guided deformation or spectral decomposition should be used. Notably, designing the guided flow to preferentially erode positive curvature generates an optimum surface for generating intuitive topographic maps (Fig. 1B, Methods). Meanwhile, spectral decomposition provides the theoretically tightest *S*_ref_(*x, y, z*). However, in practice, a large number of components may be required whilst the eigendecomposition calculation is slow and memory-intensive. We therefore reserve spectral decomposition for the most complex, branched shapes. In the absence of *a priori* cell cortex markers, we monitor the mean absolute Gaussian curvature *K* for curvature-minimizing flows, and the reconstruction error measured by chamfer distance (CD) to determine the stopping iteration (Methods). The Gaussian curvature *K* is a scale-invariant measure of local curvature^65^. *K* is thus well-suited as a termination criterion: A threshold based on the rate of decrease in absolute *K* is sufficient for simple shapes (Fig. 1B, Methods). However, *K* is sensitive to the mesh quality and varies for different motifs. The discrete Gaussian curvature we use measures the angle deficit at each vertex (Methods). Most generally, we detect changepoints in *K* or the reconstruction error to find a set of candidate *S*_ref_(*x, y, z*) for cMCF and spectral decomposition (stops in Fig. 1B, Methods). The number of found candidates reflects a hierarchy of distinct shapes within *S*(*x, y, z*). We take the first and last changepoints for cMCF and spectral decomposition, respectively, to generate the closest possible *S*_ref_(*x, y, z*) to *S*(*x, y, z*). For guided flows, by construction, the optimal *S*_ref_(*x, y, z*) corresponds to the iteration with minimum absolute *K* value. All approaches preserve topology. Any higher genus inducing imperfections like holes or handles, are still present. To ensure *S*_ref_(*x, y, z*) is genus-0 for spherical parameterization, we developed a guaranteed genus-0 shrink-wrapping algorithm (Fig. S2D, Methods, Movie S2). An initial ‘tightest’ genus-0 alpha wrap of the input mesh, 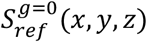 is generated^66^ with parameters found by interval bisection search (Methods). 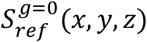 is already optimal for a *S*_ref_(*x, y, z*) devoid of surface concavities. Otherwise, 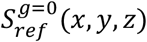 is iteratively deformed by active contours along a constructed gradient vector field (GVF)^67^ whose attractor is *S*_ref_(*x, y, z*) with punchout to ensure convergence (Methods, Movie S2). Our shrink-wrapping can be applied to arbitrarily complex *S*(*x, y, z*) (Fig. S2D,E) and to fix segmentation inaccuracies – effectively imputing large holes with a minimal bending energy surface (Fig. S2F). The 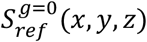 with lowest CD to *S*_ref_(*x, y, z*) is then voxelized to obtain its binary representation. The binary enables morphological dilation, hole-filling, and erosion processing for further shape refinement, notably the removal of neck-pinching singularities^63^ – sharp points and narrow necks that may impede optimal equiareal spherical parameterization (see below). The final 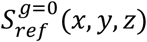 mesh (denoted *S*_*ref*_(*x, y, z*) henceforth) is obtained by remeshing the morphologically processed binary. Curvature-minimizing flows simultaneously shrink area, making holes and handles potentially smaller. Consequently, the latter voxelization without shrink-wrapping may be sufficient for 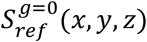, saving computing time. The genus-0 remeshed *S*_ref_(*x, y, z*) has different vertices and face topology. To restore bidirectionality between the input surface *S*(*x, y, z*) and *S*_ref_(*x, y, z*), *S*_ref_(*x, y, z*) vertices are matched by nearest normal surface distance (Methods) to a face of the intermediary reference surface that is bijective to *S*(*x, y, z*) (Methods). Any associated scalar and categorical label measurements *F*_*i*_(*S*(*x, y, z*)), including vertex coordinates are transferred to *S*_ref_(*x, y, z*) by barycentric interpolation or mode, respectively (Methods).

**Step 2** quasi-conformally maps the genus-0 reference, *S*_ref_(*x, y, z*), to the unit sphere without folds^68,69^, obtaining 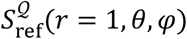 (Fig. S3D). Per the uniformization theorem, such a mapping always exists for a genus-0 surface^70-73^ but there is no unique map. u-Unwrap3D implements an efficient direct spherical projection using iterative uniform Laplacian smoothing (Methods). This is equivalent to solving Laplace’s equation with Jacobi-like iterations. u-Unwrap3D also implements an optimal direct quasi-conformal map^68^, but this is less optimal for equiareal parameterization. 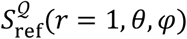 shrinks the area of surface extremities deviating from the sphere^27^, even for roughly globular shapes (Fig. S3D). The higher the curvature or longer the protrusion the greater is the undersampling and underrepresentation on 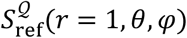 . A more conformal 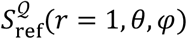 can degenerate face area and have worse face triangle quality. This can make relaxation more challenging, if not impossible. An isometric or equiareal parameterization is necessary for downstream analyses such as segmentation and tracking^27^. For pseudospherical *S*_ref_(*x, y, z*), any conformal map is already quasi-equiareal. Generally, however, the equiareal error must be diffused to ensure as uniform sampling of *S*_ref_(*x, y, z*) as possible. u-Unwrap3D implements an iterative relaxation to do this, realizing a continuous spectrum of isometric and equiareal parameterizations.

**Step 3** performs iterative bijective relaxation of the conformal parameterization, 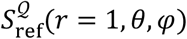 to an equiareal parameterization, 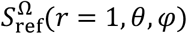 by advecting vertices without foldover, to linearly diffuse the induced per face area distortion factor of 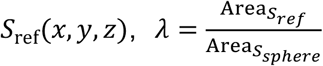 on the sphere^27^ (Fig. S3E, Methods). With this objective, relaxation can depend on the shape of *S*_ref_(*x, y, z*). Neck pinching singularities^63^ – sharp points and narrow-necked branches – commonly possessed by *S*_ref_(*x, y, z*) generated by guided curvature-minimizing flow and spectral decomposition, impede the diffusion process. Consequently, u-Unwrap3D also introduces the proxy distortion factor, 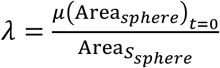 where *μ*(Area_*sphere*_)_*t*=0_ is the initial mean face area of 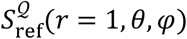 at iteration *t* = 0 (Methods). This objective exploits the area-shrinking property of conformal maps to equivalently reframe area distortion relaxation as the uniformization of face area. Because this factor is independent of *S*_ref_(*x, y, z*), optimization is more stable, and optimal relaxation can be achieved for more diverse and complex shapes. Equiareal parameterization trades off with increased conformal error (Fig. S3E). Depending on the analysis, this may not be ideal. An equiareal parameterization relaxes long protrusions and extended branches at the expense of overly compressing and distorting the representation of flatter surface regions. u-Unwrap3D’s iterative relaxation scheme allows selection of a stopping iteration to achieve a specified equiareal distortion, or the minimization of a weighted sum of conformal and equiareal distortion (Methods).

**Step 4** bijectively unwraps the quasi-equiareal sphere 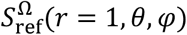 to the 2D plane, *S*(*u, v*) using an equirectangular projection, where (*u, v*) corresponds to (*θ, φ*), the spherical polar, *θ* and azimuthal, *φ* angles, respectively. The aspect ratio of the map is *N* x *2N* pixel (*N* being user-specified). This gives an isometric map of the sphere, preserving the length ratio of the north-south pole great arc to the equator circumference. When the input surface mesh, *S*(*x, y, z*) is already genus-0 or shrink-wrapped to be so, we can set *S*_ref_(*x, y, z*) = *S*(*x, y, z*), and skip Step 1, to obtain directly the 2D uv map of *S*(*x, y, z*) (Movie S1). The equiareal distortion of unwrapping either spherical parameterizations, 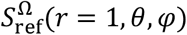, or *S*^Ω^(*r* = 1, *θ, φ*) is identical. However, *S*^Ω^(*r* = 1, *θ, φ*) has larger conformal error, (Fig. S3F). Fig S3G,H compares the reconstructed *S*(*x, y, z*) from (*u, v*) unwrapping of four spherical parameterizations: i) direct projection to the sphere^13,53^, *S*(*x, y, z*) → *S*(*r, θ, φ*) ; ii) conformal sphere, *S*(*x, y, z*) → *S*^𝒬^(*r, θ, φ*) ; iii) equiareal sphere, *S*(*x, y, z*) → *S*^Ω^(*r, θ, φ*) ; and iv) reference sphere, 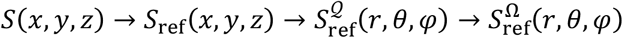. Options i) and ii) produce the largest error, and loses significant portions of the surface, when the cortical shape is elongated. Importantly, option i) is also topologically incorrect, leading to severe surface area underestimation. Option iii) delivers the most accurate reconstruction but the u-Unwrap3D proposed option iv) is almost as good, with the advantage of being more generally applicable and efficient, not requiring accurate genus-0 shrink-wrapping for complex *S*(*x, y, z*) . For any spherical parameterization, an equirectangular projection significantly distorts signals mapped to the north and south poles (Fig. S3F, S4A). Moreover, the unwrapping breaks spherical symmetry, generating a spatial discontinuity at the seam-cut (Fig. S4B). u-Unwrap3D finds an optimal (*u, v*) map to represent surface features-of-interest with minimal distortion automatically (Fig. S4C). First, we find the optimal rotation matrix to map surface features-of-interest to the equator by composing the principal eigenvectors of a vertex-based weight function, *w*, scoring the feature-of-interest. For example, for protrusions, *w* is the absolute local mean curvature of *S*(*x, y, z*), |*H*(*S*(*x, y, z*))| (Fig. S4C, see Methods for additional examples of *w*). This rotation aligns the center of the (*u, v*) parameterization with the principal eigenvector axis; the *u*-axis to the second eigenvector axis; and the unwrapping axis is the third eigenvector with minimum variation of *w*. The eigenvector axes are ambiguous in orientation. To place the seam-cut with maximum spatial continuity across the features-of-interest, u-Unwrap3D inverts the axes as necessary by a further rotation to ensure the summed contribution of a weighting function, *w*′ projected along the principal eigenvector is positive. *w*′ must be signed such that the more a vertex expresses a property of the target features-of-interest, the more positive its value. An example is the mean curvature, *w*^′^ = *H*(*S*(*x, y, z*)) (see Methods for more). u-Unwrap3D further optimizes the aspect ratio of the UV map. A *N* x 2*N* map is only optimal for rotationally-invariant spherical *S*_ref_(*x, y, z*) (Fig. S4F). For general, elongated or branched shapes, this aspect ratio is suboptimal. The resulting map, when the major shape axis corresponds to the *v*-axis, will be severely distorted. u-Unwrap3D finds an optimal width scaling factor, *h* to compute a distortion-minimized *N* x *hN* pixel (*u, v*) parameterization by minimizing a generalized length ratio or the Beltrami coefficient (Fig. S4G, Methods). The ability to optimize both unwrapping axis and aspect ratio means u-Unwrap3D, unlike other implementations of UV mapping^47,49,51,74^, can unwrap a drosophila embryo with minimal distortion along either the shorter dorsal-ventral axis or the longer antero-posterior axis minimal distortion without explicit manual rotation (Fig. S4H).

**Step 5** samples and remaps a Cartesian 3D volume *V*(*x, y, z*) and associated signals *I*_*i*_(*V*(*x, y, z*)) proximal to *S*_ref_(*x, y, z*) into a depth-extension of (*u, v*) space - a topographic volume *V*(*d, u, v*) coordinate system whereby the plane *V*(*d* = *d*_*ref*_, *u, v*) is the (*u, v*) parameterization of *S*_ref_(*x, y, z*), *S*_ref_(*u, v*) ; and *V*(*d* > *d*_*ref*_, *u, v*), *V*(*d* < *d*_*ref*_, *u, v*) corresponds to the volumetric space external and internal relative to *S*_ref_(*x, y, z*). We construct the *V*(*d, u, v*) space, and establish the mapping of (*d, u, v*) to (*x, y, z*) coordinates by propagating *S*_ref_(*u, v*), the (*u, v*) parameterization of *S*_ref_(*x, y, z*) at equidistant steps of *α* voxels along the gradient of a distance transform, 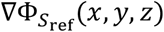 of *S*_ref_(*x, y, z*), for a total of *D* = *D*_*in*_ + *D*_*out*_ steps: *D*_*in*_ steps inside and *D*_*out*_ steps outside. *D*_*in*_ and *D*_*out*_ can be user-specified or automatically set, *D*_*in*_ as a percentile of the inner distance transform and *D*_*out*_ by the hausdorff distance between *S*_ref_(*x, y, z*) and *S*(*x, y, z*), respectively. *V*(*d, u, v*) compresses the external volume and expands the inner volume (Fig. S5A). u-Unwrap3D uses the Euclidean distance transform to perform stable outwards propagation using forward Euler with (*u, v*)-enabled uniform smoothing regularization (Methods). For inner propagation the method and distance transform, 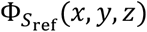 are typically chosen to tradeoff speed and accuracy (Methods). For convex *S*_ref_(*x, y, z*) or superficial propagation inside, forward Euler and Euclidean distance transform (EDT) is fastest. For general *S*_ref_(*x, y, z*), like mean curvature flow, EDT leads to neck-pinching singularities, and loss of bijectivity. The Poisson distance transform (PDT)^63,75^ is used instead, which produces a conformalized flow to stably propagate *S*_ref_(*x, y, z*) by active contours. EDT and PDT converge onto the medial-axis skeleton^76^. u-Unwrap3D can also compute a Poisson distance transform to converge to any internal shape, specified by a binary segmentation, by solving the Poisson diffusion equation with the shape as a source^76^. This latter approach allows bijective mapping of the entire internal volume of a complex *S*_ref_(*x, y, z*). Finally, u-Unwrap3D can resample the equidistantly constructed topographic space so that all vertices sample the propagated Cartesian space in the same number of steps. Trilinear interpolation of the Cartesian volumetric-associated measurements, *I*_*i*_(*V*(*x, y, z*) at the (*x, y, z*) coordinates indexed by (*d, u, v*) generates the topographic 3D volume equivalents, *I*_*i*_(*V*(*d, u, v*)).

**Step 6** generates the topographic 3D surface representation, *S*(*d, u, v*) . The Cartesian binary volume segmentation, *B*_*i*_(*V*(*x, y, z*)) is mapped into the topographic space constructed in Step 5 to obtain its topographic-equivalent binary, *B*_*i*_(*V*(*d, u, v*)). Then, *B*_*i*_(*V*(*d, u, v*)) is surface meshed to obtain *S*(*d, u, v*). To establish bidirectionality, *S*_*topo*_(*x, y, z*) is matched to *S*(*x, y, z*) by nearest normal surface distance. *S*_*topo*_(*x, y, z*) are the vertices of *S*(*d, u, v*) mapped to its Cartesian 3D coordinates using the (*d, u, v*) to (*x, y, z*) correspondence of *V*(*d, u, v*) . To restore bidirectionality of *S*(*d, u, v*) to *S*_*ref*_(*u, v*), u-Unwrap3D develops a topographic cMCF to flatten *d* to be uniformly constant, *S*(*d* = constant, *u, v*) whilst maintaining the rectangular boundary in each iteration (Methods, Movie S3).

### u-Unwrap3D maps diverse surface motifs on complex cell geometries

Fig. 2A summarizes u-Unwrap3D’s bidirectional mappings of a genus-X surface between 5 equivalent surface representations; Cartesian 3D, *S*(*x, y, z*), a 3D genus-0 reference, *S*_ref_(*x, y, z*), the unit 3D sphere, *S*(*r* = 1, *θ, φ*), the 2D plane, *S*(*u, v*) and topographic 3D, *S*(*d, u, v*). u-Unwrap3D also remaps Cartesian 3D to topographic 3D volumes (Fig. 2B). To demonstrate the broad applicability of this framework Movie S1 and Fig. 2C showcases the representations of five cellular and one multicellular surface, selected to represent a diverse spectrum of complex cell shapes and protrusions. For each, we indicate segmented protrusions or mean curvature of *S*(*x, y, z*) as surface signals *F*_*i*_(*S*(*x, y, z*)) and transfer them to all other representations. The first three examples, selected from a previously published dataset of 68 cells^39^ with protrusions detected by u-Shape3D, show the mapping of blebs, lamellipodia and filopodia, despite significant variation of cortical shape. The performance was quantitatively assessed using 4 metrics to measure the reconstruction accuracy of the final representation, *S*(*d, u, v*), remapped to 3D Cartesian, *S*_*topo*_(*x, y, z*), with the input shape, *S*(*x, y, z*) . We also tested the impact of different *S*_ref_(*x, y, z*) . In all cases, salient protrusions were reconstructed. However, *S*_*ref*_(*x, y, z*) found by cMCF using the rate of decrease in absolute *K* without shrink-wrapping was less specific for isolating protrusions. Therefore, less prominent filopodia were underrepresented in *S*(*d, u, v*). Using a tighter *S*_ref_(*x, y, z*) found by changepoint detection and genus-0 shrink-wrapping produces a more accurate representation that does reconstruct all protrusions (Fig. S5C). The fourth example maps a T cell with microvilli. Here, the microvilli labels were segmented by unsupervised clustering and connected component analysis on the mean curvature of *S*(*x, y, z*), which is enabled by u-Unwrap3D as an alternative to u-Shape3D surface motif segmentations (see below). This cell was imaged under timelapse where its cortical shape remained largely unchanged. Instead of finding *S*_*ref*_(*x, y, z*) from *S*(*x, y, z*), we found and used the mean temporal shape as the reference surface for all timepoints (Methods). As seen in Movie S1, the microvilli are dynamic: bending, fissioning, fusing and translocating on the surface. Tracking the microvilli directly in Cartesian 3D is challenging, however tracking their basal movement in *S*_ref_(*u, v*) or even 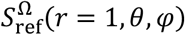 is significantly more feasible. Example 5 shows a time series of instance segmentation masks simulating a cell with branch morphogenesis using the software, FiloGen^77^. We meshed the binary of the masks to extract *S*(*x, y, z, t*), with unique coloring of each branch. To ensure *S*_ref_(*x, y, z, t*) was aligned with the branches at each timepoint, we used the minimum curvature shape generated by a guided MCF (Fig. S1C). The hierarchical nature of the branching and its development is readily evident and visualized in *S*_ref_(*u, v*) where secondary branches are mapped into the area occupied by their primary branch. Finally, example 6 remaps a cell aggregate^78^ whose overall composite surface exhibits a diverse mixture of surface motifs. Notably, using u-Unwrap3D, we remapped an independent instance cell segmentation of the same aggregate by u-Segment3D^76^ into topography space, to transfer the segmentation labels and identify the boundaries of unique cells in *S*(*x, y, z*) (Movie S1).

**Figure 2.**
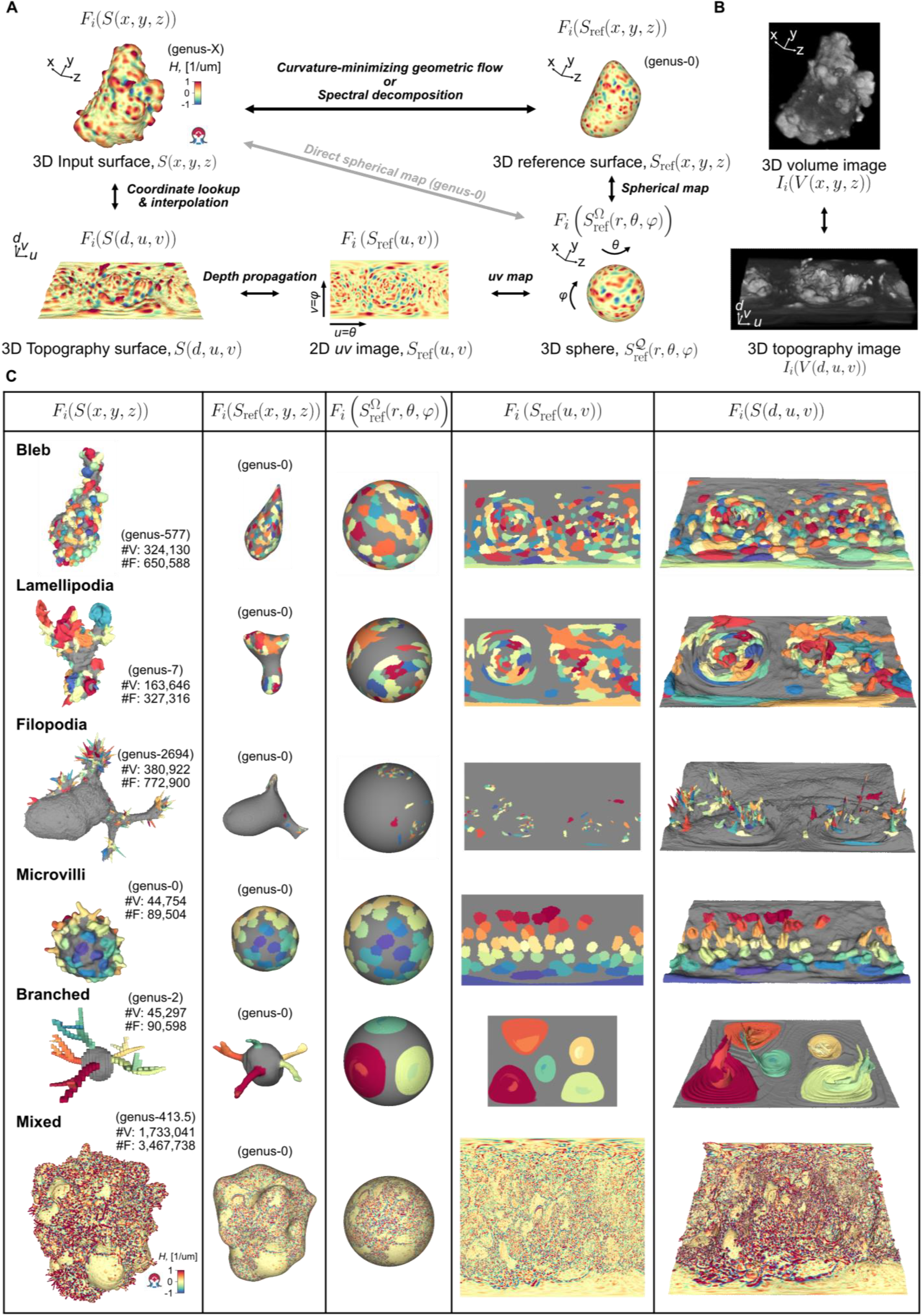
u-Unwrap3D generates a spectrum of equivalent data representations for surface-guided computing and is applicable to diverse surface protrusion morphologies. **A)** Summary of the bijective mappings between the 5 equivalent surface mesh representations generated by u-Unwrap3D. Black bidirectional arrows indicate the mapping algorithms between representations discussed in the text. Grey arrow indicates the direct spherical mapping applicable when the input mesh is genus-0. **B)** u-Unwrap3D also enables bidirectional mapping of volumetric information between a Cartesian and topographic space relative to a genus-0 reference surface. **C)** Gallery of equivalent surface representations generated on examples of cells with bleb, lamellipodia, filopodia, microvilli, branched and mixed surface protrusion morphologies. For visualization of the mappings, individual instances of morphological motifs detected by the software u-Shape3D (bleb, lamellipodia, filopodia) or u-Unwrap3D (microvilli) or provided (branched) are color-coded on surface representations. Mixed is visualized by mean curvature, *H*(*S*(*x, y, z*)) coloring.

Importantly, the bidirectionality of the mappings guarantees that for any point on any of the surface or volume representations matching points exist on any of the other surfaces or volumes. Moreover, the bidirectionality preserves the point topology, i.e. a series of points ordered in clockwise fashion on one surface representation maps to a series of points ordered in the same way on any of the other surface representations and preserves the local neighbourhood relationships. As a result, we can apply mathematical operations defined in any one of the representations and map the results to any other. u-Unwrap3D thus supports the optimal spatiotemporal analysis of unconstrained surface geometries and associated signals.

### u-Unwrap3D establishes inherent spatiotemporal correspondence between surfaces from timelapse image sequences

For cells segmented from volume timelapse imaging, the standard workflow illustrated in Fig. 3A applies u-Unwrap3D to the cell surface extracted in each timepoint, *S*(*x, y, z, t*). The main consideration differing from application to a single shape is the requirement for temporal consistency among maps of consecutive timepoints (Methods). The first step to achieving consistency is to pre-register the timelapse sequence or *S*(*x, y, z, t*) to remove global translation, rotation, and shear. Of note, scaling is inherently removed by mapping to the unit sphere. Second, several parameters in the mappings must be set globally throughout the sequence: (i) using the same stopping iteration or a temporal smoothing of the stopping iteration to ensure a continuous evolution of *S*_ref_(*x, y, z, t*); (ii) using a representative temporal reference shape to determine an unwrapping axis and aspect ratio for (*u, v*) parameterization; and (iii) using the same number of *D*_*in*_ and *D*_*out*_ steps to construct the same volumetric sized (*d, u, v*) space and align *S*_ref_(*x, y, z, t*) at the same *d* value. With these guidelines, mapping all *S*_ref_(*x, y, z, t*) to the static sphere and plane geometry automatically establishes their groupwise spatiotemporal correspondence, even when changes in *S*_ref_(*x, y, z, t*) are moderately large over the entire timelapse. In doing so, protrusions captured implicitly by scalar measurements, such as mean curvature, or explicitly by geometric reconstruction exhibit consistent dynamics in *S*_ref_(*u, v*) and *S*(*d, u, v, t*), respectively (Fig. 3A, Movie S4). If *S*_ref_(*x, y, z, t*) undergo large shape deformations over the timelapse duration, u-Unwrap3D implements functions leveraging the SimpleITK package for volumetric registration of *S*_ref_(*x, y, z, t*) using multiscale symmetric diffeomorphic demons^79,80^. Any inaccuracies in the displacement fields between the moving and reference shape are corrected for by u-Unwrap3D propagating the transformed moving shape along the gradient of a distance transform with the reference shape as attractor, similar to shrink-wrapping (Methods). Notably, using a time-varying *S*_ref_(*x, y, z, t*) effectively decouples local protrusion dynamics in *S*(*d, u, v, t*) from cell-scale movements. This feature is illustrated by Movie S4, where bleb protrusions from an elongated cell are transferred to a spherical cortical shape, i.e. *S*(*d, u, v, t*) is re-embedded into the (*d, u, v*) space of a sphere, then yielding the Cartesian 3D equivalent, *S*(*S*_*target*_(*r, θ, φ*), *t*) where target = sphere (Fig. 3B, Movie S4). This principle holds broadly, even when the blebs are decorated on the branched surfaces of a cell (Fig. 3C, Movie S4).

**Figure 3.**
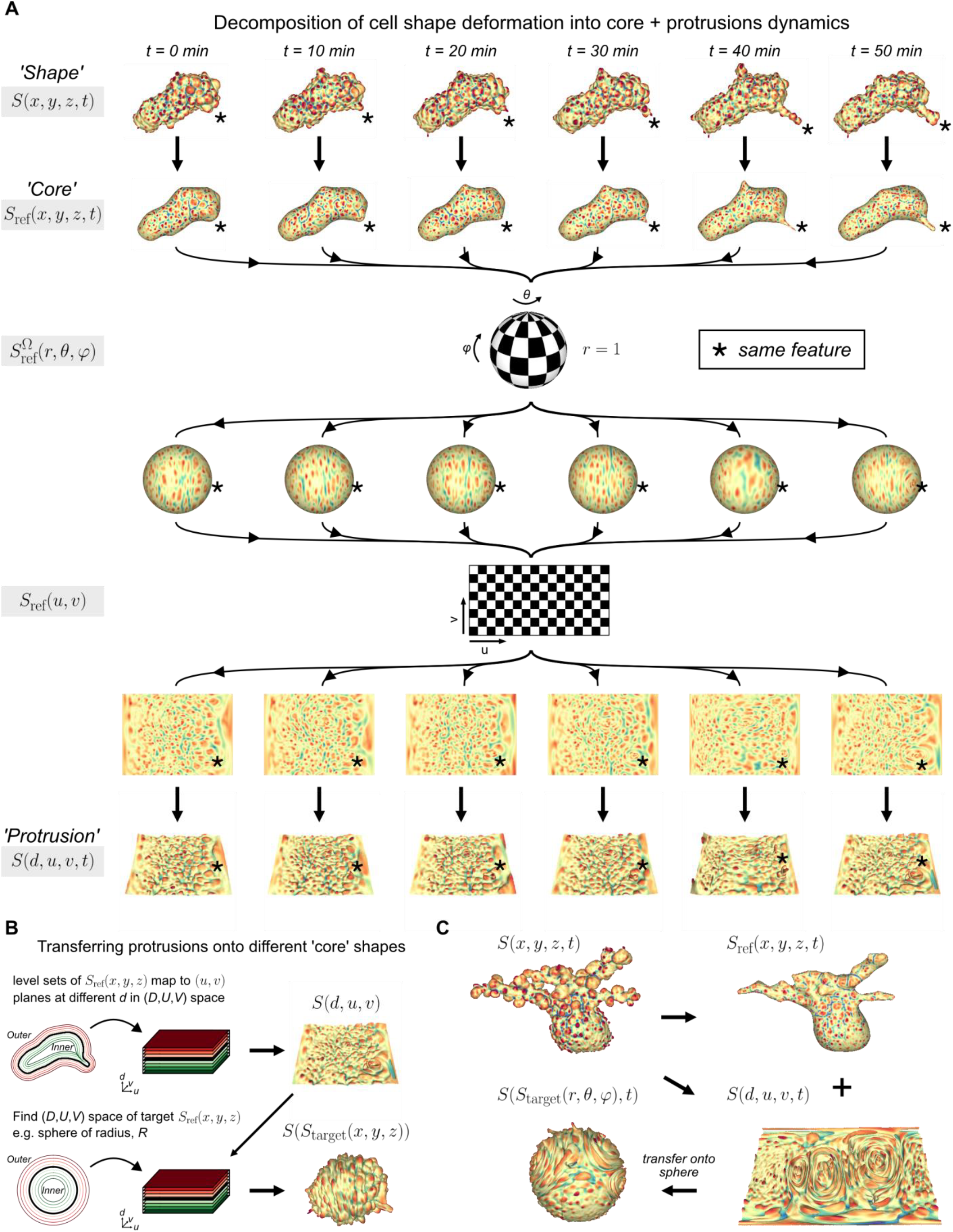
u-Unwrap3D decomposes 3D cell surface into core and protrusion morphodynamics. **A)** Application of u-Unwrap3D to individual timepoints to find the timepoint-specific reference shape representing the core ‘global’ cell shape. These references are used to map the surfaces into canonical spherical and unwrapped (u,v) coordinate systems. Performing this mapping puts the core shapes across all timepoints in correspondence, such that topography reconstruction isolates protrusion dynamics from core shape. **B)**. Procedure to transfer protrusions from one shape to another core shape by specifying the (*d*_*source*_, *u*_*source*_, *v*_*source*_) ↔ (*x*_*source*_, y_*source*_, *z*_*source*_) coordinate correspondence from source to the Cartesian coordinates of the target shape, (*d*_*source*_, *u*_*source*_, *v*_*source*_) ↔ (*x*_*target*_, y_*target*_, *z*_*target*_). This transfer is an immediate corollary of the decoupling between protrusion and core shapes. **C)** Transfer of a challenging surface morphology composed of blebs decorating a branched core onto a sphere geometry.

A central goal of live cell imaging in 3D is to visualize the spatiotemporal relations between molecular activities and cell behaviors. Progress has been made on software developments that allow unbiased and statistically meaningful analysis of cell morphology and molecular distributions^13,39,81,82^. Surface-guided computing with u-Unwrap3D now allows us to capture and quantify dynamic morphological and molecular behaviours on complex 3D cell shapes. Geometrical measurements, segmentation, temporal tracking, correlative and causal analyses can be conducted in the best-suited surface representation and results, if of interest, visualized in 3D by leveraging the bidirectional properties of the u-Unwrap3D mappings. The remainder of the paper demonstrates through four case studies, the flexible adaptation of the described u-Unwrap3D workflow to solve challenging spatiotemporal problems out of reach for computation in Cartesian 3D.

### NK cells establish retrograde flow of surface protrusions in cytolytic immunological synapse formation with cancer

Natural Killer (NK) cells are key components of the innate immune system. They recognize and destroy virally infected or oncogenically transformed cells by forming a multi-purpose interface called the immunological synapse with their target^83^. Actin rearrangements^84,85^ are critical for a productive outcome of the cytotoxic process, however, the interplay between the shapes of NK and target cells, remain poorly understood^86^. Direct imaging of heterotypic cell-cell interactions is notoriously difficult, given these interactions are random and often short lived. We apply u-Unwrap3D to interrogate the surface dynamics of an NK cell expressing a filamentous actin reporter before and during cytolytic immunological synapse with a myelogenous leukemia expressing a plasma membrane fluorescent marker, imaged by high-resolution light-sheet microscopy^86^ (Fig. 4A). The NK and cancer cell are located at the periphery of the imaged field of view, and only partially in-view initially. Fig. 4B illustrates the specific u-Unwrap3D temporal workflow. To obtain complete and temporally consistent cell surfaces throughout, we used u-Segment3D^76^ supported by 2D cellpose^87^ segmentation models and temporal tracking (Methods). We find the cortical shapes, *S*_*ref*_(*x, y, z, t*) using cMCF, which highlighted a dynamic remodeling of *S*_ref_(*x, y, z, t*), from being branched, presumably more optimal for locomotion pre-contact, to a vertically-oriented dumbbell-like shape, one head directly engaging the cancer cell, the other diametrically opposite. As *S*_ref_(*x, y, z, t*) undergoes significant deformation, we register all *S*_ref_(*x, y, z, t*) using symmetric diffeomorphic demons^79^ to a single static reference - the temporal mean of *S*_ref_(*x, y, z, t*), ⟨*S*_ref_(*x, y, z, t*)⟩ or ⟨*S*_ref_⟩(*x, y, z*) . To map the morphological development from the perspective of the interface, we found the timepoint when a stable synapse is established. Then, the contacted surface area with the cancer cell was mapped using the established registration to ⟨*S*_ref_⟩(*x, y, z*) as the weighting function, *w* to determine ⟨*S*_ref_⟩(*u, v*) maps centering the synapse formation process. Mapping the surface protrusions, captured by mean curvature, *H*(*S*(*x, y, z*)) into 2D ⟨*S*_ref_⟩(*u, v*), Movie S5 reveals dynamic ridge-like protrusion formation accompanying the ‘expansion’ of the synapse. These protrusions may be responsible for the previously observed actin accumulation and trogocytosis^86^. The random and transient protrusion dynamics challenges optical flow tracking using pairwise frames (Movie S5). Instead, we computed multiscale information flow using u-InfoTrace^88^, a framework we previously developed to apply 1D causal inference measures for 2D video analysis, using the whole video duration to extract causal relationships between individual pixels. Specifically, we use the dynamic differential covariance (DDC)^89^ measure suited for capturing short-time dynamics. Notably, u-InfoTrace can be readily adapted for *H*(⟨*S*_ref_⟩(*u, v*)), with simple correction of the metric distortion through reweighting of signals (Methods). This revealed a consensus surface retrograde flow of protrusions emanating from the center of the synapse to the negative curvature midriff of the cell. Interestingly, the flow does not proceed to the opposite head, where instead we observe a recirculating pattern of protrusion flow. This flow may serve a different function, potentially to help maintain continual physical engagement with the cancer cell. These unprecedented observations highlight the power of u-Unwrap3D to quantify activity at cell-cell interfaces in relation to the rest of the surface, which due to the surface curvature, and transience of protrusions, severely complicate a direct Cartesian 3D approach.

**Figure 4.**
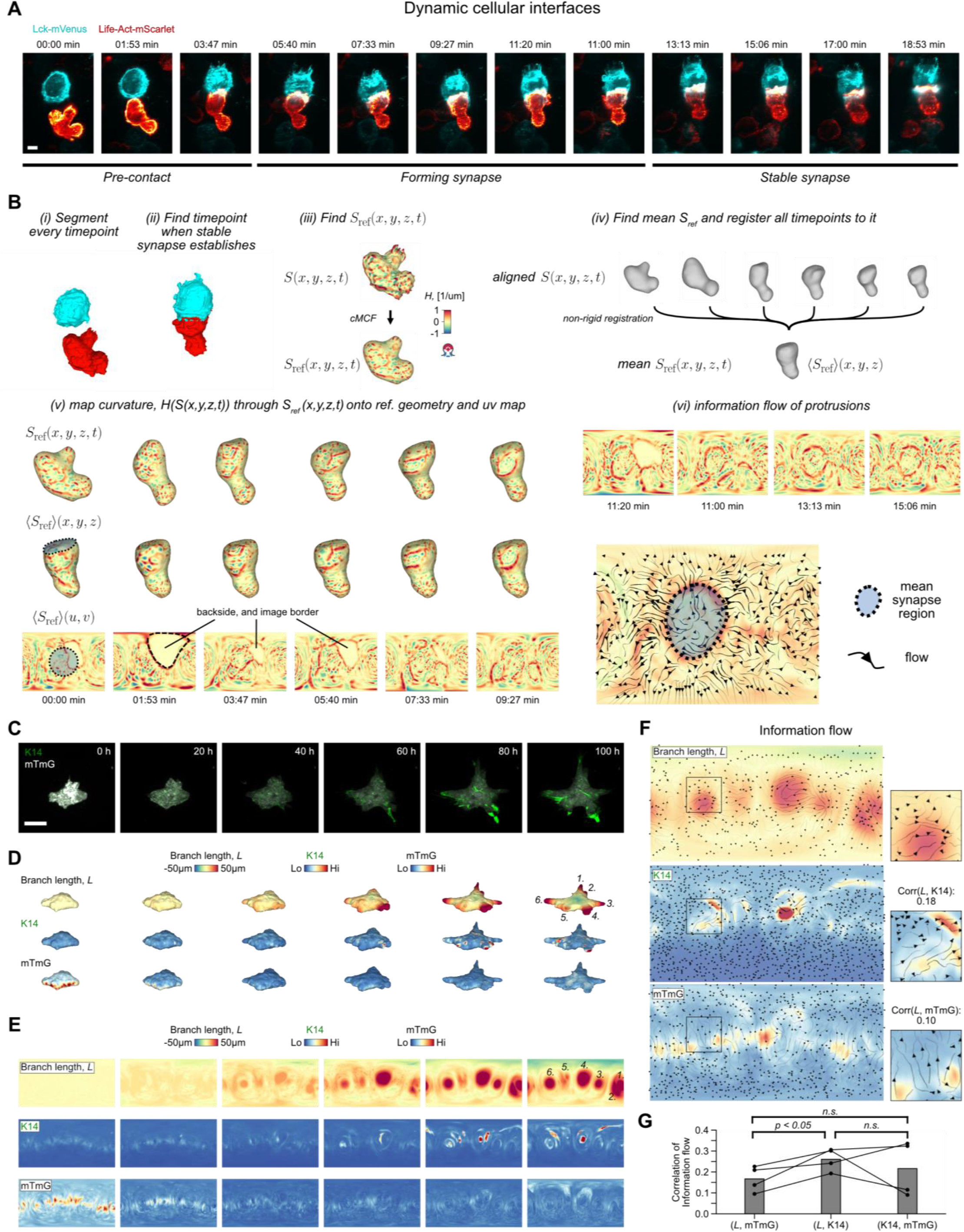
u-Unwrap3D enables causal investigation of morphology and signaling at dynamic interfaces. **A)** Snapshots of a NK-92 natural killer cell expresssing Life-Act-mScarlet before and after contact interaction with a K562 leukaemic cancer cell expressing Lck-mVenus in-vitro leading to the formation of a stable immunological synapse^86^. Scale bar: 20µm. **B)** Steps to map surface protrusions captured implicitly by the mean curvature of *S*(*x, y, z*) to canonical (u,v) image representation for visualization and tracking. **C)** Snapshots of breast cancer spheroids expressing mTmG and keratin 14 (K14) positive leader cells that locate at the tips of the invasive branches^91^. Scale bar: 200 µm. **D)** 3D surface mesh of images in C) colored by branch length relative to initial shape (Methods), and surface proximal intensities of K14 and mTmG. **E)**. (*u, v*) unwrapped image equivalent representation of measurements in D). **F)** Streamplot of dynamic differential covariance (DDC) information flow^88^ overlaid on the temporally averaged unwrapped organoid branch length, K14 and MTMG signal intensities of E), top-to-bottom. Insets zoom in on the flow surrounding a representative organoid branch. **G)** Barplot of the ensemble mean (n=4 organoids) average correlation between the information flow vector fields of measurements *A* and *B*, denoted (*A,B*). Individual organoids represented by points, with line joining correlation of measurements made in the same organoid. Statistical difference in flow correlation assessed using a paired t-test at a significance level of 0.05.

### K14+ cells causally associate with the leading edge of invasive strands in breast cancer organoids

Carcinomas typically invade as a cohesive multicellular unit, a process termed collective invasion. Characterization of invasion in tumour organoids derived from MMTV-PyMT mice^90^, a model of highly invasive breast cancer, previously identified a population of leader cells, characterized by basal epithelial genes, particularly cytokeratin-14 (K14) that may drive the development of invasive strands^91^. Imaging MMTV-PyMT mouse-derived tumor organoids co-expressing K14 and the membrane marker, mTmG, shows colocation of K14+ cells with invasive strands. However, K14+ cells also locate elsewhere in the organoid. Additionally, not all invasive strands have K14+ cells nor do they have the same number when they do (Fig. 4C, Movie S5). Lastly, whilst the invasive strand continuously grows outwards, K14+ cells also migrate, and change shape, extending long invasive protrusions spanning a large surface area that covers multiple K14-cells (Fig. 4D, Movie S5). All these suggest the need to quantitatively assess if K14+ cells causally drive invasive strand formation.

We use u-Unwrap3D to map the organoid surface and surface-proximal signal intensities in (*u, v*) images. As a means of inferring causality between the two variables we computed multiscale information flow of each signal with u-InfoTrace^88^ (Fig.4E, Movie S5, Methods). We determine a branch length, *L* everywhere on the surface at time, *t* using the total curvilinear distance of propagating a *S*(*x, y, z, t*) vertex along the gradient of the distance transform with *S*(*x, y, z, t* = 0) as the attractor (Methods). The surface-proximal K14 and mTmG signals were sampled as part of constructing the (*d, u, v*) topography space using the Poisson distance transform^75,76^. The maximum intensity projection of the topographic molecular signals up to a depth of 5 µm from *S*_ref_(*x, y, z, t*) was computed as the surface-proximal signal-of-interest in *S*_ref_(*u, v*). We rationalized that if K14+ cells and invasive tips were related, then we should find the pattern of information flow to be more similar between K14 and *L* than between mTmG and *L*. The similarity should be as least as high as between the information flows of K14 and mTmG as K14+ cells are sparse. Applying two-tailed paired *t*-tests to the flow pattern similarity computed by normalized cross-correlation (Methods), we found this was indeed the case (Fig. 4G, *n* = 4 videos). This example underscores how u-Unwrap3D routines both facilitated the measurement of a branch length in Cartesian 3D and enabled simpler 2D flow-based techniques to test a causal association between surface-associated parameters in a context where the 3D geometry is dynamically changing.

### Ruffles are driven by locally enriched filamentous actin and migrate actively on the cell surface

We also applied the capacity of u-Unwrap3D to examine putative relations between the dynamics of membrane ruffles, filamentous actin and surface actin retrograde flow. Membrane ruffles are thin, rapidly-moving, actin-rich protrusions^92,93^ proposed to arise as a consequence of inefficient adhesion in cellular lamellipodia^94^. Although ruffles have been the showcase for many volumetric light sheet microscopy advances, there are few quantitative assessment of membrane ruffling^92,95^, and even fewer analyses relating ruffle formation with molecular processes. The thin, lamellar appearance make ruffles extremely challenging to segment in 3D live-cell images^96^. Moreover, ruffles are transient, and they exhibit significant heterogeneity in nature and distribution across the cell surface. These characteristics make ruffles ill-defined compared to more distinctive surface motifs like blebs, lamellipodia and filopodia, and challenging, if not impossible to track individually in Cartesian 3D.

There are many unanswered questions about cell membrane ruffles. Here, we focus on objectively defining a membrane ruffle and measuring the speed of their traversal across the cell surface. We acquired 3D volumes of SU.86.86 pancreatic ductal adenocarcinoma cells co-expressing Tractin-mEmerald and myristoylated CyOFP1. Tractin is a marker for actin^97^, while myristoylated CyOFP1 served as a diffuse cell membrane marker. We acquired images every 2.27s for 30 timepoints, and used u-Unwrap3D to map the Cartesian 3D *S*(*x, y, z*) surface of every timepoint into topographic 3D, *S*(*d, u, v*), and into 2D, *S*(*u, v*), representations (Fig. 5A, Movie S6). The cortical shape change was relatively small, but ruffles travel on the surface. We therefore determined *S*_ref_(*x, y, z, t* = 0) by cMCF (Fig. S7A), and used it as the reference surface for all timepoints to define a common static topographic coordinate space (*d, u, v*). We then tracked the spatial location of ruffles as individual ‘ridge’ objects, directly projecting *S*(*d, u, v*) into the 2D plane, i.e. using the (*u, v*) coordinate and setting *d* = 0, (*d* = 0, *u, v*). We computed the local actin enrichment as the ratio of Tractin-mEmerald to the cytosolic CyOFP1 intensity. As expected, kymograph visualization of mean curvature, *H*(*S*(*u, v, t*)) and Tractin-mEmerald/CyOFP1 (TC) showed strong co-fluctuation. It also highlighted the transient, ‘ripple-like’ nature of ruffles and their merging and dissipation with a delay between successive ruffles of ≈20s (Fig. 5B). We applied optical flow-based region-of-interest (ROI) tracking^98,99^ to the TC intensity to track simultaneously the protrusive membrane ruffling and retrograde surface actin flow (Fig. 5C, left). The average dimensions of the tracked (*u, v*) ROI size was 23 x 23 pixels (≈ 0.06 x 0.11 μm pixel size in physical space). The 2D ROI tracks exhibit unidirectional motion towards the nucleus as shown by coloring directionality and remapping the (*u, v*) ROI tracks to polar (*r, ϕ*) coordinates, (Fig. 5C, middle, Fig. S7B) and Cartesian 3D (*x, y, z*) coordinates (Fig. 5C, right) (Movie S6). We used the 2D optical flow tracks to measure the speed of ruffling. To measure specifically the component parallel to the lamella, we projected the 3D-equivalent optical flow velocities along the same plane as the flat cell bottom. The volumetric imaging was acquired on a cover glass tilted at ≈45^0^ (Fig. S5C, left). Directly fitting a 3D plane to *S*(*x, y, z*) vertices to determine a precise tilt angle is sensitive to outlier points. However, thanks to the *S*(*u, v*) representation, whose upper and lower half map to the curved cell surface and the planar bottom respectively, we could directly annotate in 2D (*u, v*) and fit a 3D plane to only the planar bottom surface patch remapped to (*x, y, z*) (Fig. S7C, middle, right). We similarly identified only ruffle- and lamella-associated ROI tracks in (*u, v*) to compute the lateral speed. We find two populations in the speed histogram (Fig. 5D). Visualizing the speed on *S*(*x, y, z*), the faster population corresponds to higher curvature ruffles with speeds ranging from 2-10 μm/min, and an average speed of 4.2 μm/min (Movie S6). This is at least two times faster than the slower population surface retrograde actin flow ranging from 0-1 μm/min, which is consistent with the flow speeds we used to measure by 2D fluorescent speckle microscopy in the lamella of epithelial cells^100^. This result suggests ruffles are actively produced and transported across the cell surface in a mechanism separate from the myosin-driven retrograde flow. To assess the synchronicity of actin and ruffles we extracted distortion-corrected timeseries of TC and mean curvature, *H* by sampling and averaging the respective values within the tracked spatial ROIs. Averaging the temporal cross-correlation curves (mean±95% confidence interval) over individual tracks we identify a significant positive instantaneous (lag=0) correlation of 0.2 (Fig. 5E, left). Plotting the instantaneous (lag=0) correlation with mean *H* measured of the same ROI and visualizing the instantaneous correlation on *S*(*x, y, z*), we found that ruffles with higher positive surface curvature are more temporally correlated with TC intensity (Fig. 5E, middle, right). Altogether, our results show that ruffles are dynamic, transient protrusions that actively migrate on the cell surface based on filamentous actin enrichment. Our u-Unwrap3D framework provides now the platform for systematic investigation of the molecular driver of these dynamics.

**Figure 5.**
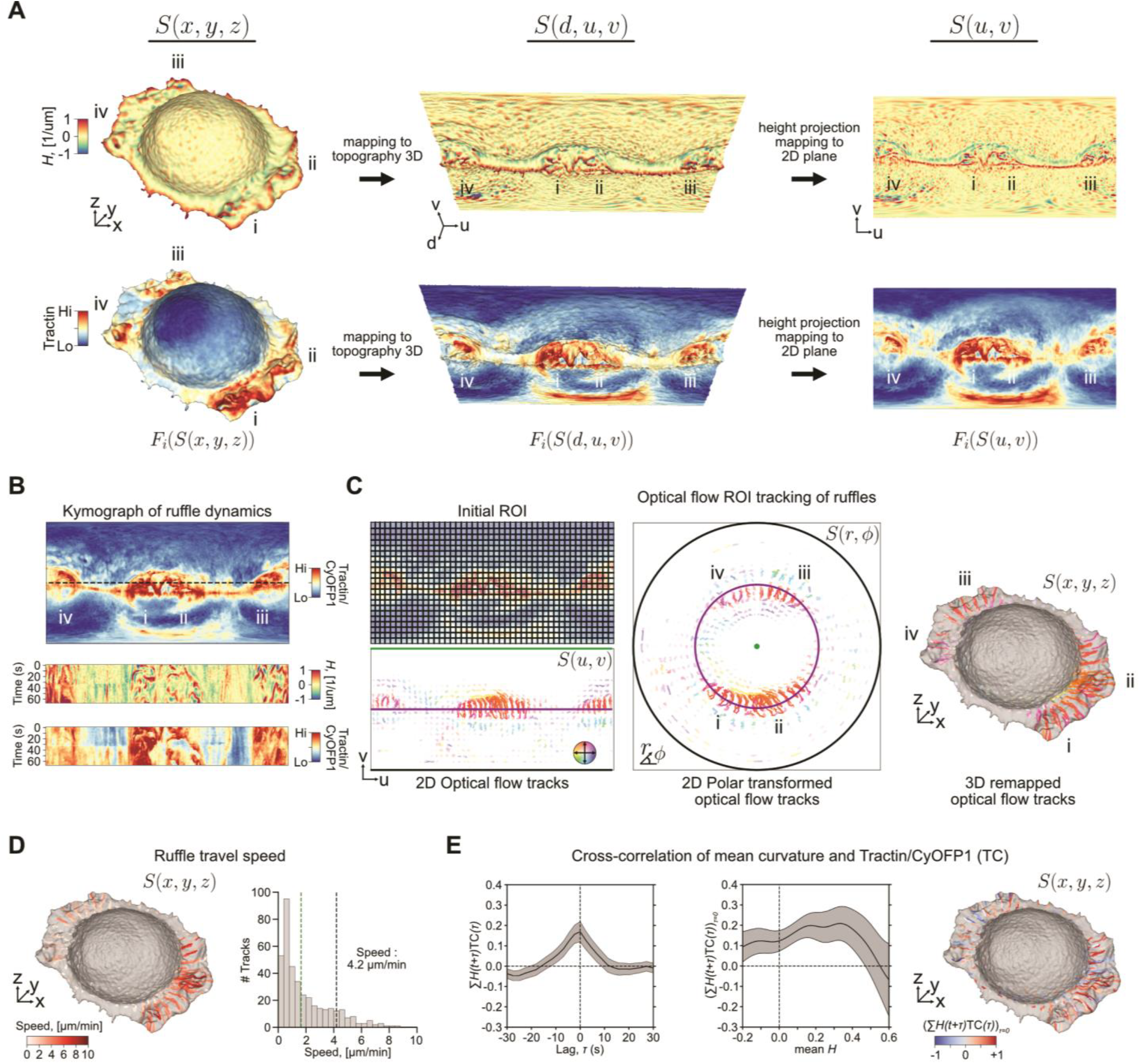
u-Unwrap3D enables tracking and characterization of morphological and molecular dynamics at the subcellular level. **A)** Unwrapping of a ruffling SU.86.86 pancreatic adenocarcinoma cell visualized in Cartesian 3D (left), topographic 3D (middle) and unwrapped 2D plane (right) representations for the first frame of a timelapse 3D image sequence sampled every 2.27s for 30 frames. Top row: mean curvature; bottom row: Tractin-mEmerald intensity sampled from a 1 μm shell proximal to the cell surface. Labels i-iv indicate corresponding landmarks in the three representations. **B)** Cross-section (dashed black line, top) to generate kymographs (bottom) of mean curvature and Tractin-mEmerald intensity normalized to myristolated CyOFP1 as a diffuse volumetric marker. **C)** Optical flow tracking on equipartitioned regions of interest (ROI) in the (u,v)-plane based on the Tractin/CyOFP1 (TC) ratiometric intensity(left). The resultant optical flow ROI tracks are colored by the mean track direction. Color saturation indicates mean track speed. ROI tracks remapped to 2D polar (*r, ϕ*) view (middle) and to Cartesian 3D (*x, y, z*) surface representation overlaid on the first time point (right). The polar transform maps the green (top), purple (middle) and black (bottom) horizontal line in the (*u, v*)-plane to the central green point, purple and black rings in the (*r, ϕ*)-view, respectively. **D)** Mean temporal planar travel speed of the ruffle-associated ROI tracks from c) plotted onto the Cartesian 3D surface representation of the first time point (left) and histogram (right). The mean ruffle travel speed of 4.2 μm/min corresponds to the faster of the two histogram populations (black vertical dashed lines) identified by 3-class Otsu thresholding (Methods). **E)** Cross-correlation curve (mean ± 95% confidence interval) between mean curvature and TC timeseries per ROI track (left). Lag 0 cross-correlation of mean curvature and TC as a function of mean curvature, *H* (middle); ROI tracks color-coded by cross-correlation magnitude plotted onto the Cartesian 3D surface representation of the first time point (right).

### u-Unwrap3D enables unsupervised instance segmentation of subcellular surface motifs

Identification and segmentation of individual cellular protrusions emerging from unconstrained 3D surface geometries is nontrivial. Even canonical morphological motifs such as blebs, lamellipodia and filopodia exhibit significant heterogeneity and ambiguity across the same cell and between cells. For example, not all blebs are spherical, lamellipodia are often plate-like with high curvature ridges but otherwise have no well-defined shape prior, and filopodia, although long and thin, can sprout from elevated ‘stumps’ or even off of each other. Especially when protrusions are dynamic and dense, it is unclear where one protrusion starts and another ends. Consequently, most existing approaches focus on specific surface features-of-interest, such as ‘ridge’ networks that can be segmented by targeted imaging filtering or through trained semantic segmentation, with morphological processing and extensive parameter tuning^96,101-103^. Few works attempt to segment protrusions generally. With u-Shape3D we introduced a multi-class morphological motif detection that partitioned the 3D surface into convex patches and applied support vector machines trained with expert annotation to classify patches into pre-specified motif types^39^. A more recent approach^104^ generated a convex hull of the surface, *S*(*x, y, z*), to define a depth function using the distance between *S*(*x, y, z*) and its convex hull boundary. Then, protrusions were detected as the extreme points of the depth function. This strategy can detect protrusion tips for counting, but does not segment the whole protrusion, and is susceptible to overdetection for motifs exhibiting multiple peaks like lamellipodia. Here, we revisit the protrusion segmentation problem using u-Unwrap3D representations to introduce conceptually simpler approaches using *S*(*d, u, v*) and volumetric representations for generalized instance protrusion segmentation.

The topographic 3D surface representation, *S*(*d, u, v*), captures in one field-of-view all surface features protruding normally to *S*_ref_(*x, y, z*). As *d* corresponds to the total Cartesian 3D curvilinear distance from *S*(*x, y, z*) to *S*_ref_(*x, y, z*) along the gradient of the distance transform, we can formally define a ‘protrusive’ feature as having a *d*-coordinate greater than that of a reference topographic surface, *S*_ref_(*d*_ref_, *u, v*). We can then measure the protrusion height as the difference, *h* = *d* − *d*_ref_. In this approach, the critical step amounts to defining the reference, *d*_ref_ = *f*(*u, v*)? An ideal *d*_ref_ should enable thresholding of *h* to specifically identify all protrusive surface features. An obvious option is to use the 2D plane, *d*_ref_ = 0 corresponding to using *S*_ref_(*x, y, z*). By default, *S*_ref_(*x, y, z*) is computed by cMCF,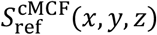. However, cMCF shrinks positive curvature features inwards and expands negative curvature outwards (Fig. S1). This causes flat surface regions to measure a positive *d*. Hence, binarizing by the threshold, 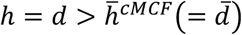, i.e. the mean value of *S*(*d, u, v*), leads to under-segmentation, and erroneous inclusion of cortical surface (Fig. 6A). Instead, we propose to define an intermediate surface, *S*_ref_(*d*_ref_ = *f*_smooth_(*u, v*), *u, v*), which interpolates between the rugged topographic cell surface, *S*(*d, u, v*) and the 2D planar cMCF cell surface 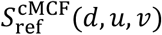. In Cartesian 3D we solve this using a positive-curvature biased guided curvature-minimizing flow (Fig. S1, Methods). However, the resulting reference may have neck-pinching singularities or correspond to the medial-axis skeleton. The measured height will be inaccurate, though still suitable for binary protrusion segmentation, which depends only on the relative height between protrusion and cortical-associated surface areas. In topographic space however, the interpolation can be naturally framed as an asymmetric least squares (ALS) optimization problem and solved using Whittaker smoothing^105,106^. The heterogeneous protrusion height is modelled by asymmetric weights; and the desired smoothness property incorporated as a regularization term (Fig. 6B, Methods). Without loss of generality, we solve ALS with a (*u, v*)-parameterized approximation of *S*(*d, u, v*) (Fig. 6B, Methods). Thresholding on the mean height with this optimal reference yields both an accurate protrusion height and the desired protrusion-specific binary segmentation (Fig. 6C).

**Figure 6.**
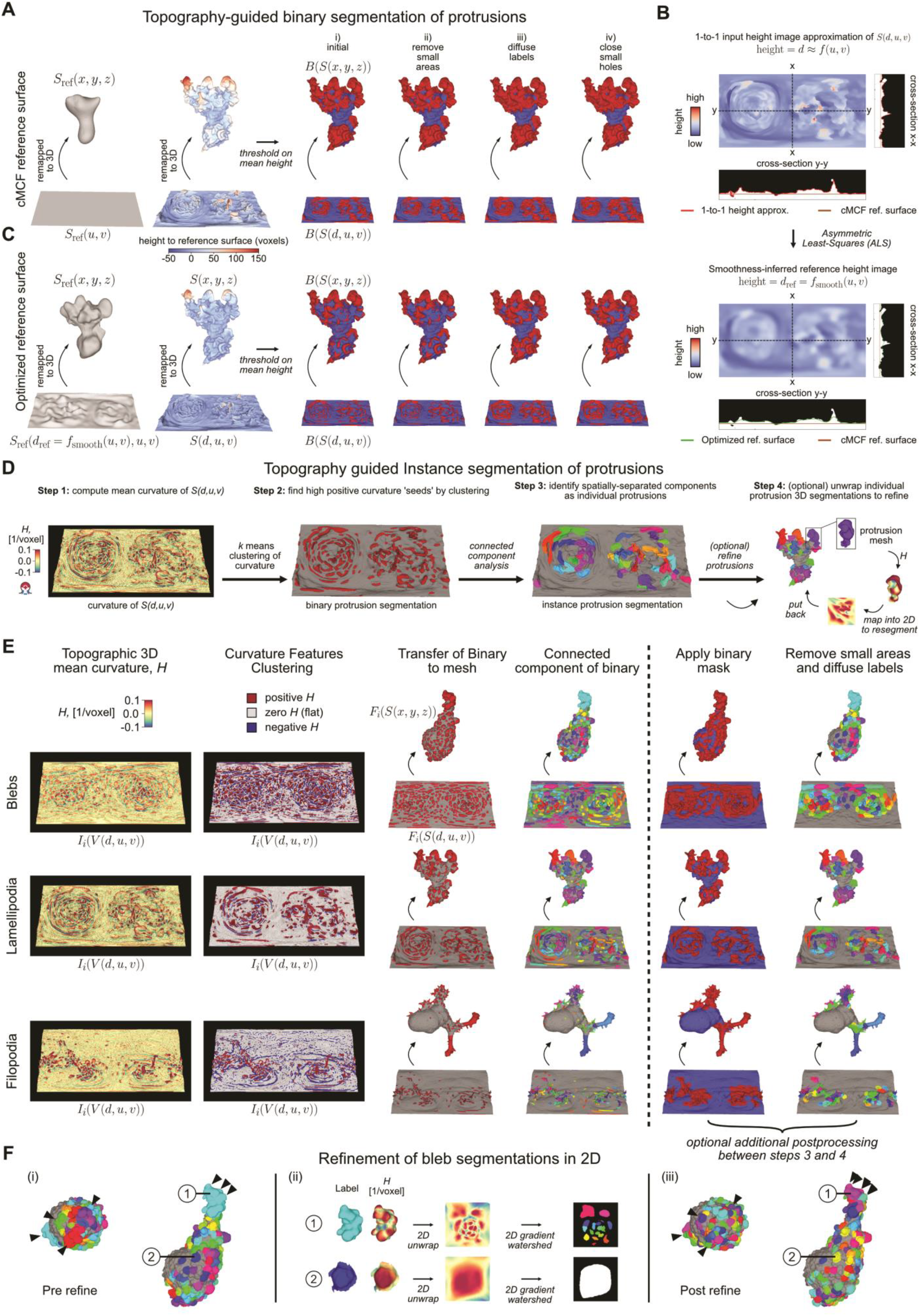
u-Unwrap3D enables unsupervised segmentation of surface protrusions guided by topographic 3D representations. **A)** Overview of the four key steps (i-iv) to binary segment protrusions by thresholding on the mean height measured relative to the cMCF reference surface. **B)** Computing an optimal reference surface for segmenting surface protrusions using asymmetric least mean squares (ALS) (Methods). The topographic surface, *S*(*d, u, v*) is approximated by a surface, *S*(*d* = *f*(*u, v*), *u, v*) that is in one-to-one correspondence to every (*u, v*) pixel and can be represented as a height image using ray-propagation (top, see Methods). The height function, *f* is depicted in cross-section cuts x-x and y-y by a red line. A flat brown line in cross-section depicts the cMCF reference, *S*_ref_(*d, u, v*). ALS with smoothness regularization is applied to *d* = *f*(*u, v*) to derive an optimal smooth reference surface with height, *d* = *f*_smooth_(*u, v*) (bottom). The height function, *f*_smooth_ is depicted in cross-section cuts x-x and y-y by a green line. **C)** Binary segmented protrusions by thresholding on the mean height measured relative to the ALS-derived reference surface from B). **D)** Overview of the sequential steps for instance protrusion segmentation by binarization and connected-component analysis of topographic volume signals, *I*_*i*_(*V*(*d, u, v*)). Undersegmented protrusions are refined in the equiareal *(u,v)* parameterization of the protrusion mesh (Methods) by applying a gradient-based watershed^76^ with flow vectors being the gradient of the mean curvature map. **E)** Instance protrusion segmentation for 3 different cell types and 3 different surface motifs; MV3 melanoma cell with blebs (top row), dendritic cell with lamellipodia (middle row) and HBEC cell with filopodia (bottom row). Initial binarization uses 3-class k-means clustering (blebs and filopodia) and 3-class Gaussian mixture model clustering (lamellipodia) of volumetric mean curvature to identify all positive curvature regions. What determines a ‘protrusion’ depends on *S*_ref_(*x, y, z*). The branching arms are therefore legitimate protrusions for the filopodia cell for the used reference (Fig. S5B). **F)** Illustration of the selective refinement of undersegmented individual protrusions. (i) Segmentation before refinement. (ii) Exemplar 2D parameterization of 2 individual segmented protrusions labelled 1, 2 into 2D (u,v) square representations for fine-grained re-segmentation of protrusions guided by mean curvature gradient. (iii) Segmentation after refinement using the bijectivity of the parameterization to map refined segmentations back onto the 3D surface. Black arrowheads highlight protrusions separated after refinement.

Exploiting further properties of the topography space, we developed an unsupervised approach to generally segment surface motifs including blebs, lamellipodia, filopodia, and microvilli with rationalized parameters to tune (Fig. 6D,E). Importantly, this approach does not rely on the design of specialized image filters^101,103^, computation and clustering of complex feature descriptors^96,107-109^, or data training^39,96,102^. By construction, in topographic 3D (*d, u, v*) space, the curvature of *S*_ref_(*x, y, z*) is integrated out and all surface protrusions are oriented upwards with increasing *d* . The tops of individual protrusions are therefore more individually separated and still detectable as local regions of high topographic mean curvatures. We segment the tops by clustering curvature features at multiple spatial scales and apply connected component labelling in the topographic volume, *V*(*d, u, v*) to obtain instance protrusion masks (Fig. 6D). The volume-based masks are transferred to label the topographic mesh, *S*(*d, u, v*). These initial ‘seed’ labels are diffused on the mesh using an affinity matrix based on geodesic distance and local dihedral angle, to also segment the ‘stem’ of individual protrusions (Methods). The dihedral angle measures the discontinuity in local mean curvature. It incorporates the prior intuition that a label on local surfaces of homogeneous curvature, e.g. atop a ‘hill’ should diffuse faster than labels with large local curvature variation such as a valley between multiple ‘hills’. The combined affinity matrix introduces a topography-aware competition between segmentation labels and provides a biophysical rationale for defining which surface patches belong to individual ‘seed’ protrusions. Furthermore, the dihedral angle is large between a protrusion and the main cortical cell surface and serves to slow diffusion to cortical areas. We can further intersect with the binary protrusion segmentation from above to guarantee exclusion of cortical surface areas. Lastly, and optionally, we can filter out protrusions that are too small, and close any small holes based on their Cartesian 3D surface area. u-Unwrap3D’s topographic map *S*(*d, u, v*) segmentation approach offers sensitivity and robustness. The initial instance segmentation before filtering detects all potential protrusive features regardless of size, and height. Furthermore, this segmentation strategy is applicable, even when not all protrusions are equiareally represented in *S*(*d, u, v*), as shown by the segmentation of the majority of blebs and filopodia in exemplar cells with suboptimal *S*_ref_(*x, y, z*) (Fig. 6E), and can also be applied in Cartesian 3D (Fig. 2C), removing erroneous segmentations internal to *S*_ref_(*x, y, z*)’s binary volume.

Individual segmented protrusions are genus-0 open surfaces. u-Unwrap3D also implements a harmonic map and equiareal distortion relaxation to map these open surfaces to the 2D plane as a disk and square with aspect ratio optimization to minimize metric distortion (Methods). This allows us to optionally refine the segmentations (Fig. 6D, step 4, Fig. 6F i), for example, by detecting and splitting under-segmented blebs using a gradient-based flow^76^ on the mean curvature in 2D (Fig. 6F ii, Methods). Because of the bidirectionality, the refined segmentation labels can then be transferred back onto the original surface mesh (Fig. 6F i, iii, c.f. before and after refinement, black triangles).

In summary, using u-Unwrap3D to map freely between topographic and Cartesian 3D surfaces and their respective volumetric representations enabled the design of simpler and more generalizable methods to detect and segment surface protrusions in an unsupervised fashion from unconstrained surface geometries. Moreover, any topographic measurements including the topographically segmented protrusions can be bijectively mapped into 2D using topographic cMCF (Movie S3, Methods), an optimal representation for tracking.

### Individual blebs dynamically recruit Septins to local surface regions during retraction

Leveraging the above-described protrusion segmentation, we analyzed the relations between dynamic surface blebbing and the recruitment of Septins. Blebs are globular membrane protrusions of 1-2 μm diameter that are thought to extend in areas of localized membrane detachment from the actin cortex^110,111^. Intracellular pressure expands the budding blebs outward, followed by rapid assembly of a contractile actomyosin network that yields retraction. Cycles of protrusion and retraction have been described to last a few tens of seconds. Associated with the cycles are molecular activities both driving and responding to the morphological dynamics. One such process is the assembly of Septin protein polymers at sites of negative curvature emerging at the bleb necks. Our previous work^38^ has shown that disrupting the bleb cycle diminishes Septin assembly at the cell surface. Here, we reanalyze this phenomenon using u-Unwrap3D to segment and track individual blebs in 2D to quantify Septin accumulation in a mean bleb cycle. We acquired 3D volumes of SEPT6-GFP-expressing MV3 melanoma cells every 1.2s for 200 timepoints. As the cortical cell body exhibits little temporal variation and blebs protrude normally to the surface, the temporal mean surface, ⟨*S*⟩(*x, y, z*), computed by meshing the mean binary volume of individual binary segmentations of *S*(*x, y, z, t*) proxied the cell cortex (Methods). We then apply u-Unwrap3D with *S*_ref_(*x, y, z*) = ⟨*S*⟩(*x, y, z*) to create a single static (*d, u, v*) coordinate space to compute *S*(*d, u, v*) and *S*_ref_(*u, v*) representations for every timepoint. Subsequently, we detect all bleb instances from *S*(*d, u, v*) using the segmentation tools discussed above and mapped the result to *S*(*x, y, z*) for visualization and *S*_ref_(*u, v*) for bleb tracking (Fig. 7A, Movie S7). We further computed mean curvature and normalized Septin intensity surface signals *F*_*i*_(*S*(*x, y, z*)) from Cartesian 3D and mapped these to topographic 3D *F*_*i*_(*S*(*d, u, v*)) for visualization and into 2D, *F*_*i*_(*S*_ref_(*u, v*)), for timeseries analysis (Fig. 7A, Movie S7).

**Figure 7.**
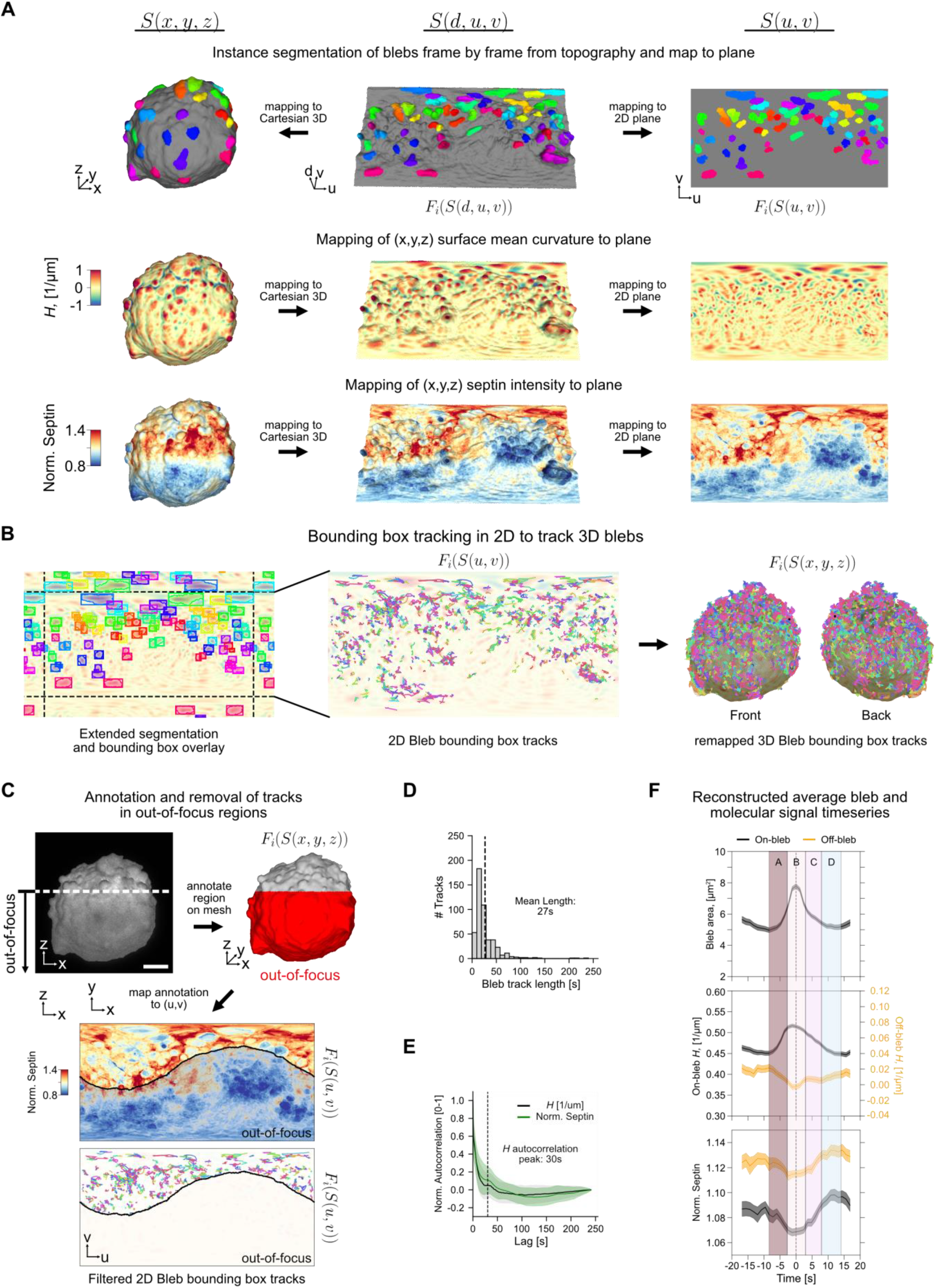
u-Unwrap3D enables tracking and characterization of blebs and associated signals. **A)** Individual blebs segmented in topographic 3D representation are mapped to Cartesian 3D for visualization and to the 2D plane for tracking (top). Individual blebs are uniquely colored. Mean curvature, *H* (middle) and normalized Septin intensities (bottom) are jointly mapped from Cartesian 3D to topographic 3D to the 2D plane. The Septin intensity is normalized at each time to the mean Septin intensity in the whole cell volume to correct for expression variation and photobleaching. **B)** Tracking of individual blebs using an optical flow-guided 2D bounding box tracker. The unwrapped (*u, v*) map is padded on all four sides to capture the continuation of the spherical surface (dashed black lines). Because of the bijectivity between representations individual bleb bounding box tracks in 2D (middle) can be mapped to 3D (right). **C)** Bijective mappings enable the transfer of manually annotated out-of-focus in Cartesian 3D to the unwrapped (*u, v*) 2D plane to restrict intensity timeseries analyses to only the bleb tracks within the in-focus surface regions. The decay in image contrast with sample depth is shown in a maximum projection image on the x-z plane of the first timepoint restricted to the segmented surface ±1 μm (Methods). Scalebar: 10 μm. **D)** Histogram of the in-focus bleb track lengths (dashed line, mean length). **E)** Autocorrelation curves (mean ± standard deviation) of mean curvature, *H* and Septin computed from Cartesian 3D meshes. Dashed black line depicts the lag time of the first autocorrelation side lobe of mean curvature, *H*. **F)**. Average (mean ± s.e.m) time course of bleb surface area (top), mean curvature (on bleb, black; off bleb, orange, and Septin intensity (bottom) over a window 17.5s before to 17.5s after the timepoint of maximum bleb size used for alignment (n=545 bleb events from m=1 cell). A-D labels distinct phases of bleb-mediated curvature Septin recruitment.

We track blebs in 2D *S*_ref_(*u, v*) using the bounding box of individual instances with image padding to account for the spherical boundary conditions (Fig. 7B left). Bleb dynamics were then followed using an established 2D multi-object bounding box tracker^40^ (Fig. 7B middle, Methods). The bidirectional mapping allowed us to map the resulting trajectories (individually colored) to (*x, y, z*) coordinates and generate Cartesian 3D bleb tracks that locate by design to the cell surface (Fig. 7B right, Movie S7). Due to the fast acquisition rate needed to capture bleb dynamics, only a northern portion of the cell could be maintained in-focus. The bidirectional mappings of u-Unwrap3D enabled us to manually annotate an out-of-focus subvolume onto *S*(*x, y, z*) and into *S*_ref_(*u, v*) and retain for the analysis only those bleb tracks within in-focus surface regions (Fig. 7C). The distribution of retained individual bleb track lifetimes showed a peak at 14s and a long tail up to 240s, suggestive of a mixture of short- and long-lived blebs (Fig. 7D). The mean bleb lifetime of 27s corroborates with a 30s periodicity of the mean curvature as inferred by the first peak of the temporal autocorrelation of *H*(*S*(*x, y, z, t*)) in Cartesian 3D (Fig. 7E). The match between the two statistics verify at the population level the accuracy of the single bleb tracking in 2D *S*_ref_(*u, v*).

Next, we computed the temporal autocorrelation of the normalized cell surface Septin signal. The curve strongly aligns with the autocorrelation curve of the mean curvature, suggesting co-fluctuation of the two surface signals. Indeed, we previously showed that surface regions of high Septin intensity with negative surface curvature for at least 30s exhibit a correlation between a negative curvature value and Septin intensity^38^. Whilst the majority of Septin pulses endured only one cycle of bleb formation and retraction, *de novo* formation of stable Septin structures appeared to be driven by several Septin pulses occurring in short succession. We thus hypothesized that these were formed by iterative bleb-driven curvature generation events resulting in local levels of Septin oligomers surpassing a threshold necessary for inter-oligomer polymerization, and enabling stabilization through the formation of higher-order structures^38^. The ability to spatiotemporally track individual blebs enabled us now to quantitatively test this model. For the duration of each tracked bleb, we sampled within its 2D bounding box, distortion-corrected timeseries of bleb surface area, on-/off-bleb surface mean curvature and Septin intensity (Methods). We use the 2D *S*_ref_(*u, v*) bounding box to define a single bleb’s spatial area-of-influence. We consider then the area of *S*(*d, u, v*) enclosed by the box as the bleb’s total 3D surface area and determine ‘on-bleb’ as the largest spatially contiguous region of high mean curvature within the box, and the remainder region as ‘off-bleb’. Low/high mean curvature separation in *S*_ref_(*u, v*) was found by the greater of two thresholds defined by 3-class Otsu thresholding of all *H*(*S*_ref_(*u, v*))(*t*). We used the extracted timeseries to reconstruct the average temporal profile of bleb area, on-/off-mean curvature and Septin intensity in a window of 35s centered on the timepoint of maximum bleb area averaged over 545 individual bleb events collected over 480 bleb tracks in one cell video. The mean timeseries suggests four distinct temporal phases of bleb-mediated Septin recruitment, each ≈ 5s long (Fig.7F, labels A-D). In phase A, the bleb begins expanding, increasing its surface area, accompanied by a sharp increase in on-bleb *H* and a decrease in Septin intensity as the plasma membrane detaches from the actin cortex. The expansion also reduces off-bleb *H*, causing a decrease in Septin intensity off-bleb, presumably due to disrupting Septin structures. In phase B, the bleb reaches maximum size and retracts, decreasing its surface area. Interestingly, unlike *H*, which decreases before the maximum bleb area, the area change is symmetrical, occurring ±2.5s relative to the time of maximum bleb area. Coincident with the bleb increasing to a maximum area, off-bleb *H* decreases to a minimum and Septin intensity both on/off bleb stabilizes at a minimum. As the bleb retracts and the actin cytoskeleton reassembles, off-bleb *H* increases and Septin intensity increases both on/off bleb. In phase C (+2.5s to +8.0s after peak bleb area), the bleb area continues to decrease but at a slower rate. Unexpectedly, the off-bleb negative curvature *H* plateaus at a value lower than its starting value (before phase A) instead of continuing to increase. Concurrently, Septin intensity undergoes the greatest rate of increase on both on-bleb and off-bleb surfaces. At the end of phase C, the Septin intensity is at the levels before phase A (c.f. -15s to -10s). In phase D (+8.0s to +14.0s), bleb area and on/off-bleb *H* recover to pre-expansion levels but Septin intensity continues to increase before plateauing on both on/off-bleb surfaces. Following phase D, the next bleb cycle begins, with similar temporal characteristics to phase A. Altogether these results suggest that blebs generate optimal curvature dynamics for Septin recruitment during the retraction phase. Furthermore, the data supports the model of Septins being recruited to negative curvature patches over multiple bleb cycles, whereby after each bleb formation-retraction a few more oligomers accumulate locally until a threshold concentration is reached to trigger inter-oligomer polymerization of a stable Septin assembly. None of these observations could have been made without the ability of u-Unwrap3D to facilitate segmentation and tracking of blebs, and the accurate conjoint sampling of the local mean curvature and surface-proximal Septin intensity.

## Discussion

Analyzing the spatiotemporal organization of molecular distributions and signaling activities on cell surfaces in 3D has been primarily limited by the lack of methods to measure and track these dynamics in spatiotemporally-consistent timeseries. Recognizing this challenge, we introduce u-Unwrap3D as a surface-guided computing framework for cell biological analyses. u-Unwrap3D enables bidirectional mapping of an arbitrarily complex genus-X Cartesian 3D surface to equivalent surface and volume representations, including the widely-used unit 3D sphere and 2D (*u, v*) representations. A given analytical task may then be solved optimally in one representation and the results transferred to other representations for further analysis or visualization. We demonstrated this principle in four case studies: i) establishment of a retrograde flow of protrusions on natural killer cells in immunological synapse formation with cancer; ii) measurement of information flow to causally link K14+ cells and invasive strands in breast cancer organoids; iii) measurement of ruffle travel speed; and iv) quantifying Septin accumulation within a single bleb cycle.

The broad applicability of u-Unwrap3D mappings, and particular its suitability for spatiotemporal analysis of cell protrusions is founded on three critical developments: i) the computation of a genus-0 *S*_ref_(*x, y, z*) which permits 3D spherical parameterization; ii) robust, bijective, and tunable relaxation of equiareal distortion on the 3D sphere by solving the alternative task of uniformizing the individual face area of the starting conformal parameterization; and iii) a genus-0 shrink-wrap algorithm that can be applied to arbitrarily complex surfaces. Development i) defines a cortical shape to enable explicit isolation from *S*(*x, y, z*) of surface protrusions topographically in *S*(*d, u, v*). Crucially, by mapping *S*_ref_(*x, y, z*) to the fixed geometries of the sphere and plane, we can establish spatiotemporal correspondence across all timepoints of timelapse image sequences. We can thus directly sample spatiotemporally-consistent timeseries of protrusion geometry with surface-proximal molecular activities in other representations. If necessary, the protrusion-less *S*_ref_(*x, y, z, t*) can be readily registered across time using existing methods. Development ii) removes the dependency on the shape properties of *S*_ref_(*x, y, z*) to always ensure the most optimal equiareal spherical parameterization of its mesh (Methods). This is particularly relevant for branched cortical shapes, which can commonly occur with cells embedded in soft microenvironments^40,112^. Similarly, development iii) is fundamental to guaranteeing all u-Unwrap3D representations are computable for arbitrarily complex surfaces *S*(*x, y, z*), with any mesh quality, and by extension to any *S*_ref_(*x, y, z*) to be used.

The concept of geometrical transformation has long existed in mathematics and physics to find simpler methods of equation solving and for plotting shapes, such as the sphere, the cylinder, the Möbius strip, and helicoids amongst others^113^. In computer graphics this is found in the common practice of mapping between surfaces through simpler intermediaries to ensure minimal distortion: for example, the texture mapping of arbitrary surfaces by optimal surface cutting and mapping of the cuts into individual 2D shapes^60^; (*u, v*) parameterization by cutting and gluing individually mapped 2D planar patches^59^; or mapping to canonical shapes such as the triangle^114^, plane^68^ or polyhedra^115^. u-Unwrap3D draws inspiration from this thinking. By developing and making available multiple rationalized 3D-to-2D representations, u-Unwrap3D projects analyses that would otherwise require specialized mathematical operations into a sequence of simpler, computationally tractable procedures for 3D mesh processing, image processing, and machine learning with specific consideration for single cell biology. Unlike computer graphics and developmental biology benchmarks, cell shapes are much more diverse with surface protrusions that are irregular, high-curvature and distributed heterogeneously. To handle this, we chose representations that remap the whole cell surface with well-behaved topologies such as the sphere and plane, and designed a robust conformal map and relaxation scheme to achieve minimal area distortion. This bypasses the numerical instabilities of stitching multiple surface maps and their tracking over time, and the use of solely quasi-conformal mappings that would detrimentally under-represent high-curvature protrusions, the two main techniques used in literature. This also avoids the common cell biology practice of enforcing star-convex approximations of cell shape^13,53^. Second, although we conceived *S*_ref_(*x, y, z*) initially as a mathematical trick to enable spherical parameterization and uv unwrapping, because it is derived explicitly from *S*(*x, y, z*) by considering curvature-minimization, we have found it also serves biologically to decompose *S*(*x, y, z*) as the ‘sum’ of a smooth cell cortex and surface protrusions. Doing so, we have observed that whereas protrusions are primarily transient and short-lived, changes in cell cortex morphology occur much more slowly. Consequently, the u-Unwrap3D introduced *S*_ref_(*x, y, z*) and *S*_topo_(*x, y, z*) could open up new opportunities to identify the different functional roles of surface protrusions and cortex shape in relation to cell state, as well as modelling and quantifying the interplay of dynamic membrane morphology and associated signals with volumetric nuclear/cytoplasmic signalling. Lastly, these new representations introduced by u-Unwrap3D, *S*_ref_(*x, y, z*) and *S*_topo_(*x, y, z*) can be remapped to Cartesian 3D, 3D sphere and 2D plane, which are standard inputs in computer vision and machine learning.

In sum, u-Unwrap3D enables standard computer vision, and machine learning methods be directly applicable to computing tasks on rugged complex surfaces, and for protrusion analysis. More recently, combining different geometric representations helped achieve state-of-the-art performance in addressing complex computational problems such as multiple 2D image views to inform 3D mesh reconstruction^116,117^ and 3D segmentation^76^, or 3D mesh vertex coordinates with 2D unwrapped images for feature extraction^112,118,119^. u-Unwrap3D is fully complementary to these research developments - unifying the common standard representations with cell-biologic representations into a single surface-guided computing framework for manipulating 3D shapes. u-Unwrap3D is made available as a Python package with extensive tutorial. The resources and validation provided by this work will aid the cell biology community to generate testable hypotheses of the spatiotemporal organization and regulation of subcellular geometry and molecular activity.

## Acknowledgments

We thank Hanieh Mazloom Farsibaf and Zach Marin for discussions on mesh processing. Funding for this work in the Danuser lab was provided by the grant R35 GM136428 (NIGMS), and grant RM1 GM145399 (NIH). AW was a fellow of the Jane Coffin Childs Memorial Fund. GMG was a Damon Runyon Fellow (DRG 2422-21). Funding for this work in the Ewald Lab was provided by the NIH/NCI (U01CA284090, U54CA268083, and 3P30CA006973 to AJE), the Breast Cancer Research Foundation (BCRF-24-048; AJE), and the Giovanis Institute for Translational Cell Biology (AJE).

## Author Contributions

FYZ conceived and developed u-Unwrap3D and conducted all analyses. VKW and AJE acquired breast cancer organoid movies of K14+ cells and the invasive organoid shrink-wrapping example. QZ developed a GUI interface to run the u-Unwrap3D workflow. EJ acquired the T cell microvilli timelapse. AW acquired timelapse of the blebbing MV3 cell. GMG, BJC and BC acquired timelapse of the ruffling SU.86.86 cell. MD generated the uShape3D protrusion instance segmentations and surfaces used in bleb, lamellipodia and filopodia examples. FYZ and GD wrote the manuscript with input from all authors. GD provided supervision and obtained funding.

## Data Availability

All data presented herein are available from the corresponding author upon request.

## Code Availability

u-Unwrap3D is open source, developed as a Python package and is available and maintained at: https://github.com/DanuserLab/u-unwrap3D. It is also available in the Python Package Index.

## Methods

### u-Unwrap3D framework

Following we describe the algorithms underpinning each step of u-Unwrap3D depicted in Fig.1, which are grouped into 4 main conceptual steps and 6 specific algorithmic steps.

#### Conceptual step (i): Generating a genus-0 reference surface

##### *Step 1: Derivation of a reference surface, S*_ref_(*x, y, z*) *from an input surface, S*(*x, y, z*)

u-Unwrap3D implements three methods to compute a reference from a given input shape. The first two are top-down approaches based on curvature-minimizing geometric flows. Starting from the input shape, protrusions with intrinsically higher positive curvature are iteratively eroded, yielding increasingly smooth shapes. The third is the reverse bottom-up approach, whereby the input shape is first spectrally decomposed followed by reconstruction of a reference surface with an increasing number of low spatial frequency components. The reconstructed shape is initially skeletal, then smooth, then gradually introduces surface protrusions. Different stopping criteria are provided in u-Unwrap3D to automatically generate candidate reference shapes for the three approaches.

### Top-down curvature minimizing flows

#### Free-form conformalized mean curvature flow (cMCF)

cMCF^63^ modifies the mean curvature flow (MCF) to avoid the formation of pinches and collapsed vertices that compromise bijectivity and cause early flow termination for watertight meshes with high curvature features. MCF evolves Φ_*t*_, the mesh at time *t* according to 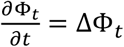, where Δ is the Laplace-Beltrami operator induced by the metric *g* and 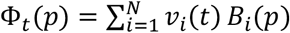 is the discrete mesh parameterization with *N* vertex positions *v*_*i*_(*t*) = {*v*_1_(*t*), …, *v*_*N*_(*t*)} ⊂ ℝ^3^, each *v*_*i*_(*t*) a 3D (*x*_*i*_(*t*), y_*i*_(*t*), *z*_*i*_(*t*)) coordinate tuple and {*B*_1_, …, *B*_*N*_} the local function basis, which for a triangle mesh is the linear hat basis spanned by the edge vectors. Galerkin’s method^120^ is used to find a weak, least-squares solution to the MCF equation within the span of {*B*_*i*_ }, by solving 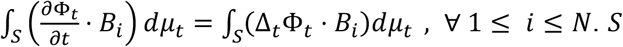 is the surface spanned by {*B*_*i*_} and *dμ*_*t*_ the volume form. The equation is solved to obtain the vertex position, *v*(*t* + *δt*) at the next iteration, *t* + *δt* using backwards Euler integration 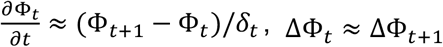 ΔΦ_*t*_ ≈ ΔΦ_*t*+1_ and noting the Laplace-Beltrami is the divergence of the gradient, Δ = ∇ ⋅ ∇ with respect to the local mesh metric *g*_*t*_ to get 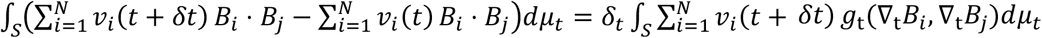 . The integrals, 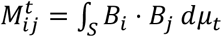 and 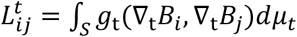 are called the mass (**M**_*t*_) and Laplacian (**L**_*t*_) matrices at time, *t*. Substituting this notation and rearranging, the MCF equation is

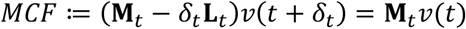

*v*(*t* + *δt*) is then computed from *v*(*t*) by direct matrix inversion. cMCF modifies the MCF equation above by using the Laplacian matrix at time *t* = 0, **L**_0_ for all timepoints. The Laplacian matrix is a measure of stiffness between local mesh faces, see connection to active contour deformation below. Using **L**_0_ for all timepoints instead of recomputing implicitly constrains mesh faces to retain the same aspect ratio and hence to conformalize the flow.

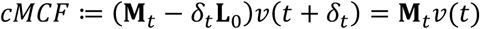

We use the *libigl* library^121^ with the barycentric mass matrix and cotangent Laplacian as the default implementations of **M**_*t*_, **L**_*t*_ respectively. We improve the numerics of solving cMCF by normalization of the surface area and recentering of vertex coordinates at the origin after each iteration as recommended in Alec Jacobson’s blog post (https://www.alecjacobson.com/weblog/?tag=mean-curvature-flow). u-Unwrap3D also implements the robust Laplacian of Sharp et al.^122^ as a drop-in replacement of the cotangent Laplacian in the cMCF equation above i.e. **L**_*t*_= robust Laplacian. The robust Laplacian introduces a mollify factor parameter to enable numerical stability and cMCF iterations to be computed for nonmanifold triangle meshes. The robust Laplacian is essential when applying cMCF to deform topographic meshes (see below) and propagating the (*u, v*) parameterized *S*_ref_(*x, y, z*) . In these two scenarios, the input mesh is nonmanifold or becomes nonmanifold during the iterations. cMCF requires a single-component mesh. u-Unwrap3D retains the largest component with highest number of vertices for an input multi-component mesh. cMCF preserves topology. Therefore for genus-X input shapes the output reference must still be made genus-0. As cMCF is area-minimizing, reducing the size of holes and handles in *S*(*x, y, z*), in some cases, voxelization with morphological operations, and remeshing, described below may be sufficient to generate genus-0 surfaces. Otherwise, genus-0 shrink-wrapping below must be used with isotropic remeshing to guarantee a genus-0 *S*_ref_(*x, y, z*) with equilateral mesh triangulation for equiareal spherical parameterization.

#### Guided curvature-minimizing flow using active contours

A general curvature minimizing flow can be defined by deforming a surface mesh along the gradient, ∇Φ of a specified signed distance transform (SDF, Φ) with stepsize, *δ* set as a function of the magnitude of the local mean curvature, *H* of the SDF, 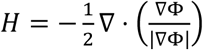 (Fig. S1A iii). u-Unwrap3D uses active contours (see below) to stably deform the mesh. In Fig. S1, for all examples we set *γ* = 1, *α* = 0.1, *β* = 0.5 and the gradient potential, 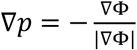 to be the unit length gradient of the Euclidean distance transform, and the stepsize *δ* = percentile normalization of *H*, linearly rescaling its values so that 0.01^st^ and 99.99^th^ percentile are 0 and 1, respectively. This modified flow ensures that the reference shape is always within the volume of the input shape and maximizes erosion of positive curvature surfaces, whilst minimizing changes at negative curvature surfaces. For the bleb cell, this targeted erosion of positively curved protrusions generates a reference surface for bleb height measurement without needing to solve the general ALS baseline estimation with *S*(*d, u, v*) (Fig. 6A-C). The result of guided active contours after postprocessing and cMCF is similar for branched structures (Fig. S1C). Postprocessing was mandatory as the choice of the Euclidean distance transform produces neck pinching singularities akin to that of MCF^63^. Using the diffusion distance transform would alleviate such singularities for this example, giving results closer to that of the cMCF. However, for absolute mitigation of these singularities for arbitrarily complex 3D shapes we have found it necessary to solve the diffusion equation, explicitly specifying the attractor shape and incorporating it as boundary conditions, similar to what we have done previously in 2D^76^ for 3D. The resulting distance transform is also the only one to allow complete extension of *S*_ref_(*u, v*) to map the entire internal volume of *S*_ref_(*x, y, z*) (see Step 6 below).

### Bottom-up shape reconstruction

#### Spectral decomposition of differential operators

Similar to Fourier analysis, the eigenfuction and eigenvalues of a differential operator decomposes an input surface into spatial frequencies. The original shape can be reconstructed incrementally by adding more components from frequencies of increasing contribution. u-Unwrap3D monitors the corresponding rate of decrease in reconstruction error, the CD between the reconstructed with *n* components to the input shape to generate candidate reference shapes. Currently, u-Unwrap3D only implements the analysis with the intrinsic Laplacian-Beltrami operator, which considers the point-to-point distances along the surface. Typically, reference shapes can be efficiently reconstructed using a small number of components (Fig. 1B). This is sufficient for reference shape computation in u-Unwrap3D or performing cellular harmonics^81^ style analysis but is inefficient if reconstructing also the high-frequency surface protrusions. In this case, a more flexible class of differential operators for decomposition are Dirac operators^64,123^ that also account for extrinsic shape properties.

### Stopping criteria for reference shape determination

A natural stopping criteria when using guided curvature-minimizing flow to selectively erode positive curvature protrusions is the iteration number with minimum positive Gaussian curvature. Beyond this, curvature changes reflect only changes to the cortical shape, which also depends on the specified signed distance function. For cMCF and spectral decomposition, the stopping criteria is less obvious. Moreover, for a shape there may be multiple reference shape candidates. u-Unwrap3D provides three stopping criteria that we have found can be adapted for most shapes.

#### Stopping criterion based on the rate of decrease in mean absolute Gaussian curvature

The ideal reference surface *S*_ref_(*x, y, z*) for topographic representation is the cortical cell shape without protrusion morphologies. This corresponds to finding changepoints in the mean absolute Gaussian curvature whereby a decrease is accompanied by a plateau (Fig. 1B) and not the convergence limit of cMCF, which is the sphere^63^. Monitoring the difference in the mean absolute Gaussian curvature over vertices, 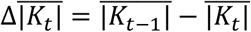 between successive iterations *t* − 1 and *t* provides an automatic stopping criterion, stop_*t*_ = max(*t*_min_, *t*_*K*_). *t*_min_ is a user-specified minimum iteration number and *t*_*K*_ the first iteration for which 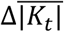 is less than a user-specified threshold, 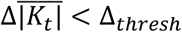 . u-Unwrap3D computes the discrete Gaussian curvature, 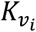 at a vertex *v*_*i*_ for computational speed. It is defined as the vertex’s angular deficit^124^, 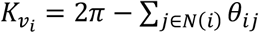 where *N*(*i*) are the triangles incident on *v*_*i*_ and *θ*_*ij*_ is the angle at vertex *i* in triangle *j*.

#### Stopping criterion based on changepoint detection of mean absolute Gaussian curvature

The rate of decrease yields only one stopping iteration suited for simple, pseudospherical cortical shapes with protrusions like the bleb example of Fig. 1B. Moreover, it is sensitive to Δ_*thresh*_, an absolute threshold that may be affected by noise such as mesh quality and protrusion type. Alternatively, u-Unwrap3D computes a baseline of the mean absolute Gaussian curvature curve, and performs changepoint detection, using peak finding to find iteration numbers whose measured mean absolute curvature significantly differs from the estimated baseline. The baseline is computed by symmetrically padded central moving average with window, *w*. The absolute difference between the curve and baseline is used for peak finding using the *find_peaks* in SciPy’s scipy.signal package, specifying a minimum peak height and minimum separation distance between successive peaks. We find this method to be generally robust, with output iteration numbers matching key shape changes, with the first peak shape being a good reference for morphological motifs regardless of the extent of branching in the cortical shape. For tuning, the key parameters is *w* and minimum peak distance. The larger the *w* the smoother the baseline, with the effect of biasing the first detected peak to later iteration numbers. The larger the minimum peak distance, the less the duplication of similar shapes due to noise in the curvature curve. Duplication of similar-looking shapes can also be avoided by increasing the minimum peak threshold.

#### Stopping criterion based on changepoint detection of reconstruction error

For bottom-up spectral decomposition whose shapes do not necessarily change continuously with number of components, monitoring the reconstruction error to the input shape is a more natural choice than mean absolute Gaussian curvature. u-Unwrap3D defines the reconstruction error based on the CD between two point-clouds for faster computation instead of more accurate Wasserstein distances. Changepoints corresponding to salient candidate reference shapes are then detected in the same manner as for mean absolute Gaussian curvature, by the same comparison to baseline and peak finding. However, now, instead of the first stop, we take the last stopping number, to generate the shape closest to *S*(*x, y, z*).

#### Genus-0 shrink-wrapping of a surface mesh

u-Unwrap3D generates a guaranteed genus-0 shrink-wrap of an input mesh *S*(*x, y, z*) by finding an initial genus-0 watertight, manifold mesh and deforming this mesh using active contour along the gradient vector flow (GVF)^67^ of *S*(*x, y, z*) to produce a final tight shrink-wrap (Fig. S2D). If *S*(*x, y, z*) is not watertight or manifold, u-Unwrap3D can use the manifold^125^ algorithm to make it watertight and manifold with minimal changes to compute a GVF. The genus, *g* of a closed orientable surface can be computed from its Euler characteristic, *χ* = 2 − 2*g* = #*V* − #*E* + #*F*. The initial genus-0 mesh is found by applying interval bisection to find the smallest alpha parameter that yields a genus-0 mesh with the alpha wrapping^66^ algorithm implemented in the computational geometry algorithms (CGAL) library. This minimum alpha is lower-bounded by the largest hole or handle in *S*(*x, y, z*). Consequently, this initial shrink-wrap is often too coarse (Fig. S2D,E), but may be optimal when shrink-wrapping reference shapes devoid of surface protrusions. Mesh deformation is performed in epochs and cycles to maintain flow even within narrow concavities and ensure convergence to *S*(*x, y, z*), and to maintain a mesh with a minimum number of vertices. In each epoch, the alpha wrapped mesh was deformed by active contours for *m* = 10 iterations. A cyclic stepsize schedule is used to aid convergence. The stepsize starts at 1 and decays each subsequent iteration by a user specified factor (default=0.5). Should the stepsize fall below a lower bound (default =0.24), then it is reset to 1. With the default parameters, the cycle is 3 iterations. At the end of the epoch, the mesh is decimated to remove short edges, isotropically remeshed, and upsampled by face subdivision to ensure the number of vertices exceeds the minimum. This ‘resetting’ prevents matrix singularity, ensures the mesh is of sufficient resolution, and has even coverage of the GVF of *S*(*x, y, z*) in the next epoch. *n* = 20 epochs constitute a cycle. For shapes devoid of complex surface concavities like *S*_ref_(*x, y, z*) due to the absence of protrusions by design, the initial genus-0 shrink-wrap or one cycle of deformation is sufficient for a good shrink-wrap. Nonconvergence after one cycle is primarily due to flow impedance caused by a ‘bunching together’ of mesh edges in concavities. In short, in these regions, vertices are ‘attracted’ in multiple opposing directions and form a ‘meniscus’ under tension. This occurs primarily when shrink-wrapping *S*(*x, y, z*) with narrow concavities like the filopodia example (Fig. S1C) or the branched organoid example (Fig. S1D). u-unwrap3D solves this by detecting and removing the ‘bunched together’ vertices that impede flow based on their larger distance from *S*(*x, y, z*) and poor triangle quality, defined as twice the ratio of circumcircle and incircle radius. Any resulting open boundary loops after removing vertices are fan-stitched, triangulating boundary vertices to the mean to reform a watertight, manifold, genus-0 mesh. Crucially, the mean point is closer to the valley of the concavity than the original ‘bunching’. Therefore, when a new cycle is run, the resulting shrink-wrap mesh will converge closer to *S*(*x, y, z*). The process of ‘punching-out’ non-convergent areas, stitching and refinement cycle deformations are repeated until the desired shrink-wrap is reached. The CD is computed per epoch, and concatenated across cycles. The mesh with smallest CD to the target *S*(*x, y, z*) or the final iteration mesh is the final shrink-wrap.

#### Mesh voxelization and remeshing

Voxelization converts a surface mesh *S*(*x, y, z*) to a binary volume image *β* where individual voxels are either 1 if they are interior to the surface or 0 if exterior. To do this we create a *X* × *Y* × *Z* voxel volume image larger than the surface with voxels initialised to 0. We then set the intensities of all voxels indexed by the mesh (*x, y, z*) barycenters to 1, i.e. *B*(*x, y, z*) = 1. To ensure a closed binary volume with all interior voxel intensities = 1, *S*(*x, y, z*) is iteratively subdivided by replacing each triangle face by four new faces formed from adding new vertices at the midpoint of every edge until the mean triangle edge length is < 1 voxel. We use the barycenter coordinates of the final mesh *S*_final_(*x, y, z*) to set the binary voxel values. In case of small holes in *S*(*x, y, z*) that would prevent a closed binary volume by binary filling only, *β* is dilated with a ball kernel of *r*_*dilate*_ voxels, binary infilled, then binary eroded with a ball kernel of *r*_*erode*_ voxels. The morphologically processed *B* is meshed with marching cubes^126^ to re-extract a surface mesh with approximately equilateral triangle faces using the isotropic incremental remeshing algorithm^62,127^. This algorithm collapses short edges and splits long edges to a target mean edge length. This remeshing is generally preferred to that of approximated centroidal voronoi diagram (ACVD)^128^ used elsewhere because its implementation within CGAL is more robust and better preserves high curvature protrusions. However, it can be slow for large meshes. A key usage of voxelization is to make *S*_ref_(*x, y, z*) suitable for equiareal spherical parameterization with or without genus-0 shrink-wrapping. In addition to the genus-0 requirement, in practice reference shapes, *S*_ref_(*x, y, z*) with point-like protrusions and narrow necks are not directly amenable to equiareal parameterization by distortion relaxation (see below) as these features impede the diffusion process. To overcome this in u-Unwrap3D, we suggest to construct a dilated version, setting *r*_*dilate*_ ≥ *r*_*erode*_ + 2 and downsampling of the mesh, so that there is sufficient minimal triangle area after conformal sphere parameterization. We note this is not a limitation. If required, GVF and active contours can always be used to transfer the equiareal sphere parameterization of the dilated version to the original *S*_ref_(*x, y, z*). If the input *S*(*x, y, z*) is a smooth shape with only small holes or handles such as the cMCF intermediary *S*_ref_(*x, y, z*) of some simple shapes like Fig. 1B, voxelization with morphological processing and isotropic remeshing may already yield suitable genus-0 *S*_ref_(*x, y, z*) circumventing the need for shrink-wrapping.

#### Conceptual step (ii) Mapping reference surface to unit sphere

A genus-0 *S*_ref_(*x, y, z*) obtained after genus-0 shrink-wrapping and voxelization-based processing is first mapped conformally onto the unit sphere, then area distortion is iteratively diffused to obtain an equiareal parameterization.

##### Step 2: Conformal mapping of reference surface to unit sphere, *S*_ref_(*x, y, z*) to 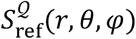 Quasi-conformal spherical parametrization of genus-0 closed surfaces

The uniformization theorem^70^ guarantees the existence of a conformal map onto the unit sphere for genus-0 surfaces but does not specify how to construct an optimal map. u-Unwrap3D implements both i) the direct method of Choi et al.^68,69^ which uses quasi-conformal theory to ensure a bijective spherical parametrization with bounded conformal error (i.e. quasi-conformal) with errors in practice, close to 0, and ii) an iterative Laplacian smoothing approach^129^. For the latter, the mesh vertices are directly projected onto the unit sphere by length normalization, 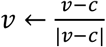 where *c* is the barycentric centroid. Laplacian smoothing is applied using the uniform Laplacian matrix, 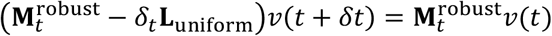 using the mass matrix, 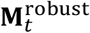 computed by the robust Laplacian approach^122^ at each iteration, *t* followed by reprojection of points to the unit sphere by length normalization. This process iteratively ‘untangles’ the vertices on the sphere. Convergence is monitored by the number of inverted faces relative to the equivalent normal on the sphere. A bijective map is achieved when there are no more inverted faces. Generally, this procedure is fast, converging within 10 iterations even for complex *S*_ref_(*x, y, z*). Non-convergence over 25 iterations, is usually an indication that the input mesh is not genus-0 or contains multiple components. We prefer the second iterative Laplacian smoothing approach. It is faster, more memory efficient and the uniform Laplacian yields better triangle quality for distortion relaxation at the expense of a slight increase in conformal error.

##### Step 3: Equiareal distortion relaxation of conformal sphere paramaterization 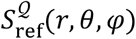 to 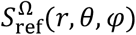

#### Equiareal spherical parameterization by mesh displacement

The vertices of 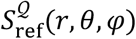 are advected tangentially on the sphere surface, preserving face connectivity to minimise the per face area distortion factor, *λ* with respect to *S*_ref_(*x, y, z*). To do this, the goal is to find the vector field, 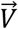 to update each vertex at each iteration. We follow the approach of Lee et al^27^, observing that the partial differential equation for diffusion is 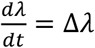 and equal to the infinitesimal change in advecting *λ* along 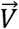, given by the Lie-derivative on 2-forms: 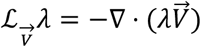 so that 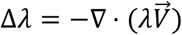. Then, using the fact that the Laplacian is the divergence of the gradient, 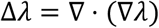, and the chain rule, we have 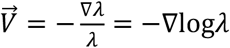. Instead of unwrapping the sphere to the 2D (*u, v*) plane to solve this partial differential equation as in Lee et al^27^, we developed a direct 3D sphere advection scheme that displaces vertices in small constant step sizes *ϵ* by modifying cMCF (see active contour cMCF below), and reprojecting each iteration to the sphere. *λ* and 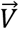 are recalculated for the updated vertex positions and advection is repeated until an equiareal parameterization is achieved or the maximum number of iterations is reached. Details of our advection scheme is given algorithmically.

**Input:** Conformal spherical parameterization mesh, 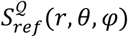 with vertices, *v*_*sphere*_ and faces, *f*_*sphere*_, matching genus-0 mesh, *S*_*ref*_(*x, y, z*) with vertices, *v*_*ref*_ and faces, *f*_*ref*_ with identical number of vertices, |*v*_*sphere*_| = |*v*_*ref*_|, and face connectivity, *f*_*sphere*_ = *f*_*ref*_; vertex step size, *ϵ*; total number of iterations, *T*; mesh stiffness factor, *δ* (also known as the time step, *δ* in cMCF)

**For** iterations *t* = 1,2, …, *T* … do :

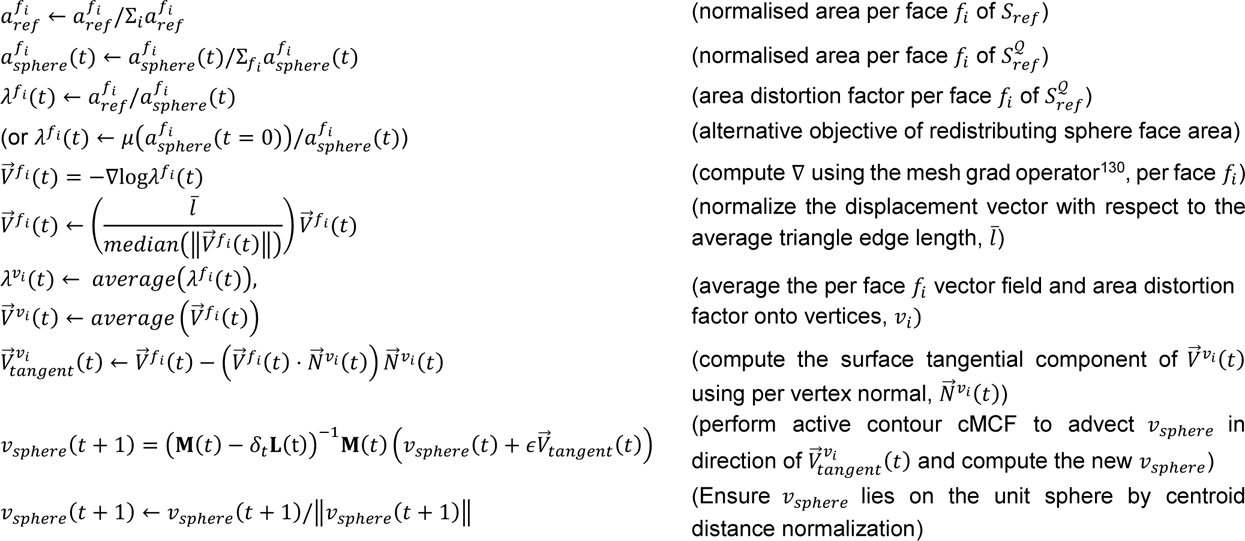

This mesh relaxation bijectively diffuses the area distortion scalar factor on the sphere surface stably until a triangle face degenerates, that is when an interior angle = 0. When this point is reached the distortion error will change sign from monotonically deceasing to fluctuating between increasing and decreasing. We use the first sign change to automatically detect stopping iteration. Unlike conformal parameterization, which comes with mathematical guarantees of existence, an equiareal parameterization is not guaranteed for a general shape. The described relaxation works robustly for the majority of shapes with *δ*-tuning, including achieving the optimal equiareal parameterization for all exemplar shapes shown in this paper. The extent of area relaxation is primarily affected by input mesh quality, which is readily resolved by isotropic remeshing of *S*_ref_(*x, y, z*) but also depends on the reference shape and the number of vertices in its mesh. The following are best practice guidelines we have obvserved: if the relaxation terminates in < 5 iterations, either the conformal is the equiareal parameterization or *S*_ref_(*x, y, z*) has ‘pointy’ and narrow neck shape features that act as diffusion traps to prevent our diffusion-based relaxation, or the mesh is multi-component. To resolve, perform the voxelization processing with larger *r*_*dilate*_. If relaxation cannot decrease to the optimum before triangle collapse, the mesh is too ‘stiff’ because of having too many vertices. To fix this, we downsample *S*_ref_(*x, y, z*) further by specifying a larger edge length in the isotropic incremental remeshing. A non-smooth monotonic decrease in distortion error indicates *δ* is too small. This can also occur for multi-component meshes. A more optimal way to stabilize the relaxation is to remove the dependency on *S*_ref_(*x, y, z*), targeting instead the alternative objective of obtaining uniform distribution of spherical face area, 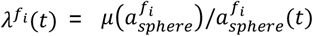 where *μ*(⋅) denotes the mean. Any further algorithm improvements and usage tips will be kept up-to-date on the GitHub repository’s code documentation.

#### Conceptual step (iii) Mapping the 3D unit sphere to the 2D plane

##### Step 4: 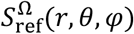 to S_ref_(u, v)

*(u,v)* parameterization involves automatic determination of unwrapping axis, performing the UV map using pullback onto an *N x 2N* aspect ratio image grid with *N* being the user-specified number of pixels, and subsequently finding an optimal scaling *h*, to generate the final *N pixel x hN* pixel *(u,v)* parameterization image with minimal metric distortion.

#### Automatic determination of unwrapping axis using weighted principal components analysis (PCA)

*S*_ref_(*u, v*) severely distorts surface features mapped to the North or South poles of the sphere (Fig. S4A). u-Unwrap3D can automatically find an optimal north-south unwrapping axis to unwrap 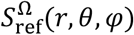 with minimal metric distortion of surface features-of-interest (Fig. S4B). This is done by solving a weighted eigendecomposition problem (Fig. S4B Steps 1-3). Let *v*_*i*_ = (*x*_*i*_, y_*i*_, *z*_*i*_) denote the coordinate of vertex *i* of the spherical parameterization, 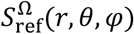 and *w*_*i*_ a vertex weight function, positively scoring the surface feature-of-interest at this vertex and its importance of being mapped with minimum geometrical distortion in *S*_ref_(*u, v*). The 3x3 weighted covariance matrix, **A** = (***w***^*T*^***v***)(***w***^*T*^***v***)^*T*^ over all vertices captures the spread of the weight over the sphere. Eigendecomposition of the symmetric matrix **A** finds the principal orthogonal directions of variance given by eigenvalues ***λ*** = [*λ*_1_, *λ*_2_, *λ*_3_], *λ*_1_ ≥ *λ*_2_ ≥ *λ*_3_ and eigenvectors **e** = [**e**_**1**_, **e**_**2**_, **e**_**3**_]. The eigenvalue captures the concentration of the weight *w* in the direction of the corresponding eigenvector. We adopt the coordinate convention of the north-south sphere axis being the z-axis. Then, the optimal north-south unwrapping axis, is the smallest eigenvector, **e**_**3**_ . To unwrap with respect to **e**_**3**_ we rotate the vertex coordinates *v*_*i*_ so that **e**_**3**_ is the new *z*-axis. As the eigenvector matrix **e** is orthonormal and thus a 3D rotation matrix, **e** is the push forward rotation matrix **R** that maps the 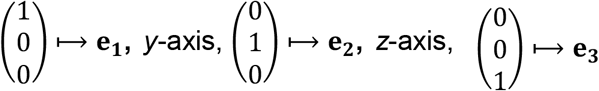,. For a ‘pure’ or proper rotation matrix without reflection the determinant of **R** must be +1, det(**R**) = 1 +1. However eigendecomposition permits det(**e**) = ±1. We construct a proper rotation, 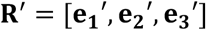 from **R** by flipping the sign of **e**_1_, **e**_2_ to have positive *x*- and y-components respectively; 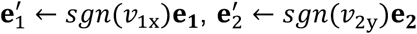 (*sgn* is the sign function) and constructing 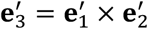 as the cross product of 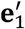 and 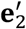. The matrix inverse of **R**^′^ (also the matrix transpose, **R**^′**T**^) is the desired pull back rotation of 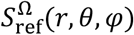 so that 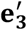 is the new *z*-axis. **R**^′^**T** maps the eigenvectors,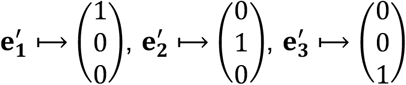. There remains a 180^0^ ambiguity whereby the surface features-of-interest are in the increasing direction of 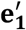 or aligned in the opposite orientation. u-Unwrap3D’s *(u,v)* parameterization centers 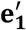 . If in the opposite orientation, this would cause features to map to the edges of the (u,v) image and be split apart. u-Unwrap3D checks for correct orientation by computing the sum of the projected signed weights, *w*′ in the direction of 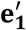, and performing a corrective 180^0^ rotation about 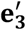 if the sign of the summed projected weights is less than 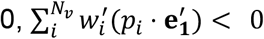 (Fig. S4C Step 4). Crucially, *w*′ should be signed to give a directional preference for the features-of-interest. Examples of suitable *w* are:

1. Magnitude of the mean curvature of *S*_ref_(*x, y, z*) or *S*(*x, y, z*) to prioritize non-flat areas: *w* = |*H*(*S*_ref_(*x, y, z*))| or *w* = |*H*(*S*(*x, y, z*))|
2. Exponential negative magnitude of the mean curvature of *S*_ref_(*x, y, z*) or *S*(*x, y, z*) to proritize flatter areas: 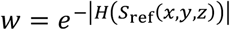 or 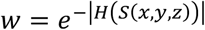
3. Binary mask of positive protrusions from specifying a mean curvature cutoff: *w* = *H*(*S*(*x, y, z*)) > *H*_*cutoff*_
4. *α* - weighted sum of |*H*(*S*(*x, y, z*))| and |*H*(*S*_ref_(*x, y, z*))|, *w* = *α* ⋅ |*H*(*S*(*x, y, z*))| + (1 − *α*) ⋅ |*H*(*S*_ref_(*x, y, z*))| to prioritize non-flat areas of both.
5. *α* - weighted sum of |*H*(*S*(*x, y, z*))| and 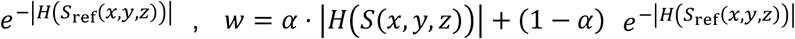, prioritizing protrusions of *S*(*x, y, z*) but flat areas of *S*_ref_(*x, y, z*).

Examples of corresponding signed weights *w*′ for the examples of *w* are:

1. mean curvature of *S*_ref_(*x, y, z*) or *S*(*x, y, z*), specifying preference for positive curvature protrusions: *w*′ = *H*(*S*_ref_(*x, y, z*)) or *w*′ = *H*(*S*(*x, y, z*))
2. *w*^′^ = +1 for flattest surface areas where |*H*(*S*(*x, y, z*))| < |*H*_*cutoff*_|, and *w*^′^ = −1 otherwise
3. *w*^′^ = +1 for *H*(*S*(*x, y, z*)) > *H*_*cutoff*_, and *w*^′^ = −1 otherwise
4. *w*′ = *H*(*S*(*x, y, z*))
5. *w*′ = *H*(*S*(*x, y, z*))

The construction of *w* and *w*′ does not need to be restricted to only geometrical properties of the surface. It can be any surface-associated signal with similar properties. Therefore, we are equally valid in using the fluorescence intensity of a membrane-associated signal.

#### (u,v) parameterization of the unit sphere

We construct an isometric UV unwrap of the unit sphere where *u*, the column image coordinate equidistantly samples the equatorial circumference of the sphere, a total length 2*π* and *v*, the row coordinate equidistantly samples the great arc distance from north to south pole, a total length *π*. This specifies a *N* × 2*N* pixel UV image with *N* as a user-defined size. By default *N* = 256 pixels. The UV mapping is constructed by pullback. Let *u* = *θ* be the azimuthal and *v* = *φ* be the inclination angles of the sphere and setup the *N* × 2*N* grid of *v* vs *u* over the parameter space [−*π*, 0] × [−*π, π*]. Convert the spherical coordinates, (*r* = 1, *θ, φ*) to cartesian coordinates, (*x, y, z*) = (sin*θ* cos *φ*, sin *θ* sin *φ*, cos *θ*). Each (*x, y, z*) coordinate is matched by nearest normal distance to a triangle face, *ABC* of 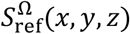 to compute barycentric coordinates giving (*x, y, z*) as a convex combination of the vertices *A, B, C*; *μ*_*A*_*A* + *μ*_*B*_*B* + *μ*_*C*_*C, μ*_*A*_, *μ*_*B*_, *μ*_*C*_ ≥ 0 and *μ*_*A*_ + *μ*_*B*_ + *μ*_*C*_ = 1 . By bijectivity of *S*_ref_(*x, y, z*) and 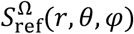, we set *A, B, C* to the respective *S*_ref_(*x, y, z*) vertex coordinates to produce the respective uv-coordinate mapping, (*u, v*) ↔ *μ*_*A*_*A* + *μ*_*B*_*B* + *μ*_*C*_*C* . Note, setting *A, B, C* to be the vertex coordinates of any mesh bijective to 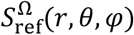 e.g. *S*(*x, y, z*) in the direct unwrapping case produces similarly the corresponding uv-coordinate mapping for that surface. The weights *μ*_*A*_, *μ*_*B*_, *μ*_*C*_ can be used to interpolate scalar vertex associated quantities from the sphere, 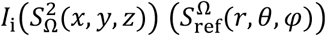 to the 2D plane, *I*_i_(*S*_ref_(*u, v*)) such as mean curvature, *H* so that 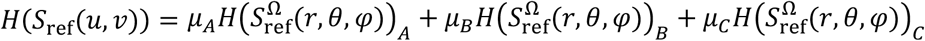 . Instead of the vertex coordinates of *A, B, C* we use the values of the scalar measurement associated with the vertices *A, B, C* replaced now by scalar values (see transfer of measurements between meshes below).

#### Optimizing the aspect ratio of (u,v) parameterization

We find an optimal scaling parameter, *h* of the horizontal *u* axis to minimize metric distortion. The resulting UV map is of pixel size (*N, hN*). This is necessary because a (*N*, 2*N*) mapping would severely distort the major length axis of an ellipsoid if this corresponded to the *v* axis. Finding a scaling parameter solves this issue without manual alignment of the ellipsoid. u-Unwrap3D implements two metric distortion objectives: length ratio and the Beltrami coefficient (Fig. S4G). Length ratio defines the scaling to be the ratio of the mean total Cartesian distance traversed horizontally in increasing *u*-direction over the mean total Cartesian distance traversed vertically in increasing *v*-direction, 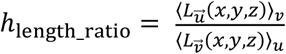. This definition naturally extends the distance ratio of the North-South great arc to equatorial circumference for the sphere to a general *(u,v)* parameterization of a closed surface. The Beltrami coefficient measures the conformal error^68^, the primary length-stretching error being minimized when optimizing for the aspect ratio (Fig. S4G iii). u-Unwrap3D finds the scaling parameter, *h*_*beltrami*_ minimizing the Beltrami coefficient over a given range of *h* using Brent’s method of bounded minimization implemented in Scipy, scipy.optimize.fminbound. By default, this range is *h* ∈ [0.1, 5] inclusive. For closed ellipsoidal-like 3D surfaces, length ratio is faster to compute and can yield qualitatively better-looking results (Fig. S4G ii). Beltrami coefficient, however, is universally applicable for closed and open surfaces. Thus, it can be used to optimize the aspect ratio of the square parameterization for protrusion segmentation refinement.

#### Conceptual step (iv) Mapping the proximal volume with respect to reference surface

##### Step 5: V(x, y, z) to V(d, u, v)

*Topographic coordinate space* (*d, u, v*) *construction*. UV-unwrapping establishes bijection between a 2D uv plane and a 3D Cartesian surface, (*u, v*) ⟷ *S*_ref_(*x, y, z*) . We can also map the local volume around *S*_ref_(*x, y, z*) by constructing a topographic (*d, u, v*) coordinate space, *V*(*d, u, v*) corresponding to a volume space normal to *S*_ref_(*x, y, z*). u-Unwrap3D does this by propagating the (*u, v*) parameterized *S*_ref_(*x, y, z*) to sample the Cartesian 3D space at equidistant steps of *α* voxels, referred to as *α* -steps, along the steepest gradient of the signed distance function of 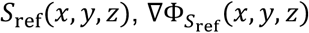 for a total of *D* steps, *d* ∈ −*D*_*in*_, …, *D*_*out*_ where *D* = *D*_*out*_ + *D*_*in*_ is the total number of *α*-steps outwards and inwards with respect to *S*_ref_(*x, y, z*) (by definition, *d* = 0). *D*_*out*_ can be automatically determined by u-Unwrap3D to ensure *V*(*d, u, v*) fully encapsulates *S*(*x, y, z*) by measuring the Hausdorff distance between *S*_ref_(*x, y, z*) and *S*(*x, y, z*) . *D*_*in*_ is primarily user-defined for computational efficiency or can be automatically determined as a fraction of the maximum internal distance transform. The signed distance function Φ(*x, y, z*) of *S*_ref_(*x, y, z*) is computed from the binary volume representation of *S*_ref_(*x, y, z*). If the binary was not generated as part of Step 2 to find a reference shape, then we voxelize *S*_ref_(*x, y, z*) (described above). Active contours (see below) is used to propagate the (*u, v*) parameterized *S*_ref_(*x, y, z*) along the gradient of the distance transform to sample the internal volume. A forward Euler scheme is used for propagating outward; 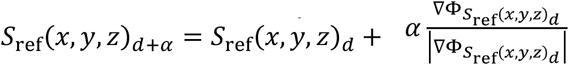 at *α* voxels from *S*_ref_ (*x, y, z*)_*d*_ and 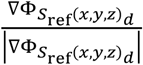 is the unit gradient of 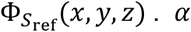 is automatically determined as the average of the mean Cartesian length a *u* and *v* pixel represents. For a large *D*_*out*_, as the intra-spacing of sampled 3D (*x, y, z*) positions increases (Fig. S3A), numerical instabilities arise with active contours, requiring implicit mesh smoothing methods, which can be slow for large meshes. The forward Euler scheme is faster and stable, leveraging efficient 1D uniform filtering along *u*-, and *v*-axes with window size set as a fraction of the *u* size, typically 1/8, to suppress instabilities. By default, u-Unwrap3D uses the Euclidean distance transform (EDT) for external propagation and the Poisson distance transform^75^ for internal propagation. Similar to the idea of cMCF, when *S*_ref_(*x, y, z*) is non-convex and especially when it is branched, the piecewise-continuous EDT gradient develops singularities with propagation biased towards and converging onto the medial-axis skeleton. Consequently, EDT does not sample the internal volume as uniformly as the inherently smooth gradients of the diffusion-based Poisson distance transform^63^. u-Unwrap3D also allows for topographic space construction with custom-defined external gradients. Notably, u-Unwrap3D implements the 3D equivalent of the explicit attractor diffusion transform used in u-Segment3D for universal shape representation^76^. This allows the flexible construction of custom gradients to completely sample the internal volume bijectively (e.g., specifying a smooth medial-axis skeletal shape as the attractor); and most generally, the composition of volumes between multiple reference surfaces. As a further complement to this feature, u-Unwrap3D also implements functions to automatically find the individual stopping iteration when each vertex first hits the attractor surface with constant *α*-step propagation. u-Unwrap3D can then resample the trajectory of each vertex from iteration 0 to its stop iteration so that all vertices hit the attractor surface simultaneously, in the same number of *D*_*in*_ steps. Generally, as u-Unwrap3D is provided as a package, users can specify variable *α* -step size and custom functions to construct alternative adaptively-sampled topography (*d, u, v*) spaces. Lastly, trilinear interpolation of the respective Cartesian volumetric signal intensities, *I*_*i*_(*V*(*x, y, z*)) at the (*x, y, z*) coordinates indexed by *V*(*d, u, v*) generates the topographic 3D equivalents, *I*_*i*_(*V*(*d, u, v*)).

##### Step 6: I_i_(V(d, u, v)) to S(d, u, v)

#### Topographic mesh S(d, u, v) construction

The binary volume representation of *S*(*x, y, z*), *I*_*i*_(*V*(*x, y, z*)) is mapped by interpolation into the topography (*d, u, v*) space to obtain its topographic equivalent binary, *I*_*i*_(*V*(*d, u, v*)). Marching cubes at isovalue = 0.5 are then used to extract an initial *S*(*d, u, v*) and remeshed with ACVD or isotropic incremental remeshing to obtain the final *S*(*d, u, v*) mesh with near-equilateral triangle faces. To construct the corresponding Cartesian 3D mesh, *S*_topo_(*x, y, z*) of *S*(*d, u, v*), we use the same face connectivity and interpolate the (*x, y, z*) vertices indexed by the (*d, u, v*) vertices. To transform surface signals, *F*_*i*_(*S*(*x, y, z*)) to *F*_*i*_(*S*(*d, u, v*)), we match *S*_topo_(*x, y, z*) vertices to the faces of *S*(*x, y, z*) by nearest normal surface distance and perform barycentric interpolation for scalar functions or take the mode for labels (see below).

### Transfer of measurements between two meshes

We assume the convention that all measurements are vertex-based and both the source and target meshes are injective onto the same shape, primarily because they are two different parameterizations. Examples of mesh pairs occurring in the main u-Unwrap3D workflow that require transfer of measurements are: 1) cMCF *S*_ref_(*x, y, z*) and genus-0 shrink-wrapped 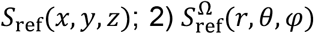 and (*u, v*) parameterized unit sphere; 3) *S*(*x, y, z*) and *S*_topo_(*x, y, z*). All measurements associated with the source mesh are considered to be either (i) scalar, taking any real value or (ii) categorical such as segmentation labels, which have semantic but no ordinal meaning. Thus, mean curvature represents a single scalar measurement, but vectorial measurements like (*x, y, z*) coordinates or (*r, g, b*) coloring represent three independent scalar measurements. Each vertex of the target mesh is matched to the nearest face in the source mesh using the *trimesh*.*proximity*.*closest_point* function in the Trimesh library. This function finds the closest vertex in the source to the queried target vertex, the distance of which is used to construct an axis aligned bounding box (AABB) centered around the query point. Then, the nearest source face intersecting the AABB is found. The barycentric coordinates of the point (*μ*_*A*_, *μ*_*B*_, *μ*_*C*_) with respect to the source face vertices, *A, B, C* where *μ*_*A*_*A* + *μ*_*B*_*B* + *μ*_*C*_*C, μ*_*A*_, *μ*_*B*_, *μ*_*C*_ ≥ 0 and *μ*_*A*_ + *μ*_*B*_ + *μ*_*C*_ = 1 is found after projecting the target point onto the source face using the source face normal with the Trimesh function, *trimesh*.*triangles*.*points_to_barycentric*. Scalar measurements at the face vertices, 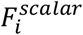 are interpolated to obtain their value at the target vertex, 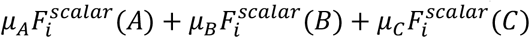. For categorical labels the value at the target vertex is computed using the mode, 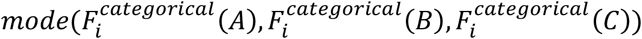. The interpolation or mode is performed for each scalar or categorical measurement *F*_*i*_ to be transferred from the source to the target mesh.

### Temporal application of u-Unwrap3D

For timeseries application, we assume that videos have been globally registered between timepoints to correct for rotation, translation and possibly shear transformations. If they are not, u-Unwrap3D provides scripts using the SimpleITK library. u-Unwrap3D is then applied individually to each timepoint to obtain the different representations as illustrated in Fig. 3A. There are several additional considerations involving parameter setting for timeseries analysis.

#### Individual timepoint reference shape, S_ref_(x, y, z, t) . S_ref_(x, y, z, t)

should change smoothly between timepoints. To ensure this, we can either: 1) derive a temporally averaged mean absolute Gaussian curvature or reconstruction error curve to apply the same single stopping number, or 2) apply a moving average filter to temporally smooth the individual stopping numbers inferred in each timepoint. Approach 2) is more general, capable of handling changes in protrusion morphological properties. We used 1) for all examples. It works when protrusions maintain similar morphological properties across timepoints like height and width.

#### Optimal uv-unwrapping axis

Different timepoints may yield a slightly different unwrapping axis, affecting the temporal consistency of representations. We choose a single reference timepoint to determine the optimal unwrapping, then apply this to uv-map the spherical parameterizations of all other timepoints. Examples of reference timepoints are the first, middle, last timepoints, or the timepoint when a particular event occurs, e.g. the point of stable synapse formation between the NK and cancer cell example, (Fig. 4A).

#### Optimal uv-map aspect ratio

The individual timepoint reference shape is also used to determine the optimal uv-map aspect ratio to apply to all timepoints.

#### Setting topography space α-stepsize and D total steps

We use the same single timepoint reference to set the *α-*stepsize to the average of the mean Cartesian distance a *u* and *v* pixel represents. Alternatively, the *α-*stepsize can also be manually set. Importantly, α-stepsize should be the same for all timepoints. We want all timepoints to map into the same (*d, u, v*) volume. Therefore, the total number of *D* steps outside and inside of *S*_ref_(*x, y, z, t*) should be the same for all timepoints. *D*_*out*_ can be automatically inferred by using the mesh corresponding to the maximum projection of the binary volumes of all timepoints.

#### Non-rigid registration prior to or after u-Unwrap3D

By mapping shapes into canonical spherical, (*u, v*), and topographic coordinates, u-Unwrap3D establishes automatically spatiotemporal correspondence across all timepoints, regardless of the shape complexity. These mappings allow powerful timeseries analysis yet they do not guarantee perfect alignment of individual surface features. For most cases, however, under the assumption of global video registration prior to unwrapping and suitably derived *S*_ref_(*x, y, z, t*), we find that individual features are sufficiently well aligned with only minor jittering in (*u, v*).Hence tracking problems can be solved by tracking segmented features (Fig. 6) or optical flow approaches (Fig. 4,7). Thanks to the modular nature of u-Unwrap3D, non-rigid spatiotemporal registration techniques can be readily incorporated to improve alignment. u-Unwrap3D implements the demons family of volumetric registration (demons^131^, diffeomorphic^80^, and symmetric^79^) through SimpleITK which can be applied to register the video before processing or register volumetric representations of meshes as we did in mapping all *S*_ref_(*x, y, z, t*) to the mean temporal reference, ⟨*S*_ref_⟩(*x, y, z*) (Fig. 4B). *ANTspy* provides further advanced volumetric registration methods. u-Unwrap3D also implements gradient vector flow (GVF) and active contour deformation, which can be used to register shape 1 to shape 2, by propagating shape 1 along the GVF of shape 2. GVF is most effective when used with the volume registration predicted shape 2 from shape 1, to correct for remaining misregistered high-curvature surface regions. u-Unwrap3D also provides a fast *k* nearest neighbor matching through the *point-cloud-utils* library, but this approach, unlike GVF, can only match two convex surfaces. Finally, methods have been developed to directly register in spherical^43,132,133^ and (*u, v*) parameterizations^134-136^ with proper handling of the domain boundary conditions. Such methods could be added to future versions of u-Unwrap3D, if necessary.

### Mesh displacement by active contours

Active contours, or ‘snakes’ ^137^, define the boundary of an image region by minimizing its contour energy, *E*. The contour energy is the sum of an internal, *E*_*int*_ and an external energy, *E*_*image*_, *E*(*v, I*) = *E*_*image*_(*v, I*) + *E*_*int*_(*v*). The internal energy is set by *E*_*int*_ = ∫ *α*|*v*^′^|^2^ + *β*|*v*^′′^|^2^ *ds*, where the number of ′ denotes the order of the spatial derivative. Here, the first term denotes tension with *α* defining the elasticity of the contour. The second term denotes the bending stiffness with *β* defining the bending rigidity of the boundary, i.e. a contour in 2D and surface mesh in 3D. The external energy is set by *E*_*image*_ = −∫ *p ds*, where *p* is an attractor image for the contour. Minimizing *E* is equivalent to solving the Euler-Lagrange equation, *αv*^′′^ − *βv*^′′′′^ = −∇*p* or in operator form, a linear equation, **A***v* = −∇*p*, with **A** representing the second and fourth order derivative operators by finite differences. Given a mesh with vertex positions *v*(*t*), its next position, *v*(*t* + 1) is found by minimizing the residual error **A***v* + ∇*p* using gradient descent and backwards Euler, giving the update, *v*(*t* + 1) = *v*(*t*) − (**A**_*t*_*v*(*t* + 1) + ∇*p*) and the linear system to be solved for, (**I** + **A**_*t*_)*v*(*t* + 1) = *v*(*t*) + ∇*p*, where **I** is the identity matrix. For a mesh, the second order and fourth order derivatives are the Laplacian, **L**_*t*_ and the bilaplacian, 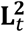 operators. The final equation is 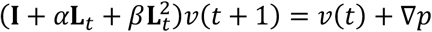. To evolve mesh vertices, *v*(*t*), we set *p* = Φ, the signed distance transform or GVF in the case of shrink-wrapping, and introduce a factor, *δ* representing the stepsize, giving 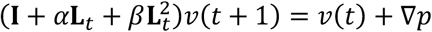. u-Unwrap3D solves this Ax=b style equation by LU decomposition (Scipy *scipy*.*sparse*.*linalg*.*spsolve* function) using the cotangent Laplacian by default. We adopt the convention that the gradient of Φ, the signed distance transform orients inwards to the cell center. Consequently, a positive *δ* > 0 deforms a reference mesh inwards, and a negative *δ* < 0 moves the mesh outwards.

### Mesh displacement by active contour cMCF

If *β* = 0 in the active contours equation above, we have the second-order equation, (**I** + *α*_*t*_**L**_*t*_)*v*(*t* + 1) = *v*(*t*) + ∇*p*. This is the MCF update without the mass matrix, **M**_*t*_. We reintroduce **M**_*t*_, replacing **L**_*t*_ by 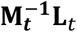 to get (**M**_*t*_ − *α*_*t*_**L**_*t*_)*v*(*t* + 1) = **M**_***t***_(*v*(*t*) + ∇*p*). Thus, the tension *α*_*t*_ plays the same role as the time-step, *δ*_*t*_ . Setting **L**_*t*_ = **L**_**0**_ determines the cMCF deformation in response to an external force, ∇*p*. As for active contours, this Ax=b style equation is solved by direct matrix inversion to deform surface meshes. In this paper, we refer to this as *active contour cMCF* and evolve mesh vertices with respect to the gradient of Φ in the same manner *p* = Φ. u-Unwrap3D primarily uses active contour cMCF for relaxing the equiareal distortion on the sphere, and in the construction of the topographic cMCF below. For general mesh deformation, active contours typically solves faster and yields smoother solutions.

### A topographic conformalized mean curvature flow (cMCF) for flattening topographic surfaces, *S*(*d, u, v*)

*S*(*d, u, v*) are open surfaces. Application of cMCF^63^, formulated for closed surfaces, maps *S*(*d, u, v*) onto the 2D plane as an elliptical disk and in the limit, to a point. We instead want to flatten topographic surfaces to converge to a planar (*u, v*) rectangle. This allows tasks such as the tracking of protrusion segmentations be performed in 2D with the result mapped back to the topographic surface and beyond. To achieve this, we modify cMCF to impose no-flux constraints in the *u*-, *v*-directions on boundary vertices, ∂*S*, and allow flow only in the depth, *d* direction by adding counteracting gradients in the *u*-, *v*-directions:

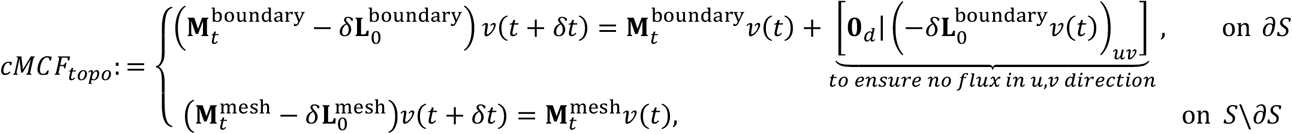

The flow for interior vertices still follow the standard cMCF formulation, using 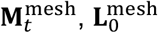 as the mass and Laplacian matrices defined for a 3D triangular mesh (see above). On the boundary, ∂*S* we use the mass, 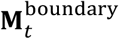 and Laplacian, 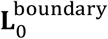 matrix defined for a 2D line to impose no-flux conditions and [**A**|**B**] denotes the augmented matrix formed by appending the columns of matrix **A** and **B**. As with cMCF, we solve for the vertex position at the next timepoint *v*(*t* + *δt*) by direct matrix inversion. *δ* controls the time-step. The larger *δ* is the faster the convergence (Fig. S6, Suppl. Movie 6). Under the described topographic cMCF, the surface *S*(*d, u, v*) converges to a planar surface with uniformly constant *d, S*(*d* = *constant, u, v*), which by bijection can be resampled to a (*u, v*) image grid. Associated measurements on the surface topography are correspondingly transferred to 2D image representations.

### Transfer of protrusions from one core shape to another using canonical coordinate systems

The unit sphere, (*u, v*) plane and (*d, u, v*) volume represent canonical coordinate systems that are invariant. As such, shapes mapped into any one of these coordinates are automatically put in bijection with each other, their original geometry being effectively integrated out. This is why applying u-Unwrap3D frame-by-frame automatically spatiotemporally registers surfaces (Fig. 3). A consequence of this property is the ability to decouple measurements and geometry associated with one shape and transfer it to another. By establishing a reference surface, *S*_ref_(*x, y, z, t*), and mapping this to canonical coordinates, u-Unwrap3D explicitly decouples protrusions from the reference, allowing its transfer by property (texture transfer) or geometry (physical transfer) to other shapes for analysis.

#### Physical transfer of geometry

This is most easily performed in topography space, which encodes the relative geometry of protrusions in a single scalar, *d* describing the relative curvilinear distance to S_ref_(*x, y, z*) (Fig. 3B). We perform first any necessary array resizing and translation to match the *d* = 0 in the topography space of the source 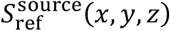 to the *d* = 0 in the topography space of the target 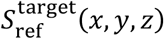. The transfer operation then simply involves embedding the source protrusions either as a topographic mesh or topographic binary volume into the aligned target topography space so that 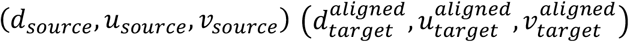 and in bijection with the Cartesian space of (*x*_*target*_, y_*target*_, *z*_*target*_). Transferring in spherical or (*u, v*) parameterization is more challenging. The distances traversed from S^source^(*x, y, z*) to 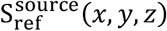 are measured, 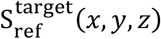 is matched to 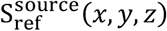 through their spherical or (*u, v*) parameterization to transfer the distances to 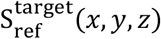. Finally, the distances need to be reversed through a flow that reverses the cMCF or active contour process to reconstruct the source protrusion geometry on 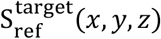, relative to its signed distance transform.

#### Texture transfer of property

For many analyses, such as tracking or comparing morphological statistics, only a measurement-of-interest representing the protrusion needs to be transferred not the actual geometry. For example, in analyzing the NK and cancer cell interaction in Fig. 3, we transfer the mean curvature, *H*(*S*(*x, y, z, t*)) to track protrusions in ⟨*S*_ref_⟩(*u, v*), the (*u, v*) parameterization of the temporal average reference shape. Since the protrusions are captured as a measurement, the transfer is done as described above for measurement transfer between two meshes. If the two shapes are not in correspondence, they can either be mapped to the same canonical coordinate space or be registered using appropriate surface or volumetric registration techniques (see above in time application).

### Quantification of geometric deformation errors for meshes

Surface mappings do not conserve local geometrical measures like angles, edge lengths and face area. Quantification of the distortion in these measures enables a task-specific optimization of the mapping and correction of statistical measurements made on the mapped surface. There are two primary geometric distortions to quantify: conformal and area distortion error (Fig. S3). An isometric deformation is one with no error, i.e., both conformal and area distortion errors are 0.

#### Conformal error

The conformal or quasi-conformal error, 𝒬_*i*_ measures the extent the shape of a mesh element *i*, e.g. a triangle face, is stretched. It is 0 if the relative distances between vertices and the angles between edges are preserved after the mapping. We compute 𝒬 of mapping a 3D surface triangle Δ*ABC* to Δ*DEF* by first reparameterizing the triangles equivalently in 2D coordinates. Let *A* = (*x*_1_, y_1_, *z*_1_), *B* = (*x*_2_, y_2_, *z*_2_), *C* = (*x*_3_, y_3_, *z*_3_) with edge vectors,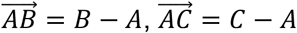, then an identical 2D triangle Δ*A*^′^*B*^′^*C*^′^ is given by 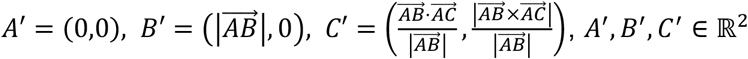. Let 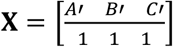, be the 3x3 homogeneous vertex coordinates of Δ*A*^′^*B*^′^*C*^′^ and 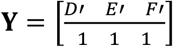, the 3x3 homogeneous vertex coordinates of Δ*D*^′^*E*^′^*F*^′^ then we solve for the 3x3 matrix, **A** that maps **X** to **Y** = **AX. A** is affine and of the form 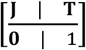 where **J** is a 2x2 transformation matrix and **T** a translation matrix. Eigenvector decomposition of **J**^**T**^**J** gives 2 eigenvalues *λ, λ, λ* < *λ* and the singular values of 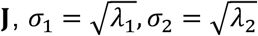 . The ratio 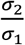 is the conformal error^138^. The global conformal error, 𝒬 of deforming a surface mesh *S*_1_ to a mesh *S*_2_ is the area weighted average of individual conformal errors 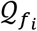 of each triangle face *f* in 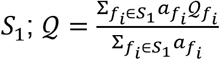 where 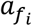 is the area of face *f*_*i*_ of *S*_1_.

#### Beltrami Coefficient

The Beltrami coefficient^68,114^ is another measure of the conformal error of a mapping from the theory of quasiconformal maps. It is less intuitive to interpret than the conformal error but faster to compute, therefore we use it only to find the optimal aspect ratio for (*u, v*) parameterization.

#### Area distortion error

The area distortion error, *λ* measures the extent the surface area fraction of a mesh face is preserved during a surface mapping. One measure of *λ* is σ_1_σ_2_, the product of the singular values of **J** and the area of the distortion ellipse^139^. Here we use the surface area fraction ratio, 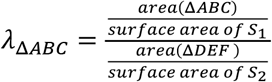 as a direct measurement of the area distortion in mapping Δ*ABC* to Δ*DEF* in 3D. The global area distortion error,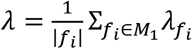 for mapping a mesh *S*_1_ to a mesh *S*_2_ is the mean over all individual area distortion 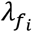 of each triangle face *f* in *S*, with |*f*_i_ | denoting the number of faces in *S*_1_ and 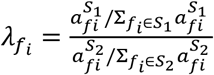 denoting the area distortion of face *f*_*i*_ . The normalization of face area by total surface area is crucial to enable the evaluation of *λ* independent of size.

### Quantification of geometric deformation error for UV images

UV mapping defines a bijective relation between the 2D (*u, v*) rectilinear grid and a 3D surface, *S*(*u, v*) ↔ *S* = *S*(*x*(*u, v*), y(*u, v*), *z*(*u, v*)). Differentials can be used to compute geometric quantities when the (*u, v*) spacing is comparable to the (*x, y, z*) spacing. The differential area of a (*u, v*) pixel is 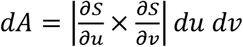, where 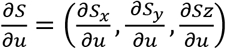 and 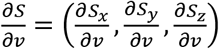 are the gradients of the *x, y, z* surface coordinates in *u, v* directions. The topographic space construction establishes bijection of the 3D (*d, u, v*) volumetric grid to a 3D volume, *V*, (*u, v, d*) ↔ *V* = *V*(*x*(*u, v, d*), y(*u, v, d*), *z*(*u, v, d*)). The differential volume of a (*d, u, v*) voxel is 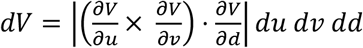 . The matrix 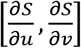 is the 2x3 Jacobian matrix, **J** and the conformal error per pixel is 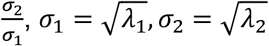 where *λ*_1_, *λ*_2_, *λ*_1_ < *λ*_2_ are the two eigenvalues of **J**^**T**^**J**. The global conformal error 𝒬 of *uv*-mapping the surface mesh *S* is the differential area weighted average of individual conformal errors 𝒬 _*uv*_ of each *uv* pixel; 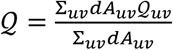 where *dA* _*uv*_ is the area element of the *uv* pixel. The area distortion error per *uv* pixel is the ratio between the surface area fraction of a *uv* pixel and the corresponding surface element on 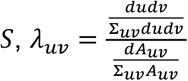 . Note *dudv* = 1 and Σ *dudv* = total number of *uv* pixels. The global area distortion error, 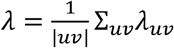 for *uv* mapping a surface *S* is the mean over all individual area distortion *λ*_*uv*_ of each *uv* pixel.

#### Stopping criteria for area distortion relaxation of S^𝒬^(r, θ, φ)

The iterative area distortion relaxation allows for intermediate parameterizations between fully conformal, *S*^𝒬^(*r, θ, φ*) and fully equiareal, *S*^Ω^(*r, θ, φ*). The two extremes are not always optimal. For example, for branched shapes, conformal maps insufficiently represent the branching, but equiareal maps overemphasize the branching, distorting the representation of surface elements of *S*_ref_(*x, y, z*) outside the branching zone too much. In these cases, aside from specifying a target equiareal distortion value, we can construct a parameterized isometric loss function, 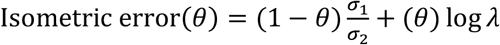, with a weight *θ*, adjusting the relative importance between 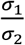, the conformal error, and log *λ*, the log of the area distortion error whose minimum value sets the stopping iteration. *θ* = 0.5 balances both error types.

### Assessment of geometrical difference between two meshes

Four metrics were used to assess the difference between two meshes *S*_1_ and *S*_2_ possessing different number of vertices and faces in Fig. S3, S5; chamfer distance (CD), Wasserstein-1 distance (*W*_1_), the difference in surface area (Δ*A*) and the difference in volume (Δ*V*). CD is the mean Euclidean distance between all vertices of *S*_1_ when matched to the nearest vertex of *S*_2_ and vice versa, 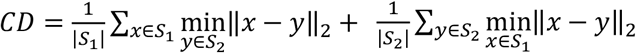 . The Wasserstein-1 distance (*W*_1_) or Earth-mover’s distance (EMD) is the minimum total area weighted distance of 1-to-1 matching vertices on *S*_1_ and *S*_2_. *W*_1_ accounts for the area of triangle faces and is minimal if the vertices of *S*_1_ is a uniform sampling of *S*_2_ or vice versa. Exact computation of *W*_1_ is impractical, even for small meshes. We compute *W*_1_ using the sliced-Wasserstein, *SW*_1_ approximation, which uses random spherical projections to sum multiple 1D EMD distances^140^. Specifically we use the *ot*.*sliced*.*max_sliced_wasserstein_distance* function from the Python *POT* library with 50 projections and average the result from 10 evaluations to report an estimate. The difference in total surface area is 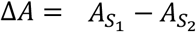 and is 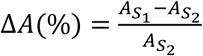 when given as a percentage. Total surface area was computed as the sum of individual triangle areas. The difference in total volume is 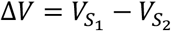 and 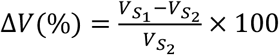 defines the relative volume difference as a percentage. The volume of a mesh was computed as the number of voxels in its binary voxelization computed as described above. We used the minimal possible dilation ball kernel size to ensure a correct volume computation - visual checking of binary voxelization and value at least 3x surface area. We use *S*_1_ = *S*_topo_(*x, y, z*) and *S*_2_ = *S*(*x, y, z*) to compute the metrics of Fig. S3, S5.

### Reference surface inference for measurement of protrusion height

The height of a protrusion can only be defined relative to a reference shape. The optimal reference may differ from the choice of *S*_ref_(*x, y, z*). The cMCF *S*_ref_(*x, y, z*) generates the smoothest references that are easiest for equiareal spherical parameterization. However, it is a poor reference for measuring protrusion height as demonstrated poorer performance in the binary segmentation of protrusions, which frequently cuts into the cortical surface (Fig, 5A). u-Unwrap3D offers three ways to measure protrusion height in different representations.

#### Total curvilinear distance to guided curvature-minimizing flow reference shape (3D Cartesian)

The guided flow under a signed distance transform produces shapes guaranteed to be internal of *S*(*x, y, z*), with flow converging towards the medial-axis skeleton. Moreover, common morphological motifs like blebs, lamellipodia, filopodia and microvilli all have positive curvature tips or ridges. By constructing a guided flow prioritising erosion of positive curvature with minimal change of negative curvature, we generate a reference shape of minimum Gaussian curvature with precise removal of morphological motifs (Fig. S1B). The protrusion height at each vertex, *v* is the total distance traversed from iteration *i* = 0 to the stop iteration.

#### Total curvilinear distance to a reference shape based on GVF or non-rigid registration (3D Cartesian)

GVF is used to measure the protrusion height of *S*(*x, y, z*) to *S*_ref_(*x, y, z*) when *S*_ref_(*x, y, z*) is not derived from *S*(*x, y, z*) by an iterative flow, or is an independent shape, e.g., a different timepoint (Fig. 4D). *S*(*x, y, z*) and *S*_ref_(*x, y, z*) should be globally registered to remove translation and rotation. *S*(*x, y, z*) is then iteratively propagated along the GVF of *S*_ref_(*x, y, z*) until all vertices are on *S*_ref_(*x, y, z*). The protrusion height at each vertex, *v* is the total distance traversed from iteration *i* = 0 to the final iteration. This height is signed, positive for vertices external to *S*_ref_(*x, y, z*) and negative for vertices internal to *S*_ref_(*x, y, z*).

#### Height from a baseline topographic surface (3D Topography)

An optimal baseline surface for protrusion segmentation should be a (*u, v*) parameterized surface i.e. *S*_ref_(*d*_ref_ = *f*(*u, v*), *u, v*) where *f*(⋅) is injective so that every (*u, v*) pixel has associated only one surface point of *S*(*d, u, v*). This follows by contradiction. Suppose a surface, *S*(*d, u, v*) has points with the same (*u, v*) but different *d* coordinates. The points with higher *d* must therefore be part of a surface protrusion and thus *S*(*d, u, v*) cannot be a *S*_ref_(*d*_ref_, *u, v*). We find a suitable *S*_ref_(*d*_ref_, *u, v*) in topographic space by formulating the asymmetric least squares problem (ALS)^141^ to compute a smooth (*u, v*) parametrized 2D ‘baseline’ surface, *d*_ref_ = *f*_smooth_(*u, v*) where 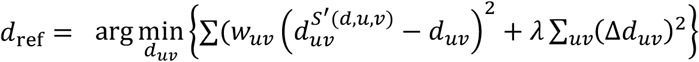 with asymmetric weights, *w* _*uv*_: *w* _*uv*_= *p* if 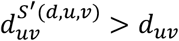 and *w*_*uv*_ = 1 − *p* otherwise. The regularization parameter, *λ* controls the contribution of the Laplacian Δ*d*_*uv*_ = ∇^2^*d*_*uv*_ and the smoothness of the resulting surface. The solution represents a surface intermediate between a (*u, v*)-parameterized approximation, 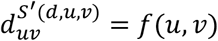 of the topographic surface *S*(*d* ≈ *f*(*u, v*), *u, v*) and the flat 2D-plane (*d* = 0) (Fig. 5B). The input (*u, v*) parameterized approximation 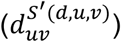 was computed by extending a vertical ray upwards at each (*u, v*) pixel and setting the *d* value as the highest value of the longest contiguous stretch of the topographic binary. The output is a (*u, v*) image of height *d* values. We downsample 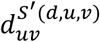 by a factor of 8 for computational efficiency and additional smoothness regularization and solve for *d*_ref_ by running 10 iterations of our modified 2D ALS^141^ using *p* = 0.25, *λ* = 1. The solution, *d*_ref_ is resized back to original (*u, v*) image size. The height, *h* of *S*(*d, u, v*) is the difference, *h* = *d* − *d*_*ref*_ between a vertex’s *d* coordinate and *d*_ref_ of the matching point on *S*_ref_(*d*_ref_ = *f*_smooth_(*u, v*), *u, v*) found by interpolation.

### Topography guided binary segmentation of protrusions

For cMCF binary segmentation of *S*(*d, u, v*), the baseline used is the 2d (*u, v*) plane, *S*_ref_(*d*_ref_, *u, v*) = *S*(*d* = 0, *u, v*) and the height is *h* = *d*. For improved segmentation, the baseline surface, *S*_ref_(*d*_ref_, *u, v*) was inferred using ALS as above with the height computed as *h* = *d* − *d*_ref_, relative to the matching point on *S*_ref_(*d*_ref_, *u, v*) with identical (*u, v*) coordinate. For both cMCF and ALS baseline, the mean height, 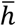 is used as the threshold for the initial binary segmentation, 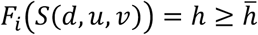 . This segmentation is postprocessed by applying mesh-based connected component analysis to remove small segmented regions with surface area < 200 voxels^2^; diffusing the result using two-class labelspreading^142^ with an affinity matrix, *A*, for 20 iterations, clamping ratio 0.99, and binarizing the label probability with a threshold of 0.25 at the start of each iteration. Lastly, any remaining small regions with surface area < 500 voxels^2^ was removed. The affinity matrix^107^, *A* is a weighted sum (*γ* = 0.9) of an affinity matrix based on geodesic distance, *A*_*dist*_ and one based on surface convexity, *A*_*convex*_; *A* = *γA*_*dist*_ + (1 − *γ*)*A*_*convex*_ of 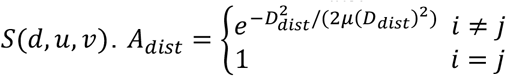 where *D*_*dist*_ is the pairwise Euclidean distance matrix between two vertices *i* and *j. A* 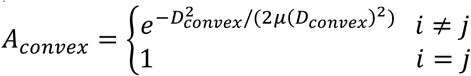 where *D*_*convex*_ is the pairwise Cosine distance, 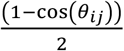 matrix of the dihedral angle, *θ*_ij_ between the normal vectors at two vertices *i* and *j. μ*(*D*) denotes the mean value of all entries in the matrix. This algorithm is also applicable to binary segment protrusions in *S*(*x, y, z*) using *S*_ref_(*x, y, z*) derived from guided curvature-minimizing flow or spectral decomposition. In the latter, protrusion height would need to be be obtained by the GVF approach.

### Topography guided instance segmentation of protrusions

Individual protrusions are segmented by identifying high curvature protrusive features and applying connected component analysis. The topographic mean curvature was computed 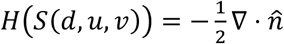 with the normal, 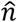 given by the unit gradient of the signed distance transform of the binary topographic volume of the cell, *I*_*binar*y_(*V*(*d, u, v*)) . We then derive a binary shell volume of the surface, *I*_*surf*_(*V*(*d, u, v*)), the intersection of the morphological dilation of *I*_*binar*y_(*V*(*d, u, v*)), with ball kernel size 2, and the morphological erosion of *I*_*binar*y_(*V*(*d, u, v*)), with ball kernel size 2. To identify high curvature surface regions, *H*_*high*_(*d, u, v*) for lamellipodia, we concatenate *H*(*S*(*d, u, v*)) Gaussian smoothed at σ = 1,3,5 as a 3-dimensional feature for all voxels in *I*_*surf*_(*V*(*d, u, v*)) and applied Gaussian mixture model (GMM) clustering (# classes = 3), using the class with the highest mean *H* to compute the binary curvature cutoff, *H*_*cutoff*_ . To identify *H*_*high*_(*V*(*d, u, v*)) for blebs and filopodia, which are circular and smaller, we use *H*(*S*(*d, u, v*)) Gaussian smoothed at the single scale, σ = 1 as a 1-dimensional feature for all voxels in *I*_*surf*_(*V*(*d, u, v*)) and apply kmeans clustering (# classes = 3), using the class with the highest mean *H*, to compute the binary curvature cutoff, *H*_*cutoff*_ . For computational efficiency, both GMM and kmeans clusterers are fitted on a random sampling of 10,000 voxels from the surface shell volume. Small regions with < 500 connected voxels are removed. Connected component analysis labels each disconnected region in *H*_*high*_(*V*(*d, u, v*)) as individual protrusions, *I*_*protrusions*_(*V*(*d, u, v*)). We expand labels by 3 voxels and transfer the segmentation to the surface mesh, *S*(*d, u, v*) by interpolation at the vertex coordinates, *F*_*protrusions*_(*S*(*d, u, v*)) for further surface-based postprocessing. We first apply the binary protrusion segmentation above, *B*_*protrusion*_(*S*(*d, u, v*)) to *F*_*protrusions*_(*S*(*d, u, v*)), taking the intersection and keeping segmentations with size > 100 voxel^2^ Cartesian 3D surface area. We diffuse segmentation labels with labelspreading, clamping ratio 0.99 for 10 iterations, with affinity matrix *A, γ* = 0.9 as defined above. We do not rebinarize the label probability at the start of each iteration. Finally, we apply *B*_*protrusion*_(*S*(*d, u, v*)) to the diffused segmentations to get the final instance segmentation labels, *F*_*protrusions*_(*S*(*d, u, v*)). Lastly, if necessary, segmentation labels at the borders can be corrected for spherical boundary conditions, by matching labels in Cartesian 3D. Like binary segmentation, the described approach can also be applied to segment *S*(*x, y, z*) directly. The advantage of the topographic representation is in segmenting small, less well-defined protrusions, which may be missed in *S*(*x, y, z*) thanks to *S*(*d, u, v*) integrating out the inherent curvature of *S*_ref_(*x, y, z*). The advantage of Cartesian 3D is not requiring special handling of the boundary conditions.

### Direct 2D unwrapping of protrusion submeshes

Segmented individual protrusions are open 3D surfaces with disk topology and can be directly unwrapped into 2D if they possess no holes or handles and have one boundary. This condition should be satisfied with our described approach. The genus, *g* of an open orientable surface with *b* boundaries is computed from the modified Euler characteristic, *χ* = 2 − 2*g* − *b* = #*V* − #*E* + #*F*. Similar to the spherical parameterization of closed 3D surfaces, the open 3D surface can be first mapped conformally to the unit disk and then relaxed to get an equiareal disk parameterization. The disk parameterization can then be mapped to a square parameterization with optional height ratio optimization to minimize the Beltrami Coefficient, similar to optimization of the (*u, v*) parameterization aspect ratio.

#### Quasi-conformal disk parametrization of genus-0 open surfaces

We obtain a quasi-conformal map of an open 3D surface to the unit disk by harmonic maps^143^. The open boundary vertices are mapped to the boundary of the unit circle, preserving the edge length fractions between neighbor points. The interior mesh vertices are then mapped into the disk interior by solving Laplace’s equation, ∇^2^*ϕ* = 0.

#### Equiareal disk parameterization by mesh relaxation

We relax the conformal disk parametrization whilst preserving the boundary topology using the area-preserving flow method^144^. We solve Poisson’s equation to compute a smooth vector field for diffusing the area distortion. Explicit Euler integration then iteratively advects vertex points with Delaunay triangle flips to ensure bijectivity at each iteration. The extent of area relaxation achieved is determined by the mesh quality and number of vertices with respect to the extremity of local area distortion. In general, relaxation was less stable compared to our relaxation for spherical surfaces above. Downsampling and uniform remeshing of the protrusion submesh by isotropic incremental remeshing is used to enable maximum area distortion relaxation. Alternatively, as for spherical surfaces, we can just minimize the proxy distortion factor of redistributing the disk face area after conformal mapping for stable equiareal parameterization.

#### Disk-to-square parameterization by analytical equations

To convert a unit disk parameterization to an *N* x *N* pixel image, we ‘square’ the disk using the elliptical grid mapping formula^145^, multiplying the resulting vertex coordinates by *N*/2 and interpolating the coordinates and associated vertex quantities onto a *N* x *N* pixel integer grid. This gives similar results but is significantly faster than solving the Beltrami equation^114^ to obtain a quasiconformal map.

#### Optimizing the height of the square to minimize metric distortion

As for (*u, v*) parameterization, (Fig. S4), elongated protrusions may be non-optimally represented by a square parameterization, leading to large conformal errors. We optimize for the aspect ratio in the same manner by finding a new height that minimizes the Beltrami Coefficient. The length ratio distortion is not a valid objective because the open surface is not homeomorphic to a sphere.

### Refinement of undersegmented blebs in 2D parameterizations

Given *S*(*x, y, z*) and the vertex ids corresponding to protrusion *i*, (*v*_*i*_) we impute any small holes in the segmentation to ensure the surface is continuous. We check that there is only one boundary loop. Otherwise, inner vertices not assigned to protrusion *i* are assigned so that the protrusion submesh, *S*_protrusion_(*x, y, z*) is a genus-0 open surface. We do this by applying mesh-based connected component analysis on the submesh formed by all vertex ids not part of protrusion *i*, {*v*}\{*v*}^*i*^. Any component with number of vertices <10% the total surface area of *S*(*x, y, z*) is assigned to protrusion *i* to form 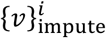. The submesh *S*_protrusion_(*x, y, z*) is formed from 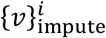. We downsample *S*_protrusion_(*x, y, z*) by ¼ the number of vertex points and remesh using ACVD or incremental isotropic remeshing to get a high quality istropic mesh, 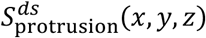 for computing an equiareal disk parameterization. 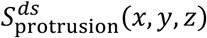 is directly unwrapped to 2D, here as a 128 x 128 pixel square image as described above. Positive curvature ‘seed’ regions are identified by thresholding the mean curvature of *S*(*x, y, z*) mapped to 2D, 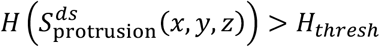 with a global *H*_*thresh*_ threshold, determined by Multi-Otsu thresholding of *H*(*S*(*x, y, z*)), then postprocessing by morphological closing, disk kernel radius 1 pixel. 3-class Otsu thresholding was applied to *H*(*S*(*x, y, z*)) to give two thresholds classifying surface regions as having negative, flat or positive mean curvature. All regions with mean curvature greater than the higher threshold *H*_*thresh*_ were positive curvature. Undersegmented blebs correspond to a binary composed of conjoined pseudo-circular regions. We use a gradient-based watershed^76,87,146^ process using the gradient of the Euclidean distance transform of the high curvature region binary to automatically separate conjoined blebs. Mesh matching and interpolation was used to map *H* and segmentation labels between *S*_protrusion_(*x, y, z*) and 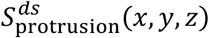. The refined segmentations were mapped as ‘seed’ labels from *S*_protrusion_(*x, y, z*) back to *S*(*x, y, z*) for every protrusion. The revised seed labels were then diffused across *S*(*x, y, z*) using the combined geometrical and convexity affinity matrix from above for 10 iterations with *α* = 0.99. The binary protrusion segmentation from above is applied, and any segmentation with Cartesian 3D surface area < 10 voxels^2^ removed to obtain the final refined protrusion segmentation instances.

### Surface curvature measurement

The mean curvature, *H* of a 3D surface was measured as the divergence of 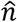, the unit surface normal^124^,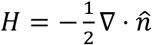. The surface mesh is voxelized to a binary volume, *B* and 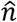 is computed as the gradient of the signed distance transform of *B* with the Euclidean distance metric. *H* computed in this manner compared to direct computation using discrete differential geometry^124^ or quadric plane fitting^147^ is less affected by the number of mesh vertices or the mesh quality. However, the computation is slower, and applicable only when the surface mesh can be voxelized to a proper closed volume. Therefore, we use *H* computed from principal curvatures based on quadric plane fitting^147^ when *H* is not used for statistical analysis, including finding the optimal unwrapping axis for uv-unwrapping.

### Mesh quality measurement

The radius ratio 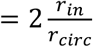, defined as twice the ratio between inradius and circumradius is used to measure the face quality for a triangle mesh during shrink-wrapping. It is a mesh quality measure in the sense that the radius ratio obtains its maximum value of 1 for an equilateral triangle; the shape which jointly maximizes all internal angles and gives the best conditioning number for the mesh Laplacian matrix^148^. If equiareal relaxation terminates early or does not achieve the minimum, we also recommend checking that the minimum and maximum internal angles of the triangle faces lie between ≈30^0^-120^0^ to avoid singular matrices.

### Surface rendering

Triangle meshes were exported from Python using the Python Trimesh library into .obj mesh files and visualized in MeshLab^149^. Volumetric images were rendered in Fiji ImageJ through the volume viewer plugin, and intensities were contrast-enhanced for inclusion in the figures using Microsoft PowerPoint. The local surface maximum intensity projection image of Fig. 7C was produced by extending z-axis (depth) rays at every xy pixel, and taking the maximum intensity of voxels within ±9 voxels (±1µm) of the cell surface.

## Datasets

We describe the methods and parameters used to generate the main paper examples to highlight the numerous different ways u-Unwrap3D can be implemented. The choices do not reflect necessarily the most optimal parameters. They are reported here only with the intention of demonstrating the flexibility and tolerance of u-Unwrap3D to diverse experimental data.

### Cells with morphological motifs

#### Blebs, lamellipodia and filopodia (Fig. 1,2)

Cell surfaces were segmented and surface protrusions classified using u-Shape3D and acquired from high resolution light sheet microscopy^4,150^ as previously described^39,81^. u-Unwrap3D was applied with the following parameters for each step without shrink-wrapping: Step 1, cMCF with maximum iterations = 50, *δ*_*t*_ = 5 × 10^−4^, a rate-based stopping threshold, Δ_*thresh*_ = 1 × 10^−5^ for blebs, = 1 × 10^−5^ for lamellipodia, = 1 × 10^−4^ for filopodia, *S*_ref_(*x, y, z*) mesh voxelization with morphological dilation and erosion with ball kernel radius 5 voxels, Gaussian smoothing σ = 1 of binary volume and initial *S*_ref_(*x, y, z*) meshing with marching cubes at isovalue 0.5, ACVD remeshing with number of clusters = 90% the number of vertices in the marching cubes mesh; Step 2, direct quasiconformal parameterization; for Step 3, equiareal relaxation with maximum iterations = 100, *δ*_*t*_ = 0.1, stepsize *ε* = 1 ; for Step 4, *w* = |*H*(*S*_ref_(*x, y, z*))|, *w*′ = *H*(*S*_ref_(*x, y, z*)) for determining the unwrapping axis, a 512 x 1024 pixel (*u, v*) grid; for Step 5, Euclidean distance transform for external and internal propagation, *α* = 0.5 voxel steps, *D*_*in*_ = 40 steps, 2D robust smoothing^151^ with smoothing factor = 50 for each iteration; for Step 6, rebinarization of the topographic 3D mapped cell binary at a threshold of 0.5, Gaussian smoothing σ = 1 and initial marching cubes meshing at isovalue 0.5, and ACVD remeshing with number of clusters = 50% the number of vertices in the marching cubes mesh. Without shrink-wrapping the choice of *S*_ref_(*x, y, z*) is limited to smooth shapes with small holes that can be closed by voxelization and morphological processing (Fig. S1B). As shown by the filopodia in Fig. 2, 5, Fig. S5 this produces a *S*_ref_(*x, y, z*) that under-represents the filopodia tips in topographic space (Fig. S5B), and causes the larger branches to be segmented as protrusions (Fig. 5). With shrink-wrapping, any cMCF *S*_ref_(*x, y, z*) can be selected, allowing an optimal selection of *S*_ref_(*x, y, z*) that accurately represents the filopodia in topographic space (Fig. S5C). This revised topographic mesh was also obtained using Poisson distance transform for internal propagation, which is stable and faster than robust smoothing.

#### Microvilli (Fig. 2)

Blasted human CD8^+^ T cells, were produced by activating naïve T cells isolated from PBMCs using anti-CD3/CD28 Dynabeads for 2 days, then rested for 5 days after removing the beads. Cells were frozen on day 5 of resting and thawed 48 hours before use. T cells were grown in complete RPMI-1640 (10% FCS, 1% Pen/Strep, 1% Glutamine, 1% HEPES) + 50 U/ml of IL-2. To image membrane protrusions, 8-well glass-bottom IBIDI chambers were coated with 1% BSA for 1h at room temperature to prevent extensive surface adhesion of T cells to the surface. 0.5x10^6^ blasted human CD8+ T cells were labelled with CellMask DeepRed diluted to a 1x working solution in imaging buffer (colorless RPMI-1640 with 1% added Pen/Strep, 1% glutamine, 1% HEPES) for 30 min at 37°C. Cells and the glass slides were washed and resuspended in pre-warmed (37°C) imaging buffer. 0.1-0.2x10^6^ cells were gently added to the coated glass slides and left to settle for 15 min before imaging. Movies of the cells were imaged using the Zeiss Lattice Lightsheet microscope 7 (LLSM7) using the 641nm laser at 4% power with 4 ms of exposure. ∼150 sections were used per volume (for single cells), with a complete cell volume taken every second. Individual timepoints were registered by rigid body transformation. Cells were then segmented by binary Otsu thresholding after preprocessing the registered timepoints by adaptive histogram equalization and gamma correction. To segment individual protrusions, the surfaces of individual timepoints, *S*(*x, y, z, t*) were extracted by Marching cubes (Gaussian presmoothing, σ = 1.1 at contour level 0.5, ACVD remeshing, 0.5 downsample). Subsequently, unsupervised mean curvature and connected component analysis as described above was applied directly in Cartesian 3D with *S*(*x, y, z, t*). The cortical shape does not change significantly across time. Therefore, u-Unwrap3D was applied to the mean temporal shape, ⟨*S*(*x, y, z*)⟩_*t*_, derived by Otsu thresholding of the mean of individual binary segmentation volumes, and surface meshing with Marching cubes (Gaussian presmoothing, σ = 3. at contour level 0.5, ACVD remeshing, 0.2 downsample). We use this as *S*_ref_(*x, y, z*) = ⟨*S*(*x, y, z*)⟩_*t*_. It is also already genus-0, as the cortical shape is almost spherical. We projected *S*_ref_(*x, y, z*) to the sphere using the direct quasiconformal map^68^ and performed equiareal distortion relaxation with maximum iteration=50, *δ*_*t*_ = 0.1, *ε* = 0.1 . The microvilli are relatively uniformly distributed therefore we mapped to a 256 x 512 uv image without optimizing unwrapping axis or aspect ratio. As *S*_ref_(*x, y, z*) is convex, we built the topographic space using Forward Euler for both external (to cover the maximum length microvilli in the video) and internal (30 voxels) propagation with *α* automatically set as the mean Cartesian length of a *u* and *v* pixel. Individual timepoint topographic meshes were computed by Marching cubes meshing (Gaussian presmoothing, σ = 1, contour level=0.5, ACVD remeshing, 0.5 downsample) the binary volumes mapped into the constructed topographic space.

#### Branched (Fig. 2)

This phospo-defective A549 lung cancer cell with hierarchical branching was computer-generated using Filogen^77^ and part of the example dataset downloaded from the Filogen software website (phospho-defective/ID450), https://cbia.fi.muni.cz/research/simulations/filogen.html. The intensity image and segmentation masks with unique labels for individual branches are provided as a volume image with voxel size 0.126 x 0.126 x 1.0 microns. After rescaling to isotropic voxels, we used marching cubes at contour level 0.5, after presmoothing the binarized mask with a Gaussian, σ = 0.5, and ACVD remeshing to extract the mesh. We find *S*_ref_(*x, y, z, t*) of minimum Gaussian curvature, using active contour (*γ* = 1, *α* = 0.1, *β* = 0.5) and curvature-weighted gradient of the Euclidean distance transform (Fig. S1). We use the genus-0 *S*_ref_(*x, y, z, t*), after voxelizing *S*_ref_(*x, y, z, t*), dilating by *r*_*dilate*_ = 5 voxels, eroding by *r*_*erosion*_ = 1 voxels, and remeshing with marching cubes, isotropic meshing with ACVD and downsampling to 0.25x vertices. We conformal mapped *S*_ref_(*x, y, z, t*) to the sphere using the iterative method, and performed equiareal relaxation using maximum iterations = 350, *ε* = 1, *δ*_*t*_ = 0.1. The unwrapping axis was optimized based on the last timepoint with maximal branching, using a binarized positive mean curvature, *w* = *H*(*S*(*x, y, z, t* = *T*)) > 0, to treat all branching equally, *w*^′^ = *H*(*S*(*x, y, z, t* = *T*)), and (*u, v*) parameterized to a 512 x 1024 image, then width optimized by minimizing the Beltrami coefficient. We construct the topographic space, *α* = 1, propagating outwards 150 voxels with Forward Euler and uniform smoothing (window size 1/5 the uv image width), propagating inwards 20 voxels with active contour (*γ* = 1, *α* = 0.1, *β* = 0.5) and Euclidean distance transform. Topographic meshes were obtained with marching cubes, contour level 0.5, presmoothing the topographic binary with Gaussian σ = 0.5, and ACVD remeshing.

#### Mixed (Fig. 2)

The cell aggregate comprising transformed HBEC cells expressing eGFP-Kras^V12^ imaged by meSPIM was published previously^*78*^. *S*(*x, y, z*) was obtained by Marching cubes meshing (presmoothing, σ = 1, level=0.5, ACVD remeshing, 0.9 downsample) the binary segmentation of the aggregate, obtained by combining one binary segmentation targeting the low-frequency core shape and another, targeting the high-frequency surface protrusions. The image was preprocessed by gamma correction (*γ* = 0.8), Wiener deconvolved with the meSPIM PSF and percentile intensity normalized. We obtain the core shape segmentation by anisotropic smoothing with guided filtering^152^ (radius=5, regularization=0.1), mean intensity binary thresholding, largest component with binary closing and hole-filling. We obtain the protrusion segmentation by a two-level approach. The first obtains a base segmentation using the lower threshold from applying 3-class Otsu thresholding to the preprocessed image, largest component, then binary closing and hole-filling. The second segments a normalized difference of Gaussian (*DoG*) image, *DoG* computed by subtracting the Gaussian filter, σ = 5 of the preprocessed image from the preprocessed image which enhances ridge-like features, 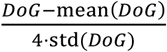 that enhances ridge-like features, where mean(⋅) and st*d*(⋅) is the mean and standard deviation of the 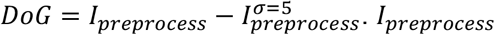 denotes the preprocessed image and 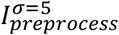 denotes the preprocessed image after Gaussian filtering, σ = 5. The final protrusion binary is the union of the two. The core segmentation was eroded to match as-best-as-possible the protrusion segmentation. The final binary segmentation of the aggregate is the largest component after the union of core and protrusion binaries, after hole-closing and hole-filling. u-Unwrap3D was applied with the following: cMCF *S*_ref_(*x, y, z*) with maximum iteration=50, *δ*_*t*_ = 5 × 10^−4^; first change-point stopping iteration based on mean absolute Gaussian curvature with window size = 9, minimum peak height=1 × 10^−7^, minimum peak distance=5; genus-0 *S*_ref_(*x, y, z*) after voxelization, *r*_*dilate*_ = *r*_*erode*_ = 5, and Marching cubes meshing (presmoothing, σ = 1, level=0.5, ACVD remeshing, 0.1 downsample); sphere parameterization with direct quasiconformal map, equiareal relaxation, maximum iterations=100, *δ*_*t*_ = 5 × 10^−3^, *ε* = 0.5 . (*u, v*) parameterization, initial 1024 x 2048 grid with unwrapping axis optimization, *w* = |*H*(*S*_ref_(*x, y, z*))|, *w*^′^ = *H*(*S*(*x, y, z*)), and Beltrami coefficient optimized final grid size; topographic space construction with *α*-step size set automatically as the mean of the Cartesian length represented by *u* and *v* pixels, internal (60 voxels) and external (auto-set to cover maximum protrusion) propagation with Forward Euler and Euclidean distance transform; and topographic mesh by Marching cubes (presmoothing, σ = 1, level=0.5, ACVD remeshing, 0.5 downsample). The binary-equivalent of the u-Segment3D instance cell segmentation^76^ is not identical to the u-Unwrap3D analyzed binary in Fig. 2. To map the u-Segment3D instance cell masks, we expanded the labels without overlap by 35 voxels and sampled it at *S*(*x, y, z*) vertices. This was mapped into topographic space to sample onto *S*(*d, u, v*).

### Shrink-wrapping datasets (Fig. S2)

u-Unwrap3D genus-0 shrink-wrapping is performed on an image grid whose size is automatically inferred from the input mesh vertices. The process can be slow for large shapes, but can be expedited by dividing the mesh vertices by a constant factor, then remultiplying this factor at the end. This will effectively run shrink-wrapping on a downsized grid. The default shrink-wrapping parameters that worked for the majority of cells are: GVF: 15 iterations of the GVF^67^ diffusion with regularization, *μ* = 0.01, applied to the attractor field of *S*(*x, y, z*) composed using the Euclidean distance transform of the voxelized mesh; mesh voxelization: *r*_*dilate*_ = 3, *r*_*erode*_ = 2 ; alpha wrapping search parameters: alpha=0.15 of the bounding box extent, offset=0.001; each shrink-wrap cycle: first cycle, 20 epochs, subsequent refinement cycles with punchout, 10 epochs; initial shrink-wrap epoch: at start, decimating mesh to half the number of vertices to remove small edges and upsampling by face subdivision to exceed minimum number of vertices then propagating along GVF gradient by 10 iterations, initially with step size=1, minimum mesh number vertices=80,000, minimum step size=0.24, step decay fraction = 0.5, active contours (*γ* = 1, *α* = 0.1, *β* = 0.5); and subsequent epochs for refinement cycles: as for initial cycle but reduced *r*_*dilate*_ = 2 and step decay fraction = 1.0 (i.e. no decay). We perform punchout refinement between cycles by first detecting all unconverged triangles part of ‘plate-like’ structures, defined by having flat mean curvature, *H* < 0.01 and radius 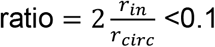 or nearest distance of vertex to *S*(*x, y, z*) > 1.25 + 2 × standard deviation of nearest distance of all vertices or part of face with area > mean + 2 × standard deviation of all face areas. Then we perform mesh-based connected component analysis, removing regions with fewer than 10 vertices. Lastly, we voxelize the nonconverged vertices, and dilate with a ball kernel of radius 3 to ensure removal involves a whole continuous patch. The final nonconverged vertices marked for removal are obtained by sampling the voxelized segmentation at vertex coordinates. To patch and ensure genus-0, we exhaustively detect each open boundary loop and fan-stitch the boundary vertices to form triangle faces with a new point - the mean of all boundary vertices. Below, we note any individual modifications for the different exemplars. Movie S2 animates the shrink-wrapping process for all shapes.

#### Lamellipodia (Fig. S1C)

Genus-0 shrink-wrap by alpha wrapping with parameters obtained by interval bisection search. As this shape is smooth, obtained from cMCF, shrink-wrapping with GVF does not further improve the genus-0 alpha wrapped mesh (Movie S2). This holds generally for reference shapes devoid of surface protrusions.

#### Filopodia (Fig. S1D)

The initial genus-0 alpha wrap found by interval bisection does not provide a tight wrap. Shrink-wrapping was performed with standard parameters. After the first cycle, two punchout refinement cycles was then performed.

#### Branched organoid (Fig. S1E)

The organoid was cultured and imaged as described in detail below for timelapse u-Unwrap3D analysis. This example was acquired on a Nikon AXR laser scanning confocal microscope with a z-step of 0.426 μm and lateral xy pixel size of 0.426 μm. It was segmented by binary Otsu thresholding. We modify the first shrink-wrap cycle to have a tighter voxelization, *r*_*dilate*_ = 2, *r*_*erode*_ = 2. After the first cycle, four punchout refinement cycles were performed.

#### Ruffles (Fig. S1F)

This COR-L23 cell (Human Caucasian lung large cell carcinoma) and its microscope acquisition were previously published^76^. The cell was segmented with binary Otsu thresholding following Wiener deconvolution using the meSPIM PSF as a synthetic point-spread function. Inhomogeneous staining results in an incomplete surface mesh with large holes. Standard genus-0 alpha wrap found by interval bisection finds the tightest wrap around the shape. To impute the large holes, we manually specify the initial alpha wrap with alpha=0.5 of the bounding box extent. This wraps the entirety of the shape whereas the smallest alpha found by interval bisection preferred to wrap the internal without closing outer holes. We then performed GVF refinement with standard parameters. After the first cycle, two punchout refinement cycles were performed. GVF gradients are not defined where there are holes. Yet, combined with active contours, the shrink-wrap imputed a minimum bending thin plate for each hole. This example illustrates how our shrink-wrapping accommodates partially segmented cells.

### 3D timelapse lightsheet imaging of cell with polarized blebbing with simultaneous cortical shape changes

The analysis of this cell in Fig. 3 demonstrates the standard prototypical workflow for applying u-Unwrap3D to temporal data.

#### Cell culture and timelapse imaging

MV3 melanoma cells expressing GFP-AktPH were cultured and imaged using lightsheet microscopy tunnelling through collagen by blebbing as published previously^40^.

#### Cell segmentation and surface meshing

Cells were segmented at each timepoint using the two-level approach for segmenting the cell aggregate described above. The surface, *S*(*x, y, z, t*) was extracted with Marching cubes (presmoothing, σ = 1, level=0.5, ACVD remeshing, 0.9 downsample).

#### u-Unwrap3D analysis

cMCF was applied to individual *S*(*x, y, z, t*), maximum iterations = 50, *δ*_*t*_ = 5 × 10^−4^. To ensure cMCF *S*_ref_(*x, y, z, t*) is temporally consistent, we computed the mean absolute Gaussian curvature curve across all timepoints. We then used this mean curve to obtain the same stopping iteration number by rate of change, Δ_*thresh*_ = 5 × 10^−5^ for every timepoint. Genus-0 *S*_ref_(*x, y, z, t*) was obtained by Marching cubes meshing (presmooth, σ = 1, level=0.5, ACVD remeshing, 0.1 downsample, minimum number of vertices=60,000) of the voxelized *S*_ref_(*x, y, z, t*) (*r*_*dilate*_ = *r*_*erode*_ = 5). Equiareal sphere parameterization was obtained by direct quasiconformal mapping, with distortion relaxation (max iterations=100, *δ*_*t*_ = 0.1, *ε* = 1). To optimize the unwrapping axis and an initial 256 x 512 (*u, v*) map aspect ratio by Beltrami coefficient, we used the surface *S*(*x, y, z, t*) with most blebs, found by the maximum sum of mean curvature, as reference, using its equiareal sphere with *w* = |*H*(*S*(*x, y, z*))|, *w*′ = *H*(*S*(*x, y, z*)). We constructed topographic spaces relative to the (*u, v*) parameterized reference at each timepoint, *t, S*_ref_(*u, v*)(*t*) using *α* = 0.5 voxel, and Forward Euler propagating externally 100 voxels and internally 50 voxels, uniform smoothing with window size 1/5 the (*u, v*) map width. The constructed volumes were all the same size in voxels with *d* = 0 corresponding to their respective *S*_ref_(*x, y, z, t*). Individual timepoint topographic meshes were obtained by Marching cubes meshing (presmoothing, σ = 1, level=0.5, ACVD remeshing, 0.5 downsample) of the binary segmentation of *S*(*x, y, z, t*) remapped into their topographic space.

### 3D timelapse lightsheet imaging of interaction between an NK and cancer cell

#### Cell culture and timelapse imaging

NK-92 natural killer (NK) cells expressing Life-Act-mScarlet and K562 leukemic cells expressing Lck-mVenus imaged using an oblique plane microscope was published previously^86^.

#### Cell segmentation and surface meshing

The NK and leukemic cells were captured in different color channels and segmented independently. To produce candidate cell instance segmentations for each timepoint, we applied u-Segment3D^74^ with pretrained 2D Cellpose models, which acted as shape priors and could impute complete cell shapes to imperfect membrane staining, with diffusion and guided filter refinement. Given the NK cell moves at the boundary of the field of views, the segmentations may capture the whole cell, parts of the cell or neighboring NK cells, or parts of segmentations of the cancer cell, whose fluorescence bleeds into the NK cell channel. Crucially, they still represent volumes, not hollow surface shells as would have been obtained by Otsu-based thresholding. We generate the final segmentations with the help of temporal tracking. We merge segmentations at timepoint 0 to get the whole NK cell. This served as the template to identify merges in subsequent timepoints. Specifically, the NK segmentation at time *t* − 1 is used to find candidates at time, *t* sharing significant volume overlap (> 5000 voxels). These candidates are then merged to form the NK cell at time *t*. We check if this NK cell overlapped with other segmentations at time *t*. If there is overlap > 100 voxels, we dilated the other segmentation with ball kernel radius 1 and removed the overlapped volume from the NK cell segmentation. The leukemic cell was segmented similarly.

#### Determining time of stable synapse formation, t_sy_napse

We identify stable synapse formation as maintaining a constant overlapped volume between NK and leukemic cell. We computed the overlap by taking the binary segmentation of NK and leukemic cells dilated with a ball kernel of radius 1 voxel at each timepoint, and summing their intersection. The overlapped volume increases over time before it plateaus. We determined a cutoff by applying Otsu thresholding to the overlapped volume timeseries. The first timepoint whose overlap volume exceeded this cutoff was used to define the onset of the plateau phase, corresponding to the first establishment of a stable synapse.

#### u-Unwrap3D analysis

We apply u-Unwrap3D to the NK cell to analyze the interaction from its perspective. We find *S*_ref_(*x, y, z, t*) by cMCF (iterations=50, *δ*_*t*_ = 5 × 10^−4^). To ensure temporal consistency, the stopping iteration was determined from the temporal mean of absolute Gaussian curvature curves, using the first stop found by the changepoint method (windows size=9, minimum peak height= 1 × 10^−7^, minimum peak distance=5). We register *S*_ref_(*x, y, z, t*) to a fixed temporal reference shape ⟨*S*_ref_⟩(*x, y, z*), which we define to be the temporal mean of all *S*_ref_(*x, y, z, t*) (see below). Equiareal sphere parameterization of ⟨*S*_ref_⟩(*x, y, z*) by direction quasiconformal map, and relaxation (maximum iteration=250, *δ*_*t*_ = 0.5,*ε* = 0.5). The unwrapping axis of ⟨*S*_ref_⟩(*x, y, z*) was optimized by remapping the binary of the overlapped surface between the NK and leukemic cell at *t*_*s*y*napse*_ to the ⟨*S*_ref_⟩(*x, y, z*) geometry. This ensured that the synapse will be mapped to the center of the (*u, v*) parameterization. The initial 512 x 1024 (*u, v*) parameterization was optimized for aspect ratio by Beltrami coefficient using range *h* = [0.05,5] inclusive. The (*u, v*) parameterization of *S*_ref_(*x, y, z, t*) was then found by applying the registration transforms to ⟨*S*_ref_⟩(*u, v*) . The topographic space of each *S*_ref_(*x, y, z, t*) was then built by Forward Euler propagation of its *S*_ref_(*u, v*)(*t*) with Euclidean distance transform, externally for 100 voxels and internally for 25 voxels with uniform smoothing, window size = 1/8 width of (*u, v*) map. *α* was set to the mean Cartesian length represented by a *u* and *v* pixel.

#### Registering S_ref_(x, y, z, t) to temporal reference shape

We use volumetric registration-based methods. All *S*_ref_(*x, y, z, t*) were voxelized, and the mean across all timepoints rebinarized with cutoff 0.5 was defined as the fixed temporal reference, ⟨*S*_ref_⟩(*x, y, z*). *S*_ref_(*x, y, z, t*) binary volumes were first individually registered to ⟨*S*_ref_⟩(*x, y, z*) using a similarity transform to account for scale, rotation and translation. Then, non-rigid registration was performed using symmetric demons in a multiscale manner, at downsampling of 8x, 4x, 2x, then 1x, maximum 500 iterations at each scale. Finally, we use the established geometric transformations to register the surface representations, sampling the non-rigid displacements at the respective mesh vertices.

#### Information flow analysis

We use u-Infotrace^88^, a framework we developed previously to apply 1D causal measures, to compute multiscale pixel-based information flow in 2D videos. We apply u-Infotrace to the (*u, v*) maps of *S*(*x, y, z, t*) mean curvature after weighting for area distortion and normalization, using dynamic differential covariance as the 1D causal measure, ridge regression regularization=1x10^-2^, and a multiscale of 1x, 2x, 4x, 8x, 16x downsampled images. Signals in (*u, v*) were corrected by weighting it with the (*u, v*) pixel area of *S*_ref_(*u, v*)(*t*) at time, *t*. This follows as each signal is a scalar, *F* and the mean of *F* integrated over a local surface area, Ω_*S*_ of the total surface, *S* in Cartesian 3D is equivalent to computing a weighted mean over the corresponding (*u, v*) area,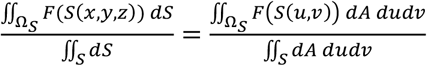. The weight, *dA* is the magnitude of the differential area element, 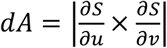 described above. The weight is computed and applied at each timepoint, *t* as *S*_ref_(*u, v*)(*t*) varies. The corrected signals are min-max normalized using the minimum and maximum value across time.

### 3D timelapse confocal imaging of K14+ cells in breast cancer organoids

*Mice*. All mouse husbandry, procedures performed, and assay end points were performed in accordance with protocols approved by the Johns Hopkins Medical Institute Animal Care and Use Committee (ACUC; MO23M200) as accredited by AAALAC international. The FVB/NJ (RRID: IMSR_JAX:001800) and FVB/N-Tg(MMTV-PyVT)634Mul/J (RRID:IMSR_JAX:002374)^90^ lines were obtained from The Jackson Laboratory and were kept on an FVB/N background in a specific pathogen-free facility. For timelapse experiments, MMTV-PyMT mice were crossed with mT/mG mice and K14-GFP-actin mice.

#### Isolation and culture of murine mammary tumor organoids

Organoids were isolated fresh for every experiment, as previously described^91,153^.^91,153^. Tumors were collected from MMTV-PyMT mice with a tumor burden of 16-20 mm. Tumors were minced with a scalpel and enzymatically digested in solution on a shaker at 180 RPM at 37°C for 60 min^153,154^.^153,154^. Digestion solutions were as follows: Collagenase (2 mg mL ^−1^) (Sigma-Aldrich, C2139), trypsin (2 mg mL^−1^) (Gibco, 27250-018), 5% fetal bovine serum (FBS) (Sigma-Aldrich, F0926), insulin (5 μg mL^−1^) (Sigma-Aldrich, I9278), and gentamicin (50 μg ml^−1^) (Gibco, 15750) in 30 mL of Dulbecco’s modified Eagle’s medium/nutrient mixture F12 (DMEM-F12, Gibco, 10565-018). After digestion, tumor tissue was centrifuged at 400g for 10 min. After supernatant removal, deoxyribonuclease (DNase) (2 U μL^−1^) (Sigma-Aldrich, D4263) was added in DMEM-F12 and incubated for 5 min at room temperature. Next, the tissue was centrifuged at 400g for 10 min, followed by three differential centrifugations (400g, 3s) to separate the epithelial cell clusters from stromal cells. Small and large organoids (100-200 cells versus 250-500 cells, respectively) were then separated using differential centrifugation at a lower speed (150g, 3s) for tumor organoids. Large tumor organoids were then embedded in a neutralized 3D fibrillar rat tail collagen I matrix (3 mg/mL, 354236; Corning). Organoids were seeded at a density of one organoid per μL. Collagen gels were allowed to polymerize for 0.5–1 h at 37°C, after which organoid growth media was added to each well. Murine organoids were cultured in DMEM-F12, 1% (v/v) insulin-transferrin-selenium-ethanolamine (Life Technologies, 51500-056), 1% (v/v) penicillin-streptomycin (Sigma-Aldrich, P4333), and 2.4 nM fibroblast growth factor 2 (FGF2) (Sigma-Aldrich, F0291).

#### Timelapse imaging

Confocal images were acquired using a Nikon AXR laser scanning microscope with a tunable DUX-VB gas detector and a 20X Plan Apo λD objective with a numerical aperture (NA) of 0.8 and an 800 µm working distance. Images were acquired using the NIS Elements software. Timelapse images were taken as z-stacks with a z-step of 2 μm, 1024x1024 pixel size with lateral xy resolution of 0.85 μm. Organoids were seeded on day 0 and, starting on day 1, were imaged every 20 min for 5 days.

#### Cell segmentation and surface meshing

The video was downsampled in xy so that the volume has isotropic voxels of 2 μm. Individual timepoints were then globally registered using the middle timepoint as reference by similarity transform to remove scaling, rotation and translation. Organoids were then segmented using the MTMG channel. The binary segmentation was obtained by binary Otsu thresholding of the gamma corrected image (*γ* = 0.5) followed by binary closing, retaining the largest connected component, and binary infilling. Segmentations were temporally smoothed using a moving average of 5 frames, rebinarizing at a cutoff 0.1, keeping the largest connected component, and binary eroding with a ball kernel of radius 1. Organoid surfaces were then obtained by Marching cubes (presmoothing, σ = 0.5, level=0.5, ACVD remeshing, 0.9 downsample).

#### u-Unwrap3D analysis

In this case, the organoid surface is also the reference surface, *S*(*x, y, z, t*) = *S*_ref_(*x, y, z, t*). We first shrink-wrapped to obtain a genus-0 *S*(*x, y, z, t*) and smoothed the mesh with one iteration of cMCF (*δ*_*t*_ = 2.5 × 10^−5^). The mesh was first decimated to 1/3 the number of vertices, and subsequently decimated by a factor 3/4 until its mesh quality > 0.1. For each decimation, the mesh is upsampled by subdivision if the number of mesh vertices is less than 10,000. This procedure still retains shape accuracy but reduces the density of vertices. It is necessary to achieve optimal equiareal distortion relaxation (iterations = 500, *δ*_*t*_ = 0.1, *ε* = 1) of the iterative spherical conformal map of *S*(*x, y, z, t*) . Alternatively, we could have generated a dilated *S*(*x, y, z, t*) by voxelization with *r*_*dilate*_ > *r*_*erode*_ + 2 to avoid sharp branch points and narrow branches. We then used the mid timepoint as the reference to determine the unwrapping axis, *w* = *H*(*S*(*x, y, z*)) > *H*_*cutoff*_ where *H*_*cutoff*_ = Otsu threshold of *H*(*S*(*x, y, z*)) and *w*^′^ = *H*(*S*(*x, y, z*)). We used a 512 x 1024 (*u, v*) grid for parameterization without aspect ratio optimization, and constructed the topographic space for each timepoint, *t* by propagating 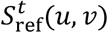 externally by 40 voxels, and internally by 40 voxels, with step size, *α* = 0.5 voxels using active contours and the Poisson distance transform.

#### Measuring proximal surface signal intensity

We resampled the MTMG and K14 signal into topographic space where each *d -* column of (*u, v*) pixel intensities corresponds to sampling the intensity along the steepest gradient external and internal from the surface *S*(*x, y, z, t*) (i.e. *d* = 0). We then construct the proximal surface signal intensity (*u, v*) map for analysis as the maximum intensity projection of the topographic subvolume given by *d* = [−5,0] inclusive, corresponding to a depth of 5 µm.

#### Measuring branch length

We measured the length of each *S*(*x, y, z, t*) vertex from *S*(*x, y, z, t* = 0) to compute a scalar (*u, v*) map of branch length from the starting organoid shape. To measure the length we constructed the gradient vector flow with *S*(*x, y, z, t* = 0) as the attractor using the Poisson distance transform. Each *S*(*x, y, z, t*) was then propagated by active contours (iterations=200, *γ* = 1, *α* = 0.1, *β* = 1.0, robust Laplacian) along this gradient vector flow to converge onto *S*(*x, y, z, t* = 0). The convergence iteration of *S*(*x, y, z, t*) was determined as the first iteration where the chamfer distance between the propagated *S*(*x, y, z, t*) and *S*(*x, y, z, t* = 0) was <1.5 voxels. The branch length is the total distance propagated under active contours of each vertex of *S*(*x, y, z, t*) from iteration 0 to the convergence iteration. The length is positive if the vertex of *S*(*x, y, z, t*) is located external of *S*(*x, y, z, t* = 0) and negative if located internal of *S*(*x, y, z, t* = 0).

#### Information flow analysis

The multiscale information flow of each signal and branch length were computed independently with u-Infotrace^88^ as for the NK and cancer cell analysis above, with the same metric distortion correction and normalization of signals, and the same parameters but reduced ridge regression regularization = 1x10^-3^. The magnitude of flow indicates the causal strength. If A causes B and vice versa in a region, we expect the flow to be identical, with similar magnitude and direction. We define a correlation measure of flow, 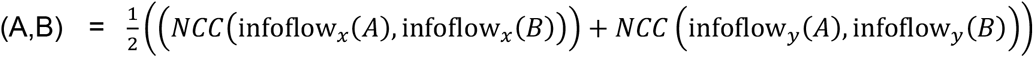 matching of *x* and y components of the information flow using normalized cross-correlation (NCC)^155^ (implemented by scikit-image, skimage.feature.match_template function). We used a paired t-test to assess statistical significance of the information flow correlation between different pairs of variables.

### 3D timelapse lightsheet imaging and analysis of ruffles

*Cell culture and timelapse lightsheet imaging*. SU.86.86 cells were purchased from American Type Culture Collection (CRL-1837). The cells were transfected with integrating lentiviral plasmids carrying genes for myristoylated CyOFP1 and Tractin-mEmerald. The cells were cultured in RPMI medium supplemented with 10% fetal bovine serum and 1% anti-anti (Gibco 15240062), at 37°C in a humidified incubator and 5% CO_2_. SU.86.86 cells were imaged on fibronectin-coated coverslips on a custom axially swept light sheet microscope^156^.^156^. The microscope detection system comprises a 25X NA1.1 water immersion objective (Nikon, CFI75 Apo, MRD77220) and a 500mm tube lens. The illumination was done through a 28.6X NA=0.66 water immersion objective (Special Optics, 54-10-7). The movie analyzed in Fig. 5 is a total of 30 frames acquired at a frequency of 2.27 s per frame. Each frame is a two-channel 151 x 1024 x 1024 size 3D volume with a voxel resolution of 0.300 x 0.104 x 0.104 µm.

#### Cell segmentation and surface meshing

All timepoints were rigid registered volumetrically to the first timepoint, *t* = 0 to compensate for drift. The CyOFP1 image was also rigid registered to the Tractin-mEmerald in each timepoint. For every timepoint, the volumetric image was segmented using a multi-level method^157^ involving local contrast enhancement, deconvolution and edge enhancement and surface meshed as described above to obtain the surface mesh, *S*(*x, y, z*) . The vertex Tractin-mEmerald and CyOFP1 intensities were calculated by extending a trajectory to an absolute depth of 1 µm along the steepest gradient of the distance transform to the mesh surface, and taking the mean intensity along the trajectory.

#### u-Unwrap3D analysis

The first timepoint surface mesh was used as the input *S*(*x, y, z*) to u-Unwrap3D to create a common static (*d, u, v*) coordinate space that the surface meshes from all timepoints were then mapped to in Step 5 of u-Unwrap3D to generate *S*(*d, u, v, t*) . We use Unwrap-3D with the following parameters for each step: Step 1, cMCF with maximum iterations = 50, *δ*_*t*_ = 1 × 10^−5^, automatic stopping threshold, Δ_*thresh*_ = 5 × 10^−5^, *S*_ref_(*x, y, z*) mesh voxelization with morphological dilation and erosion with ball kernel radius 5 voxels, Gaussian smoothing σ = 1 of the binary volume and initial marching cubes meshing at isovalue 0.5, ACVD remeshing with number of clusters = 10% the number of vertices in the marching cubes mesh, and further volume constrained Laplacian mesh smoothing^158^ with implicit time integration, time step 0.5 for 15 iterations; for Step 3 area distortion relaxation with maximum iterations = 100, *δ*_*t*_ = 0.1, stepsize *ε* = 1; for Step 4 we use the binary positive curvature region of *S*_ref_(*x, y, z*) given by 3-class Otsu thresholding as the weight for determining the unwrapping axis, and use a 512 x 1024 pixel (*u, v*) grid; for Step 5, for outwards propagation, an upsampling factor of 3 for binary voxelization, *α* = minimum of 0.5 and 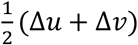 voxel -steps where Δ*u*, Δ*v* is the mean Cartesian 3D distance of traversing one pixel in *u, v* directions and a separable 1D uniform box filter smoother with a window 5 pixels; for inwards propagation, we use active contour cMCF with *δ*_*t*_ = 5 × 10^−4^, and robust mesh Laplacian^122^, mollify factor 1 × 10^−5^ for better numerical stability with the (*u, v*) parameterized *S*_ref_(*x, y, z*) converted into a triangle mesh by triangulating the quadrilateral pixel connectivity and ‘stitching’ the image boundaries into a spherical topology, and *D*_*in*_ = (maximum internal distance transform value) / *α* steps; for Step 6, marching cubes meshing of the topographic 3D mapped binary cell segmentation at isovalue 0.5 following Gaussian smoothing σ = 1, ACVD remesh with number of clusters = 50% the number of vertices in the marching cubes mesh and retaining the largest connected component mesh. Ruffles were height-projected, setting *S*(*d, u, v, t*) to *S*(*d* = 0, *u, v, t*) to map to the plane, and obtain *S*(*u, v*)(*t*).

#### Optical flow ROI tracking

We used motion sensing superpixels (MOSES)^98,99^ in dense tracking mode, which automatically monitors the spatial coverage of ROIs and introduces new ROIs dynamically to ensure uniform spatial tracking at every timepoint. We partitioned the image with an initial user-specified 1000 non-overlapping rectangular regions-of-interest (ROI). Each ROI was tracked over time by subsequently updating its centroid by the median optical flow^159^ within the ROI. Optical flow was computed from the Traction/CyOFP1 (TC) signal after rescaling TC(*S*(*u, v*)) to be an 8-bit grayscale image using the video minimum and maximum TC values.

#### ROI timeseries extraction and cross-correlation

Distortion-corrected average timeseries of mean Tractin/CyOFP1 (TC) and *H* were computed for a track using the same weighted mean as for blebs. A square bounding box of the mean MOSES ROI width centered at the track (*u, v*) coordinate was used to sample the scalar values at each timepoint. The distortion-corrected timeseries can be treated as a standard 1D timeseries. The 1D normalized cross-correlation was thus computed between the distortion-corrected TC and *H* timeseries for individual tracks without modification. ROI cross-correlation curves were averaged at all time-lags to derive the mean and 95% confidence interval ROI cross-correlation curve. A deviation of the curve greater than the 95% confidence interval at a time lag of 0 indicated significant instantaneous correlation.

#### Retrograde actin flow and mean ruffle travel speed

Computing the speed histogram with 25 bins and speed range 0-10 μm/min showed a slow and fast population (Fig. 5D). We inferred the mean speed of the two populations as the two thresholds generated by 3-class Otsu thresholding. The lower and faster of the thresholds are the mean speed of retrograde actin flow and ruffles respectively.

#### Cross-correlation and curvature relationship

We computed the continuous relationship of mean curvature, *H* and the lag 0 cross-correlation of TC and *H* (Fig. 5E) over ROI tracks using kernel density. Gaussian kernel density with a bandwidth set by Scott’s rule was used to derive the joint density distribution of *H* and cross-correlation i.e. *p*(*X, Y*), with *X*: *H, Y*: cross-correlation over the closed intervals *X* ∈ [−0.2, 0.6] and *Y* ∈ [−1, 1]. The continuous relationship is then given by the marginal expectation with capital denoting the random variable and 𝔼[⋅] the expectation operator, 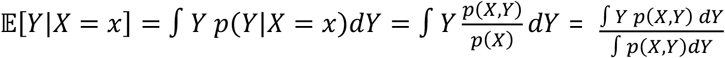 with standard deviation equivalently defined as the square root of the variance,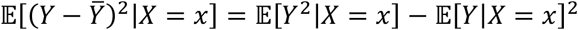. The evaluation of the integrals uses 100 bins for both *H* and cross-correlation.

### 3D timelapse lightsheet imaging and analysis of blebs

#### Cell culture and timelapse imaging

All details of the cell line creation, culture and imaging of the MV3 GFP-expressing melanoma cell movie in Fig. 7 were previously published^160^.^160^. The movie is a total of 200 frames acquired at a frequency of 1.21 s per frame. Each frame is a 104 x 512 x 512 size 3D volume with a voxel resolution of 0.300 x 0.104 x 0.104 µm.

#### Cell segmentation and surface meshing

The 200 timepoints were spatiotemporally registered volumetrically to the first timepoint, *t* = 0 as previously described^160^.^160^. The cell surface at *t* = 0 was segmented using a multi-level method^157^ involving local contrast enhancement, deconvolution and edge enhancement and surface meshed as described above to obtain *S*(*x, y, z, t* = 0). Images were deconvolved using the Wiener-Hunt deconvolution approach^161^ with our previously published point-spread function^39^.The surface mesh at all subsequent timepoints, *S*(*x, y, z, t*) were reconstructed using the non-rigid registration deformation field from volumetric registration^160^.^160^. The vertex Septin intensity was calculated by extending from the surface a trajectory to an absolute depth of 1 µm along the steepest gradient of the distance transform to the mesh surface and assigning the 95^th^ percentile of intensity sampled along that trajectory to the originating vertex to capture the systematically brightest accumulation of Septin signal in the cortical shell. The raw Septin intensity suffers decay from bleaching. We simultaneously normalized and corrected the vertex Septin intensity by computing a normalized Septin intensity as the raw intensity divided by the mean Septin intensity in the whole cell volume at each timepoint.

#### u-Unwrap3D analysis

We computed a mean surface mesh, 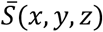 from all *S*(*x, y, z, t*) as the input surface to u-Unwrap3D. This was done by surface meshing the mean binary volume over all binary voxelizations of individual *S*(*x, y, z, t*) at an isovalue of 0.5. u-Unwrap3D was applied to 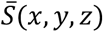 to create a common static (*d, u, v*) coordinate space that all *S*(*x, y, z, t*) is mapped into in Step 5 of u-Unwrap3D to generate *S*(*d, u, v, t*). u-Unwrap3D was run with the same parameters for all steps as for blebs in the validation dataset, except for the following modifications: step 1, the same automatic stopping iteration number but +5 steps, and ACVD with 10% of the marching cubes mesh to get a smoother *S*_ref_(*x, y, z*); step 4, a smaller 512 x 256 pixel size (*u, v*) grid and not optimizing the unwrapping axis - this axis passed through a large bleb and affected tracking; step 5, *D*_*in*_ = 96 steps - a total of 5µm. Topographic cMCF with the robust mesh Laplacian^122^, mollify factor = 1 × 10^−5^, *δ*_*t*_ = 5 × 10^4^ was applied to each *S*(*d, u, v*) for 10 iterations to compute the corresponding *S*_ref_(*u, v*)(*t*).

#### Bleb segmentation and tracking

Blebs were segmented from *S*(*d, u, v*) at every timepoint using the instance segmentation algorithm with refinement for undersegmented blebs as described above. In computing the binary protrusion segmentation we use a downsampling factor of 4 due to the smaller 512 x 256 pixel (*u, v*) grid and diffuse the segmentation for 5 iterations as the blebs were smaller than the validation dataset. The segmented 512 x 256 (*u, v*) bleb images, *F*_*bleb*_(*S*(*u, v*)) were padded 50 pixels on all four sides respecting spherical topology. This is done by periodic padding along the *u*-axis. For the *v*-axis, we pad the top of the image by reflecting the pixels with respect to the first image row (i.e. all pixels in row 2 to row 51) and then flipping in the *u*-axis. Similarly, the bottom is padded by refecting the pixels with respect to the last image row (i.e. all pixels in row 2 to row 511) and then flipping in the *u*-axis. For each unique bleb in every timepoint, we computed the bounding box of the bleb given by top left, (*u*_*min*_, *v*_*min*_) and bottom right (*u*_*max*_, *v*_*max*_) coordinates. The bleb bounding boxes were tracked using an optical flow assisted bounding box tracker^40^. Boxes were linked over time into tracks using bipartite matching and the intersection over union (IoU>0.25 for valid match) of bounding boxes as the distance function between pairs. To handle large changes in box size, the matching between the current and next frame was carried out on the predicted bounding box coordinates by local optical flow^159^.^159^. Optical flow was computed using the mean curvature, *H*(*S*(*u, v*)) after rescaling *H*(*S*(*u, v*)) to be an 8-bit grayscale image using the global minimum and maximum curvature values over time. In case of temporary occlusion or missed segmentation, any non-matched blebs were propagated for up to 5 frames (6s) using the estimated optical flow before track termination. Tracks with > 5 frames (6s) and a mean positive curvature, *H* > 0.1 µm^−1^ were retained as bleb tracks. The coordinates of retained tracks was corrected to account for the initial padding of 50 pixels. To remove erroneous and duplicated tracks, we uniquely match every segmented bleb in each timepoint to a track by IoU. For each track, we then computed the fraction of its lifetime that could be matched to a bleb and removed all tracks for which this proportion was < 50%. Lastly for each track we checked for sudden changes in the bounding box area, which was indicative of an erroneous bounding box in need of substitution by an inferred corrected bounding box. We applied this procedure to each track in order to construct the timeseries of the bounding box area over the track lifetime and compute a smooth reference timeseries using the central moving average with a window of 3 frames. The bounding box at a timepoint is erroneous if the instantaneous difference between the raw and smooth bounding box area > 500 pixel^2^ (the mean (*u, v*) bleb box area is 361 pixel^2^). The coordinates of a corrected bounding box is inferred from non-erroneous bounding boxes by interpolation using a linear spline. The tracks that remained fully in-focus over its lifetime were retained for analysis.

#### Bleb timeseries extraction

We detected (*u, v*) pixels on blebs by labelling spatially contiguous areas of positive mean curvature based on 3-class Otsu thresholding defining positive, flat or negative curvature. The largest connected component within a bleb bounding box was defined as on-bleb and the remainder area within the bounding box as off-bleb. We extracted distortion-corrected average timeseries of bleb area, mean curvature and septin intensity, that is of a scalar quantity, *F* by observing that the mean of *F* over a Cartesian 3D surface area is equivalent to computing a weighted mean over the equivalent (*u, v*) area,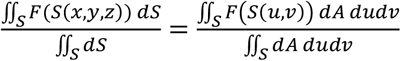. The weight, *dA* is the magnitude of the differential area element, 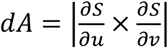 described above.

#### Bleb event alignment

Individual blebbing events were detected within a track by applying peak finding after central moving averaging of bleb area timeseries with a window of 3 timepoints. A peak was defined as having a prominence > 0.5 and separated from a neighboring peak by at least 3 timepoints. Individual bleb event timeseries were constructed and temporally aligned using the detected timepoint of maximal bleb area as timepoint 0 and taking a window of 14 timepoints on either side (a total 29 timepoints, 35 s).

**Figure S1.**
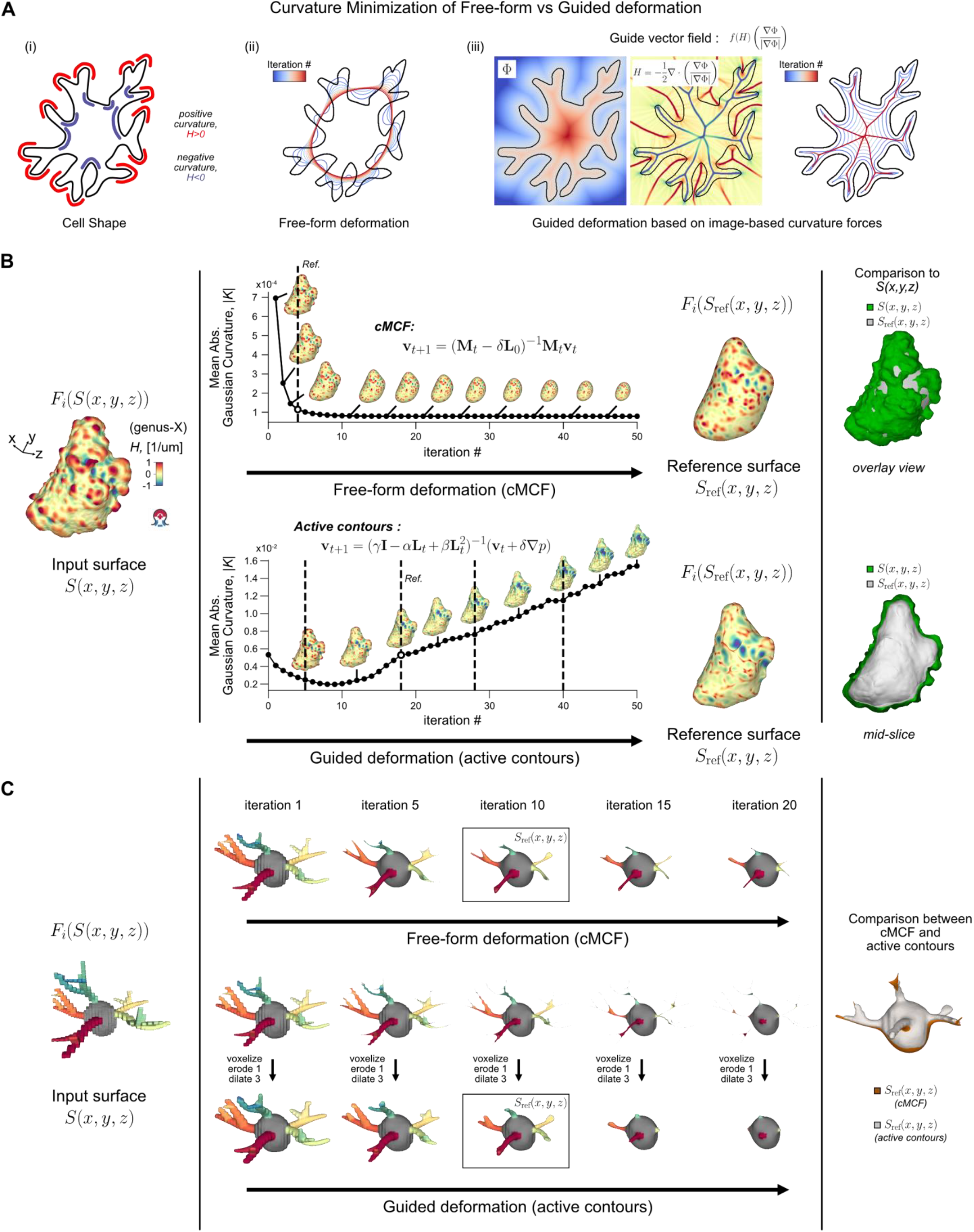
Free-form vs guided curvature minimizing flow for determining reference shape from input surface mesh. **A)** Illustration of the same i) 2D cell contour, under ii) free-form deformation whereby positive curvature segments contracts inwards and negative curvature segments expands outwards; and under the guidance of the iii) mean curvature of the Euclidean distance transform gradient whereby the contours always contracts inwards converging towards the ridges of the distance transform. Crucially, the guided always evolves within the area of the original shape. **B)** Iterative deformation of an input surface with blebs (left) and resulting automatic inferred reference shapes by changepoint detection (Methods) under free-form cMCF (upper) or guided active contours (lower) that selectively erodes only positive mean curvature surface features (middle). Comparison to original surface by overlay for cMCF to show this reference is not confined by volume (upper) and by cutout for active contours to show this reference always lies within the original (lower). **v** denotes mesh vertex coordinates, **M** is the mesh mass matrix, **L**_**0**_ is the mesh Laplacian matrix at iteration number *t* = 0, *γ* = 1 is a regularization parameter and *α, β* are continuity and smoothness control parameters in active contours, ∇*p* is the guiding gradient flow, here the mean curvature weighted Euclidean distance transform gradients in A) iii). **C)** Iterative deformation of a multi-branched cell surface (left) under cMCF and the same guided active contours in B) (middle). Overlay of the reference surface at iteration 10 between the two deformations. cMCF branch centerlines align with that of active contours and therefore by extension of *S*(*x, y, z*) branches. However, active contours leads to branch thinning and neck pinching singularities. To be suitable for equiareal spherical parameterization, reference shapes need to be voxelized, eroding overly thin regions and dilated to have sufficient thickness (Methods).

**Figure S2.**
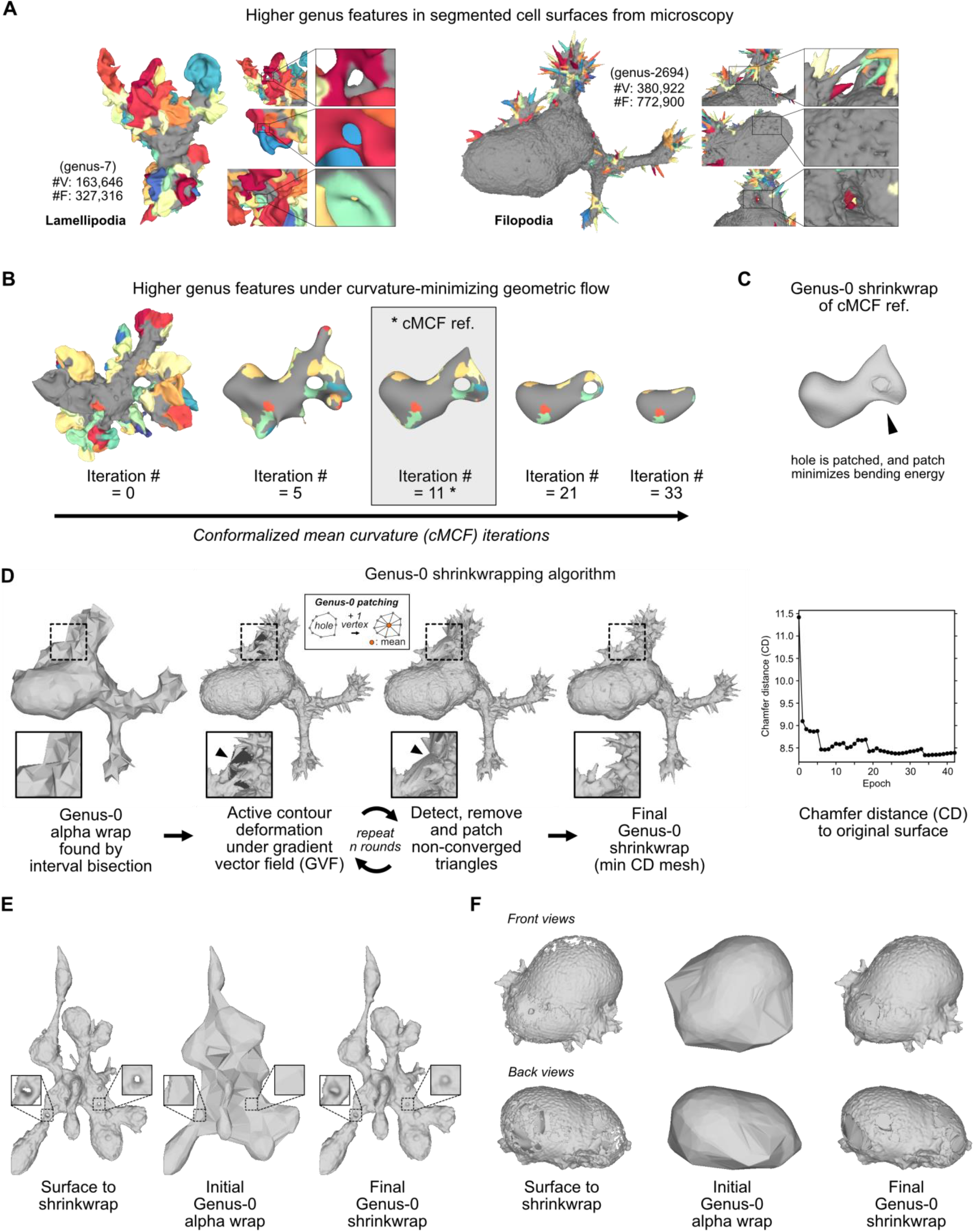
Guaranteed genus-0 shrink-wrap of surfaces by u-Unwrap3D. **A)** Typical examples of higher genus inducing ‘hole’ and ‘handle’ errors in 3D meshes obtained by marching cubes meshing of binary segmentations from volumetric lightsheet microscopy images for two morphological motifs and two cell types; a dendritic cell with lamellipodia (left) and a HBEC cell with filopodia (right). **B)** Example of large handles too large to be closed by simple voxelization and morphological operations to make the reference shape computed by conformalized mean curvature flow (cMCF) genus-0. **C)** genus-0 equivalent of cMCF reference mesh in B) after u-Unwrap3D genus-0 shrink-wrapping. **D)** Overview of u-Unwrap3D’s genus-0 shrink-wrapping algorithm (Methods) which initializes an initial genus-0, watertight, intersection-free, 2-manifold mesh using 3D alpha wrapping^66^. The initial mesh is iteratively deformed according to the target shape’s gradient vector field^67^, whilst preserving the genus to produce an optimized shrink-wrap (left). Chamfer distance between shrink-wrap and target shape per epoch (right). **E)** Shrink-wrapping of a spheroid with significant heterogenous branching and a narrow gap between branches. **F)** Shrink-wrapping of a partially segmented cell with large holes. The final shrink-wrap produces a minimum bending energy imputation of holes whilst simultaneously preserving the intricacies of the ruffles. See Movie S2 for iterative shrink-wrapping of the surfaces in panels D-F.

**Figure S3.**
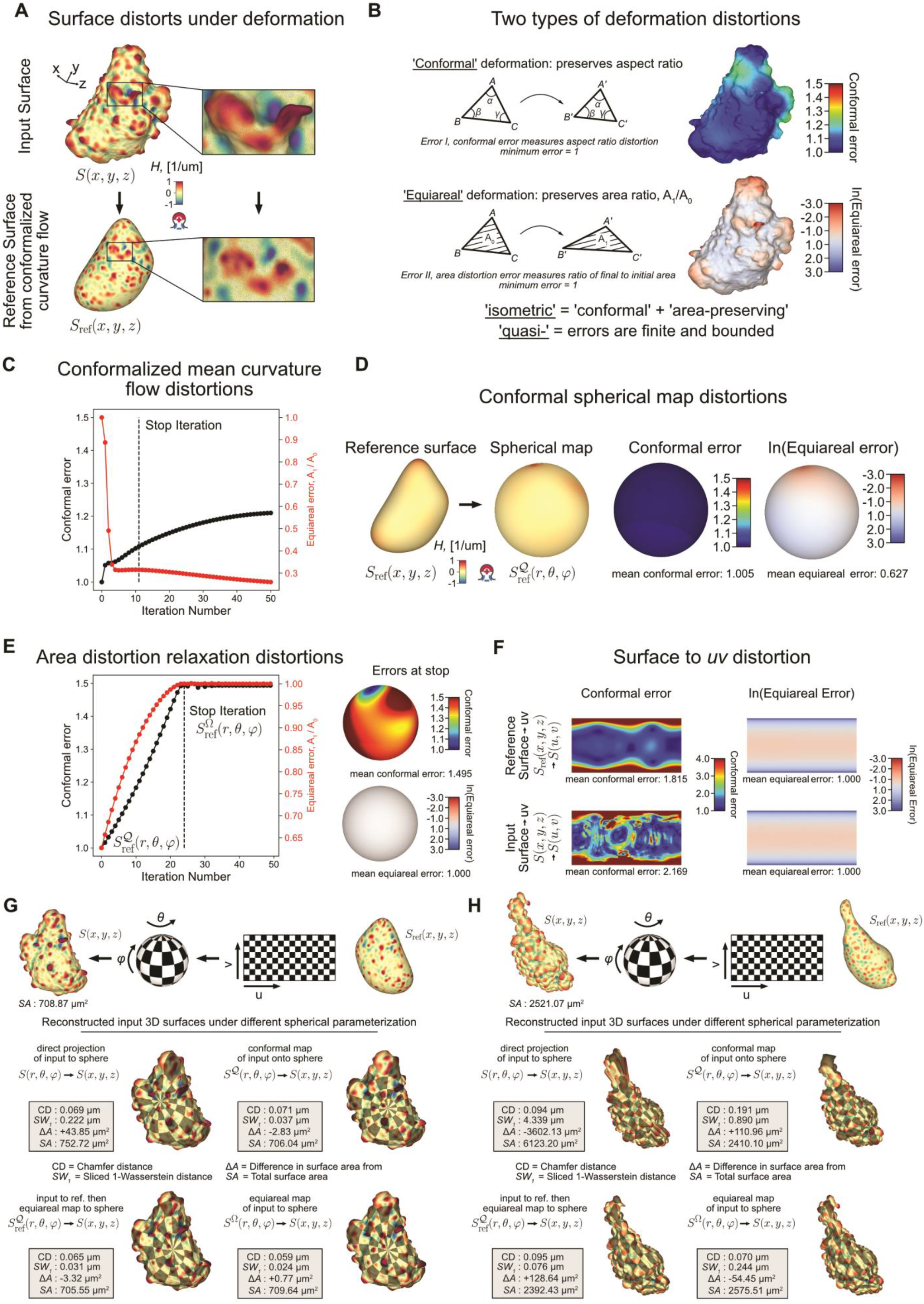
Measuring and optimizing mesh distortion under surface deformation. **A)** Any deformation of a closed 3D surface mesh (top) such as by conformalized mean curvature flow (cMCF) (bottom) distorts local geometrical distances and areas as illustrated by zoom-ins. **B)** Illustration of the two types of metric distortion incurred by deformation; conformal and equiareal. In general, lower conformal error is at the expense of equiareal preservation and vice versa. **C)** Plot of the conformal error (black dotted line, black left y-axis) and equiareal error (red dotted line, red right y-axis) for each iteration of cMCF (Step 1, Fig. 1B) for the example mesh in A) and Fig. 1 with stop iteration indicated by a black vertical dashed line. **D)** Rendering of the conformal and equiareal error at each triangle face for quasi-conformal spherical parametrization of the smooth shape to the sphere (Step 2, Fig. 1B). **E)** Plot of the conformal error (black dotted line, black left y-axis) and equiareal error (red dotted line, red right y-axis) for each iteration of the spherical area distortion relaxation with stop iteration indicated by a black vertical dashed line (Step 3, Fig. 1B), (left). Rendering of the conformal and equiareal error of individual triangle faces at the stop iteration, (right). **F)** Comparison of the per pixel conformal error (left column) and equiareal error (right column) of the 2D (*u, v*) unwrapping of the cMCF smooth shape, *S*_ref_(*x, y, z*) (upper row) or direct unwrapping of the input shape, *S*(*x, y, z*) (lower row). **G)** Quantitative assessment of four different options of 2D (*u, v*) unwrapping of an input surface, *S*(*x, y, z*) with blebs via different spherical parameterizations, *S*(*r, θ, φ*) by measuring the difference between the Cartesian 3D reconstructed mesh from *S*(*u, v*) and *S*(*x, y, z*). *CD* = Chamfer distance, *SW*_1_ = sliced 1-Wasserstein distance between vertices of the input and reconstructed mesh. *SA* = the total surface. Δ*A* = difference in total surface area between the input and reconstructed mesh. Qualitative assessment by uv-remapping the chessboard pattern and blending with the mean curvature, *H* colors. **H)** Same as G) for an elongated cell with blebs.

**Figure S4.**
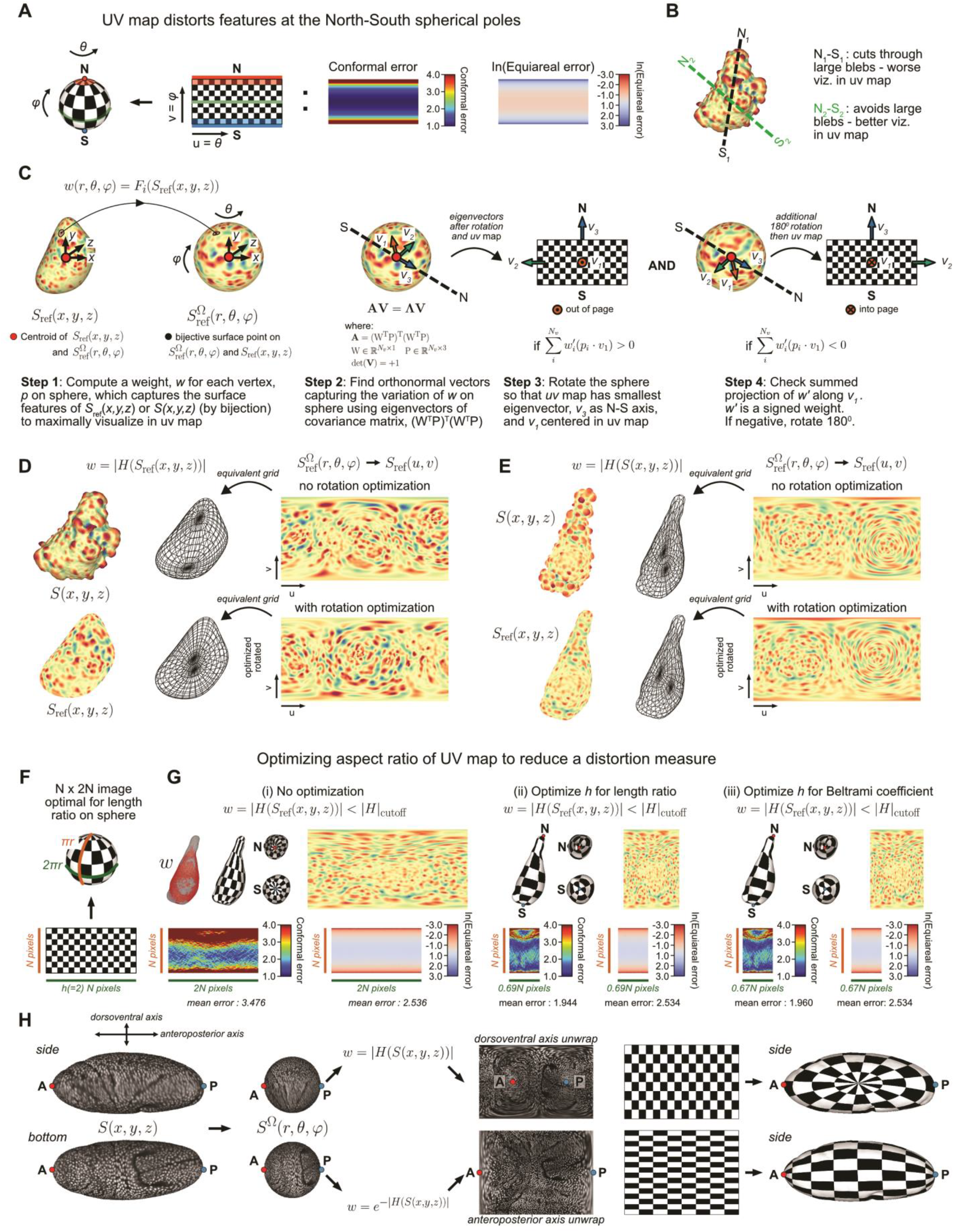
Automatic determination of *(u,v)* unwrapping axis based on vertex weight function and optimization of (u,v) parameterization aspect ratio. **A)** Illustration of the increase in conformal and area distortion towards the north and south poles of the sphere. **B)** Illustration of how choice of unwrapping axis impacts the representation of surface features-of-interest, here bleb size, due to the increased distortion at the poles in the *(u,v)* parameterization. **C)** Schematic of the automatic unwrapping axis determination based on eigendecomposition from the spherical parameterization of the reference surface with a given vertex weight function *w* (Methods). *w* should be specified to highly score features-of-interest that should be mapped to the equatorial band of the sphere. **D)** *(u,v)* parameterization of a cell with blebs with and without optimizing the unwrapping axis. *w* is defined as the absolute mean curvature of *S*_ref_(*x, y, z*) . **E)** *(u,v)* parameterization of an elongated cell with blebs with and without optimizing the unwrapping axis. *w* is defined as the absolute mean curvature of *S*(*x, y, z*) . **F)** Schematic of the default 2:1 ratio of (u,v) parameterization based on the ratio of the great arc lengths of the sphere. **G)** *S*(*x, y, z*) mean curvature, conformal and equiareal distortion with (i) no optimization, and optimization of the width of the *(u,v)* parameterization by (ii) the ratio between the mean Cartesian length traversed along *u* coordinates vs that traversed along *v* coordinates; and (iii) by minimizing for the Beltrami coefficient. For all cases, *w* is defined as a binary to target flatter surface areas to the equatorial band. **H)** *(u,v)* parametrization of an ellipsoidal drosophila embryo along dorsoventral axis (top) or anteroposterior axis (bottom) using different *w* derived from mean curvature, without needing to explicitly specify the unwrapping axis or pre-rotating the embryo.

**Figure S5.**
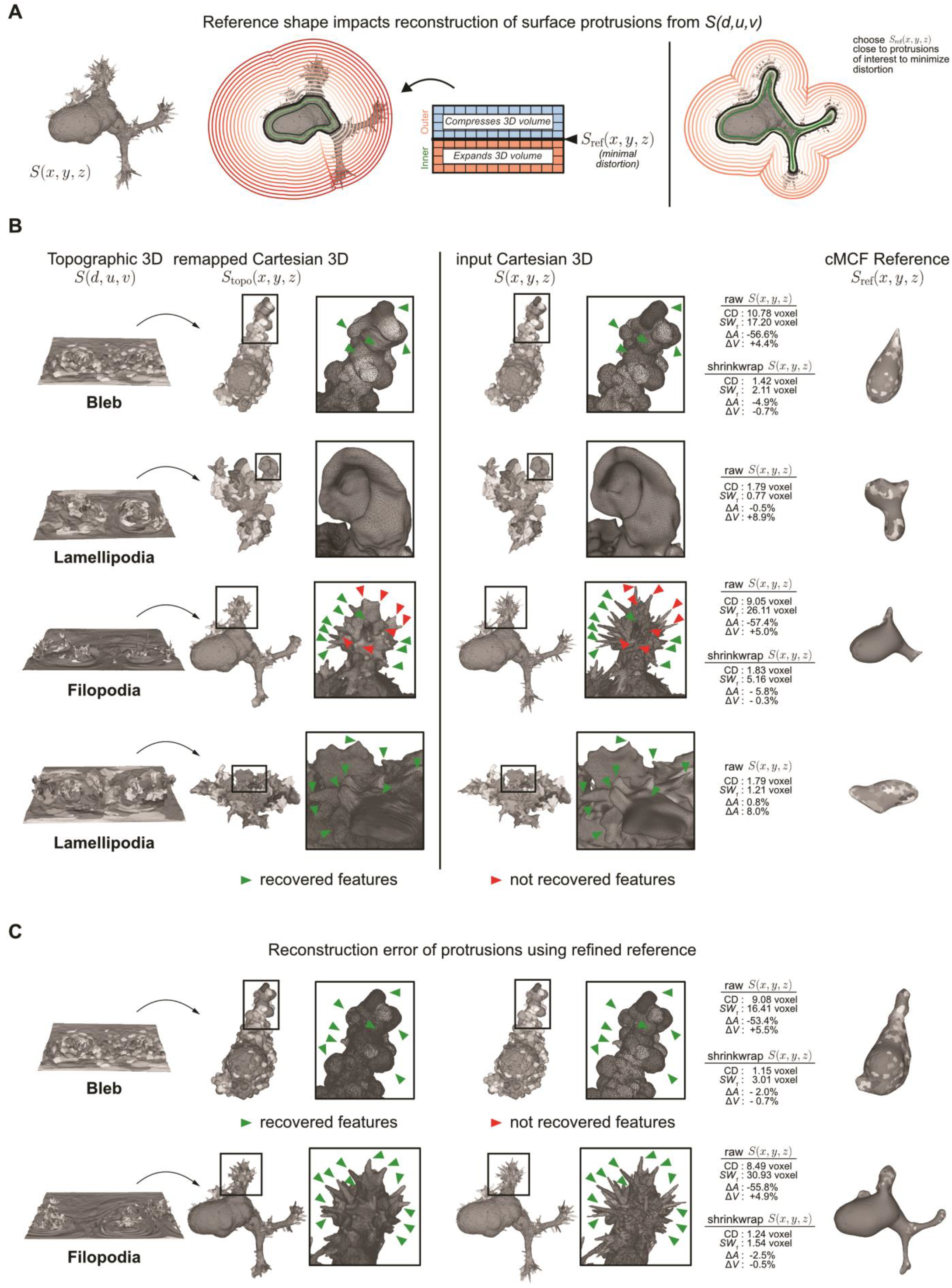
Choice of reference shape affects reconstruction accuracy of protrusions from the *(d,u,v)* topographic representation to Cartesian 3D. **A)** Illustration of the metric distortion of the Cartesian volume proximal to the reference surface when mapped into topography space (left). The space external to *S*_ref_(*x, y, z*) is contracted whilst the internal space is expanded. The degree of distortion increases with distance to *S*_ref_(*x, y, z*) . Consequently, a *S*_ref_(*x, y, z*) closest to the protrusions-of-interest maximizes the resolution in topographic representation (right). **B)** Quantitative and qualitative comparison of the Cartesian 3D remapping, *S*_topo_(*x, y, z*) of the topographic mesh, *S*(*d, u, v*) (left) and the original input mesh *S*(*x, y, z*) (middle) for 4 cell examples with different morphological motifs in relation to their chosen reference surface, *S*_ref_(*x, y, z*) (right). Box shows a zoom-in of the local surface region for each example. Green triangles highlight exemplar protrusion morphologies that are well reconstructed from *S*_topo_(*x, y, z*) in Cartesian 3D. Red triangles highlight exemplar protrusion morphologies, particularly filopodia which are poorly reconstructed from *S*_topo_(*x, y, z*), due to being distant from *S*_ref_(*x, y, z*). Gray shading of the surface mesh visualizes individual protrusion morphologies. **C)** Quantitative and qualitative comparison of the bleb and filopodia exemplars in B) with topographic mesh, *S*(*d, u, v*) computed using a reference shape closer to the original *S*(*x, y, z*).

**Figure S6.**
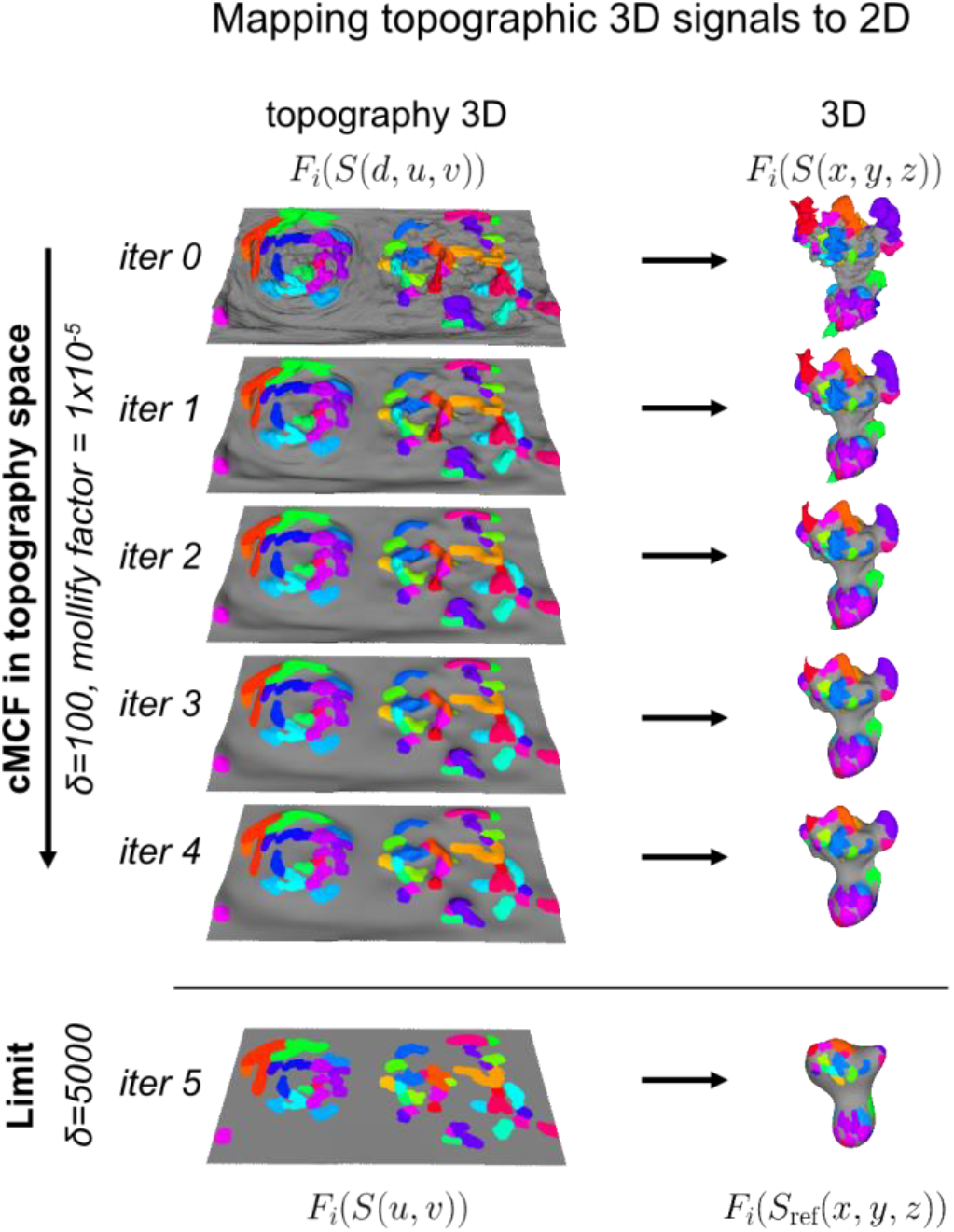
Topographic conformalized mean curvature flow to flatten topographic signals into 2D for visualization and analysis. Illustration of the topographic conformalized mean curvature flow (cMCF) (Methods) using the robust Laplacian matrix for nonmanifold triangle meshes^122^ to directly map topographic 3D surfaces and associated signals (here, segmented protrusions marked by unique colors) to the 2D plane, an optimal representation for tracking individual protrusions, and correcting protrusion segmentations performed in the topography representation to take into account the spherical boundary conditions. A few iterations is sufficient for convergence. The robust Laplacian matrix is defined with an additional parameter, the mollify factor, which controls numerical stability. The larger the mollify factor, the greater the stability. Otherwise, the robust Laplacian drop-in substitutes for the standard cotangent Laplacian matrix without modifying the cMCF update equation (Methods). As with the standard Cartesian 3D cMCF, increasing *δ*, increases the time step, and increases convergence speed.

**Figure S7.**
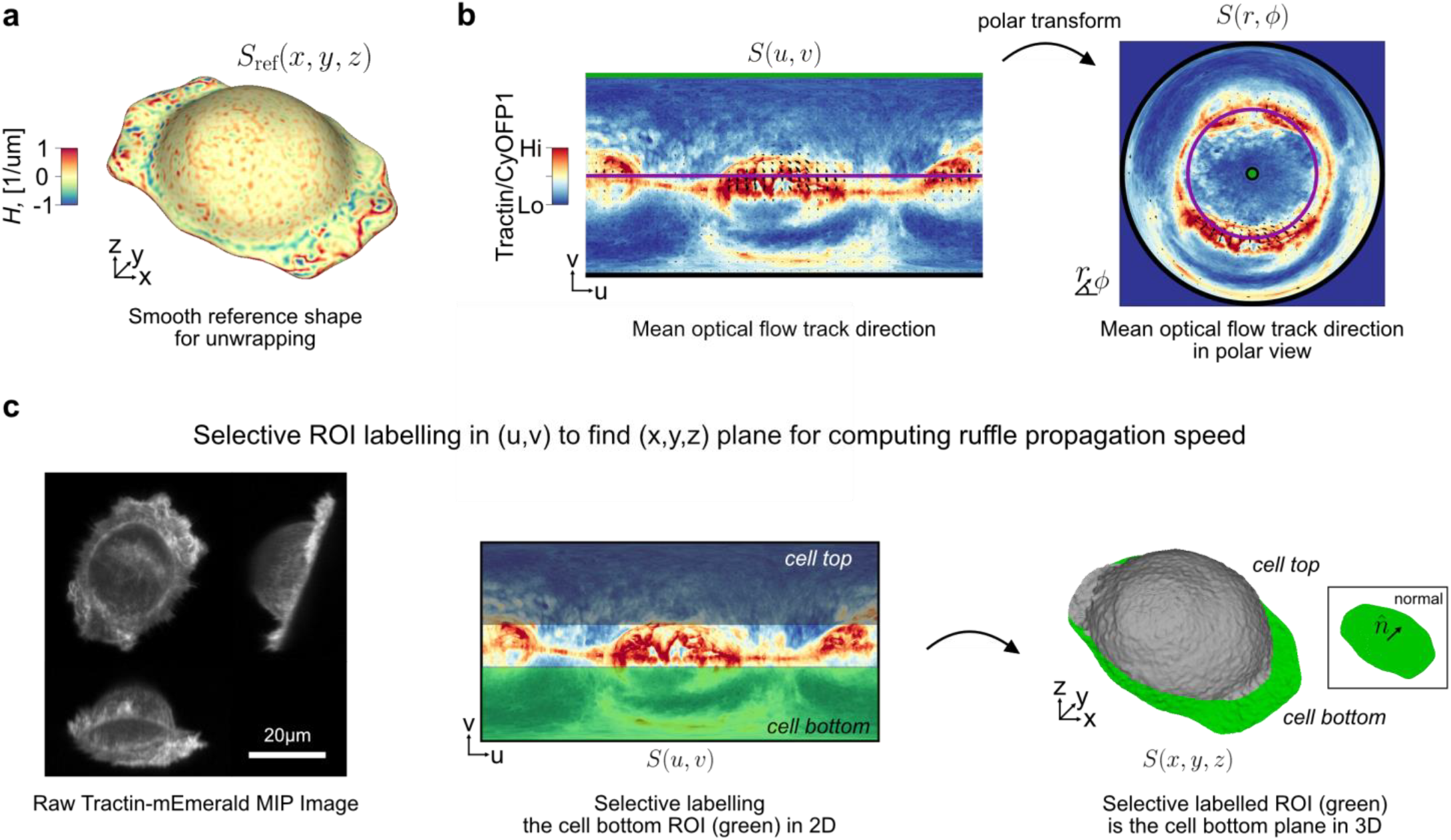
u-Unwrap3D enabled measurement of ruffle in-plane travel speed. **A)** Cortical reference shape, *S*_ref_(*x, y, z*) found by applying conformalized mean curvature flow to the first timepoint of the ruffling SU.86.86 cell in Fig. 5A and used to unwrap all timepoints into a common temporally-invariant (*d, u, v*) coordinate space. **B)** The mean track velocity of the optical flow region-of-interest (ROI) tracks, plotted as black arrows at the initial track coordinate overlaid on the unwrapped Tractin-mEmerald/CyOFP1 intensity image of Fig. 5B (left) and its corresponding polar transformed equivalent (right). The polar transform maps the green (top), purple (middle) and black (bottom) horizontal line on the left to the central green point, second purple and third black rings in the polar image. **C)** Orthogonal cross-section maximum intensity projection (MIP) image of the raw Tractin-mEmerald intensity channel showing the tilted lightsheet acquisition (left). Selective ROI isolation of the top and bottom surface of the cell by grey and green bounding box selection in unwrapped view (middle) and visualized in 3D with grey and green surfaces, respectively (right). The normal vector, 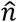 describing the best fit plane through only the cell bottom vertices found by principal component analysis (inset rectangle).

## Supplementary Movies

**Movie S1**. Overview of the key steps of u-Unwrap3D and illustration of direct unwrapping using the input cell surface as reference in conjunction with the spatial activation pattern of PI3K signaling products. Application of u-Unwrap3D to an MV3 melanoma cell with bleb surface motifs, dendritic cell with lamellipodia surface motifs, and HBEC cell with filopodia surface motifs segmented by the u-shape3D software. Each motif is labelled with a random color to demonstrate the local surface mappings and their distortions in different representations. All reference shapes were determined by conformalized mean curvature flow by threshold on the absolute Gaussian curvature change. Additional application to map movies of T cells with microvilli, FiloGen simulated cells developing branching, and the surface of a complex cell aggregate with a diverse mixture of motifs.

**Movie S2**. Application of u-Unwrap3D’s genus-0 shrink-wrapping to tightly wrap complex surfaces including those with extensive concavities, and narrow necks, even incomplete surfaces with numerous large holes due to imperfect cell segmentation. The examples underscore the need for guided mesh deformation to guarantee tight wrapping following an initial genus-0 alphawrap. However, if the initial genus-0 alphawrap is already optimal, mesh deformation does not further improve the wrapping. If performed, the minimum chamfer distance shrink-wrap to the input will still return the initial genus-0 alphawrap.

**Movie S3**. Application of topographic conformalized mean curvature flow to directly map the topographic surface *S*(*d, u, v*) of a dendritic cell with segmented lamellipodia surface motifs to the 2D plane, *S*_*ref*_(*u, v*) for two different time steps, *δ* = 100 and *δ* = 5000. The smaller *δ* enables gradual relaxation and the ability to sample and use intermediate shapes during the flow. However, smooth low curvature folds remain such that we do not fully converge to the plane even if continued to 100 iterations, though this will still be diffeomorphic to the 2D plane. As the flow is stable, generally, for direct mapping we will always use a large *δ* to ensure convergence within 50 iterations.

**Movie S4**. Application of u-Unwrap3D to each timepoint to analyze examples of timelapse data. Mapping of *S*_*ref*_(*x, y, z, t*) to the static geometries of the 3D sphere or 2D plane automatically establishes spatiotemporal correspondence across all timepoints. Moreover, the introduction of cortical *S*_*ref*_(*x, y, z, t*) isolates surface protrusions in the topographic representation, *S*(*d, u, v, t*) achieving a natural decomposition of the cell surface. This is demonstrated by transferring *S*(*x, y, z, t*) blebs to decorate a spherical cortical shape.

**Movie S5**. Application of u-Unwrap3D to interrogate surface protrusion dynamics within the developing immunological synapse between a natural killer cell and a leukaemic cancer cell. TV-L1 optical flow tracking of protrusions in 2D is compared to the multiscale information flow behaviour computed by u-InfoTrace. Application of u-Unwrap3D to investigate cytokeratin 14 expressing leader cell behavior at and around the leading edge of developing invasive strands in breast organoids over time.

**Movie S6**. Application of u-Unwrap3D to track the surface ruffling and actin flows of a SU.86.86 pancreatic adenocarcinoma. View 1: Projections of cell surface, mean curvature and Tractin-mEmerald into topographic surface and (*u, v*) unwrapped reference surface representations. View 2: Regional ruffling and actin flows are tracked in 2D with optical flow and trajectories projected into a 2D polar representation and back to the original 3D surface. View 3: Select measurement of instantaneous ruffle and actin flow speeds and cross-correlation of actin and curvature within the lamella and lamellipodia taking advantage of the unwrapped (*u, v*) representation is projected back to the original 3D surface. The timelapse volumes were acquired every 2.27s for 30 frames. Scalebar: 20μm.

**Movie S7**. Application of u-Unwrap3D to enable segmentation and tracking of blebs on a MV3 melanoma cell in topographic representation. View 1: Projections of cell surface, mean curvature and normalized SEPT6-GFP into topographic surface and (*u, v*) unwrapped reference surface representations. Individual blebs are segmented by thresholding in the topographic surface representation. Leveraging the bidirectionality of u-Unwrap3D mappings bleb labels are projected back to the 3D surface. View 2: The 2D segmented blebs are tracked and trajectories projected back to the original 3D surface. The timelapse volumes were acquired every 1.21s for 200 frames. Scalebar: 10μm.

